# Chemical History Shapes the Antigenic Identity of MHC-I Epitopes

**DOI:** 10.1101/2025.11.06.687013

**Authors:** Sarah E. Newkirk, Joey J. Kelly, Karl Ocius, Carson Starnes, Sameek Singh, Tian Zhang, Marcos M. Pires

**Affiliations:** Department of Chemistry, University of Virginia, Charlottesville, VA 22904, United States; Department of Biochemistry and Molecular Genetics, School of Medicine, University of Virginia, Charlottesville, VA 22904, United States; Department of Microbiology, Immunology, and Cancer, University of Virginia, Charlottesville, VA 22904, United States

## Abstract

Cytotoxic T lymphocytes recognize infected and transformed cells through peptide antigens presented by MHC-I. Peptide sequence is assumed to dictate recognition, yet non-enzymatic post-translational modifications (PTMs) installed by reactive electrophiles can render a peptide chemically distinct from the form against which tolerance was established. Here we show that non-enzymatic PTMs arising from oxidative, inflammatory, metabolic, and carbonyl stress alter peptide-MHC-I stability and frequently disrupt recognition by a cognate TCR despite preserved MHC-I binding. Using a thioester probe, we demonstrate proof-of-concept capture of acylation-susceptible MHC-I ligands from the cellular immunopeptidome. Environmental chemicals, pharmaceuticals, and dietary isothiocyanates likewise modify antigenic peptides and impair T cell activation, and chemically distinct adducts at a single residue span the range from complete loss to full retention. Finally, methylglyoxal glycation impairs antigen-specific activation, and scavenging by creatine prevents it. Antigenic identity therefore reflects not only sequence but a peptide’s cumulative chemical history.

## INTRODUCTION

The adaptive immune system detects infected and transformed cells through peptide antigens displayed by major histocompatibility complex (MHC) molecules.^1, 2^ MHC class I (MHC-I) molecules are expressed on nearly all nucleated cells and present peptides drawn from intracellular pools that report the proteomic state of the cell. Cytotoxic T cells survey these complexes and eliminate cells displaying peptides not recognized as ‘self’. This discrimination is established during central tolerance: negative selection in the thymus eliminates T cells whose receptors (TCRs) bind self-peptide-MHC complexes too strongly, shaping the circulating repertoire.^3^ Central tolerance can only cover what the thymus presents. A chemical modification that is rare or absent in the thymic peptide pool is therefore poorly represented during negative selection, and this gap widens with age as thymic output declines. Peptides that emerge or become chemically modified later in life can consequently fall outside the immune system’s ‘self’ registry, although peripheral mechanisms including anergy and regulatory T cell suppression provide additional checkpoints that we do not address here.

Post-translational modification (PTM) reshapes the proteome through both regulated enzymatic pathways and non-enzymatic chemistry. Canonical enzymatic PTMs, including phosphorylation, glycosylation, and methylation, can markedly alter MHC-I binding affinity and TCR recognition once the modified peptides are processed and presented,^4, 5^ an effect we recently surveyed systematically.^6^ Critically, such modifications remain tightly regulated: typically reversible and cleared by dedicated ‘eraser’ enzymes (**Fig. 1a**).^7^ Non-enzymatic modifications are installed stochastically by reactive electrophiles from endogenous metabolism and environmental exposure, and most lack a corresponding eraser. Dedicated repair systems exist for a minority of marks, including methionine sulfoxide reductases, sulfiredoxin, and the glyoxalase pathway, but the majority of adducts persist once formed and are therefore long-lived at the cell surface.^8^ Modifications that are poorly represented in the thymus, including those that accumulate after age-related involution, may therefore be incompletely covered by central tolerance.

**Figure 1.**
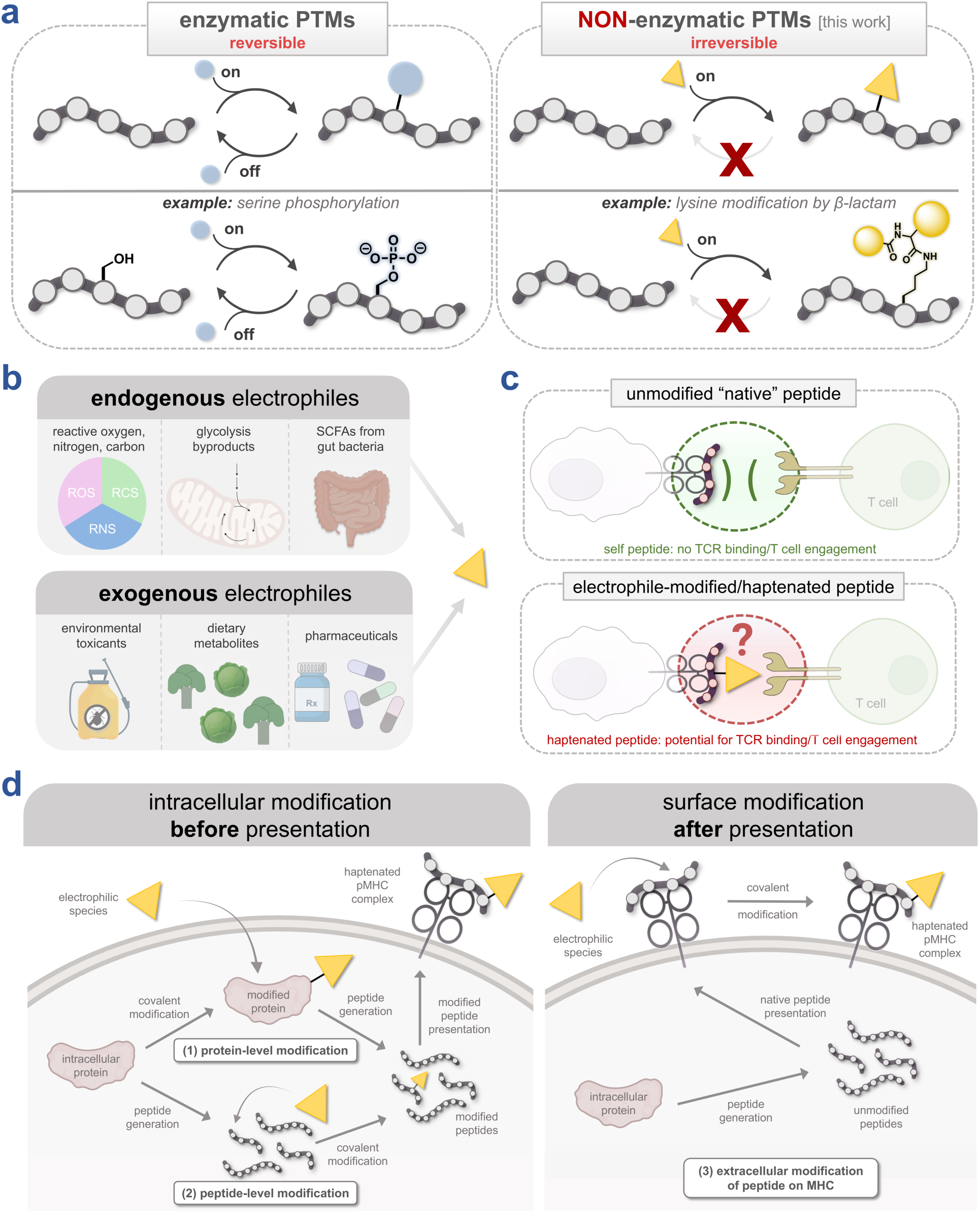
**(a)** Representation of the reversibility of most canonical enzymatic PTMs (left) relative to the typically irreversible non-enzymatic PTMs (right). **(b)** Schematic illustrating the sources of endogenous and exogenous electrophiles capable of modifying nucleophilic amino acid residues. **(c)** Representation of how electrophilic chemical species may alter immune response through modification of an MHC-I presented peptide. **(d)** Schematic depiction of potential pathways of electrophilic modification, including intracellular modification of proteins or peptides prior to MHC-I presentation, and modification of MHC-I-presented peptides on the cell surface.

The electrophiles that install these modifications are chemically diverse but converge on a single mode of action, covalent adduct formation at nucleophilic side chains (**Fig. 1b,c**). Endogenous sources include reactive oxygen and nitrogen species and reactive carbonyl species such as methylglyoxal (MGO), together with short-chain fatty acids, which are produced by the gut microbiota from dietary fiber and drive widespread lysine acylation once activated to reactive acyl-CoA intermediates.^9^ Exogenous sources include environmental pesticides,^10^ pharmaceuticals,^11^ and dietary metabolites.^12^ Several widely used pesticides are potent electrophiles with reactivities comparable to those of covalent drugs. Urushiol, the primary immunogen of poison ivy, is a pro-hapten that after oxidation to its ortho-quinone covalently modifies skin proteins, generating haptenated peptides that trigger T cell-mediated allergic responses.^13^

Here, we investigate how non-enzymatic modifications, installed by both endogenous and exogenous electrophiles, reshape MHC-I antigen presentation and T cell recognition (**Fig. 1d**). Using defined synthetic and cancer-derived epitopes, we show that individual modifications can either stabilize or destabilize the pMHC-I complex and, independently, can abrogate recognition by a defined TCR even when MHC binding is preserved. We then develop a chemical-probe enrichment strategy that captures modified ligands directly from the cellular immunopeptidome. Finally, we define the chemistries that install these modifications, spanning endogenous metabolites, environmental chemicals, and dietary isothiocyanates (ITCs), and identify two strategies that prevent them. One principle links these threads: the antigenic identity of a peptide reflects not only its sequence but its cumulative chemical history. This broadens the immunological definition of ‘non-self’ to include non-enzymatically modified peptides of either endogenous or exogenous origin.

## RESULTS

### Non-enzymatic PTMs alter pMHC-I stability and T cell activation

To examine how non-enzymatic PTMs affect MHC-I display, we used the canonical ovalbumin epitope SIINFEKL (**ovaWT**), whose interactions with H-2K^b^ and cognate TCRs are well defined,^14^ as a model scaffold (**Fig. 2a**). We measured pMHC-I stabilization with TAP-deficient RMA-S cells, in which impaired endogenous loading leaves MHC-I molecules empty and thermally unstable, so that surface H-2K^b^ abundance reports on the stability of the complex formed with an exogenously supplied peptide.^15, 16^ Throughout, we refer to this readout as stabilization rather than affinity, since it is dominated by complex half-life rather than by an equilibrium constant. As a negative control, we used SNFVSAGI (**cntPEP**), which does not bind H-2K^b^.^17^ T cell recognition was assessed by co-culture with B3Z cells, which express a SIINFEKL-specific TCR and an NFAT-LacZ reporter.^18^

**Figure 2.**
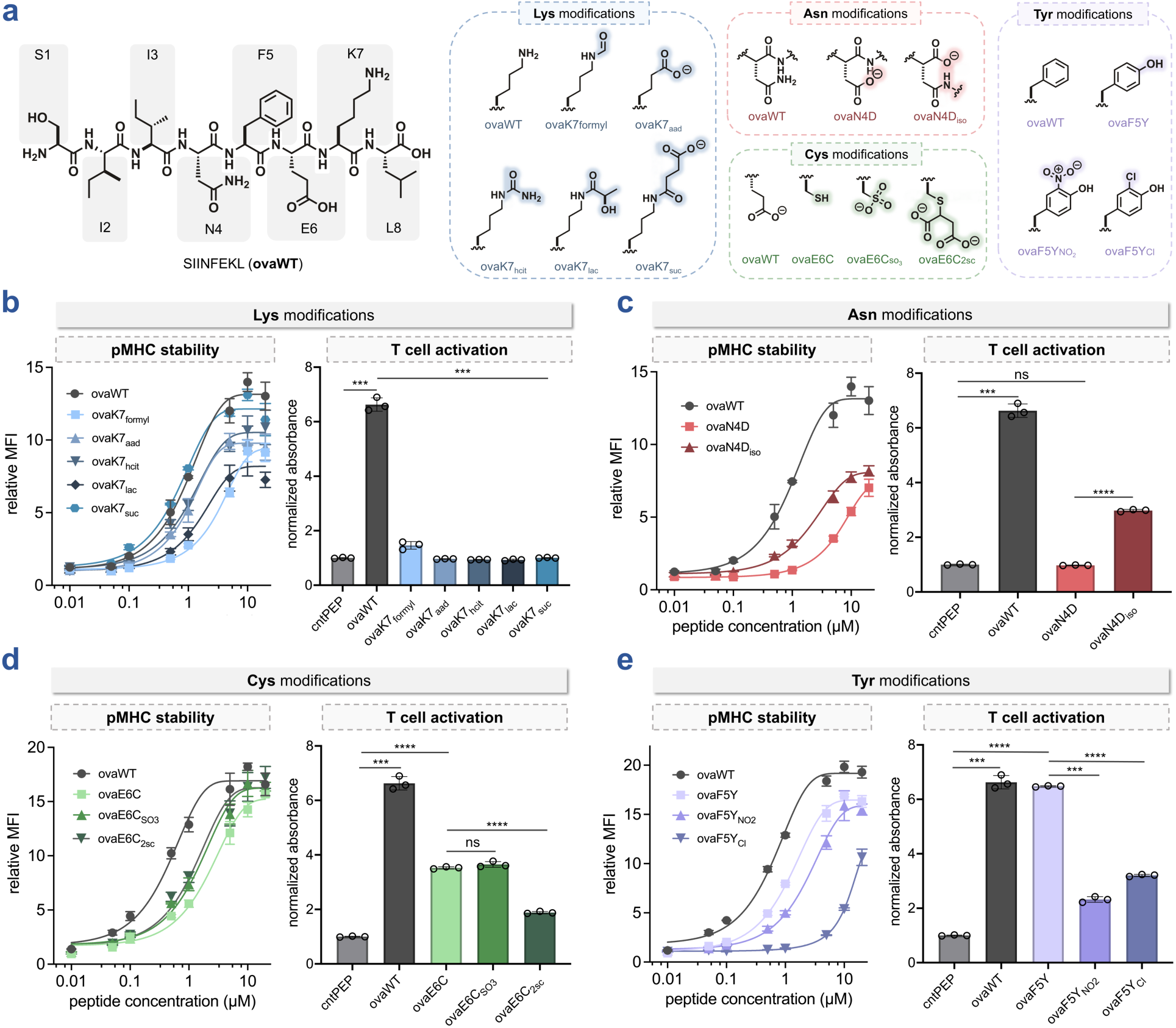
**(a)** Chemical structure of **ovaWT** and associated non-enzymatic PTMs to lysine (blue), asparagine (red), cysteine (green), and tyrosine (purple). **(b – e)** Dose-response curves from flow cytometry analysis of the RMA-S stabilization assay, and B3Z T cell activation assay for non-enzymatic PTMs to **(b)** lysine, **(c)** asparagine, **(d)** cysteine, and **(e)** tyrosine residues of SIINFEKL. For RMA-S stabilization assay, RMA-S cells were incubated with the indicated concentration of peptides and analyzed via flow cytometry for H-2K^b^ expression by APC anti-mouse H-2K^b^ antibody. MFI is the mean fluorescence intensity of the level of fluorescence relative to the negative control peptide (**cntPEP**). Data are represented as mean ± SD (*n* = 3) of technical replicates, and Boltzmann sigmoidal curves were fitted to the data using GraphPad Prism. For B3Z T cell activation assay, RMA-S cells were incubated with 5 μM of indicated peptides for 1 h at 26°C. RMA-S cells were subsequently co-incubated with B3Z T cells for 6 h at 37°C. β-galactosidase expression was then measured via the conversion of the colorimetric reagent chlorophenol red-β-D-galactopyranoside (CPRG) on a plate reader at 570 nm. The data presented have been normalized to the absorbance of the negative control peptide (**cntPEP**). Data are represented as mean ± SD (*n* = 3) of technical replicates. *P*-values were determined by a two-tailed *t*-test (ns = not significant, *** *p* < 0.001, **** *p* < 0.0001).

We began at position 7, which carries a lysine, one of the most frequently modified residues in the proteome. We synthesized variants representing marks associated with oxidative, inflammatory, and metabolic stress: formylated lysine (**ovaK7_formyl_**), aminoadipic acid (**ovaK7_aad_**), homocitrulline (**ovaK7_hcit_**), D-lactylated lysine (**ovaK7_lac_**), and succinylated lysine (**ovaK7_suc_**) (**Fig. 2a**). These span distinct formation routes: direct oxidative damage to lysine,^19^ reaction with electrophiles generated during oxidative processes,^20^ carbamylation by isocyanic acid,^21^ non-enzymatic acyl transfer from succinyl-CoA,^22^ and glycolysis-derived intermediates such as MGO.^23, 24^ All five variants stabilized H-2K^b^, consistent with this residue being solvent-exposed and making limited contact with the MHC (**Fig. 2b**).^25^ **ovaK7_formyl_** and **ovaK7_lac_** exhibited marginally less stabilization than **ovaWT**, consistent with the idea that even solvent-exposed substitutions can perturb peptide conformational dynamics within the binding groove. Activation diverged sharply from stabilization: despite preserved MHC-I display, every lysine-modified variant reduced activation to background or near-background, with **ovaK7_formyl_** the only variant measurably above background. Even subtle lysine modification therefore impairs TCR engagement at this contact residue, extending to non-enzymatic marks what we previously observed for enzymatic ones.^6^ Because this readout depends on a single SIINFEKL-specific receptor, loss of activation reports loss of recognition by that TCR rather than a global loss of immunogenicity, a distinction we return to below.

We next examined deamidation, the principal non-enzymatic modification of asparagine (position 4). The reaction proceeds through a cyclic succinimide intermediate that resolves to either aspartate or isoaspartate, the latter inserting an additional backbone methylene as a β-peptide linkage (**Fig. 2a**).^26^ Deamidated peptides have been identified directly in the MHC-I immunopeptidome, and the aspartate isomer can also arise enzymatically, since *N*-glycosidase-mediated release of an *N*-linked glycan converts the underlying asparagine to aspartate.^27^ In contrast to the lysine set, **ovaN4D** and **ovaN4D_iso_** both markedly reduced surface pMHC (**Fig. 2c**), consistent with prior reports that backbone modifications alter pMHC stability.^28^ The two isomers then diverged: **ovaN4D_iso_** retained both higher stability and measurable T cell activation, whereas **ovaN4D** elicited no detectable response. That both parameters moved in the same direction is itself notable, since modifications more often preserve one at the expense of the other. Incorporating the β-carboxyl into the backbone lengthens the mainchain by one methylene and reorients the acidic group. We propose that this repositions the epitope so that contacts required for both stable presentation and productive TCR engagement are partially preserved. Position 4 is known to tolerate substantial side-chain variation,^29^ and the present data extend that permissiveness, in attenuated form, to a change in backbone connectivity. Subtle changes in backbone connectivity, independent of side-chain identity, can therefore exert non-intuitive effects on both parameters.

We next asked whether a non-enzymatic PTM could mimic a canonical residue closely enough to drive recognition, generating a functional epitope through chemistry rather than sequence. Reactive oxygen species oxidize cysteine thiols stepwise through sulfenic and sulfinic intermediates to a terminal sulfonic acid,^30^ and oxidative modifications are known to generate neoantigens that escape central tolerance.^31^ In pancreatic β cells, oxidative stress converts a redox-sensitive cysteine of the insulin β-chain to serine, generating an HLA class II neoepitope that expands autoreactive T cells, and the same residue carries cysteinylation, glutathionylation, and the cysteic acid mark we install synthetically here.^32^ We therefore replaced the native glutamate at position 6 with cysteine (**ovaE6C**) to remove the negative charge, then oxidized it to the sulfonic acid (**ovaE6C_SO3_**) to reintroduce an acidic group that is sulfur-centered rather than carboxyl-based (**Fig. 2a**). We also prepared an *S*-succinated variant (**ovaE6C_2sc_**), in which fumarate-mediated Michael addition appends a branched dicarboxylic acid.^33^ All three peptides stabilized H-2K^b^ to comparable levels (**Fig. 2d**), but T cell activation was far more discriminating. The **ovaE6C** substitution itself partially reduced activation, consistent with the native glutamate contributing to, but not being strictly required for, TCR engagement. Restoring the negative charge did not recover that activation: **ovaE6C_SO3_** was indistinguishable from **ovaE6C**, and **ovaE6C_2sc_** reduced activation further still. Reintroducing an anionic group at position 6 therefore does not reconstitute the recognition lost when glutamate is removed, and charge equivalence alone cannot be the operative variable. These results argue against a simple charge-based model of mimicry and implicate the geometry, length, and connectivity of the native side chain. We test this directly below with an isosteric lysine surrogate (**Fig. 6b**).

Finally, because the parent epitope lacks a tyrosine, we installed one at position 5 (**ovaF5Y**), a site that tolerates aromatic substitution,^29^ as the baseline for two physiologically distinct modifications: 3-nitrotyrosine (**ovaF5Y_NO2_**), generated by peroxynitrite and widely used as a marker of nitrosative stress,^34^ and 3-chlorotyrosine (**ovaF5YCl**), which arises from hypochlorous acid produced by the neutrophil myeloperoxidase system.^35^ **ovaF5Y** activated T cells comparably to **ovaWT**, and both modifications reduced activation, nitration slightly more than chlorination (**Fig. 2e**). Each appends an electron-withdrawing substituent at a residue held tightly within the binding groove.^29^ Notably, the two assays ranked the modifications oppositely: chlorination markedly destabilized pMHC while nitration left stabilization near **ovaF5Y** levels. Because activation was assayed at 5 µM, which saturates stabilization for both variants, the reduced activation by **ovaF5Y_NO2_** cannot be attributed to lower surface display. A single ring substituent can therefore act on pMHC stability and TCR recognition in opposite directions, reinforcing that these two properties are separable.

### Non-enzymatic PTMs on cancer-associated peptides

Recent proteomic work indicates that non-enzymatic modifications substantially shape the cancer immunopeptidome. The most comprehensive catalog to date mapped thousands of PTM-bearing antigens in MC38 mouse colon cancer cells and found that non-enzymatic modifications, predominantly oxidation, deamidation, and cysteinylation, account for the majority of modified MHC-I ligands.^36^ Mass spectrometry establishes that these peptides are present but cannot resolve how a given modification affects pMHC binding. To address that gap, we synthesized defined variants of five peptides previously identified on cancer cells, introducing each modification at its reported position (**Supplementary Fig. S1**). Because cognate TCRs are not available for these epitopes, this series was evaluated for stabilization only. Methionine oxidation disrupted H-2K^b^ display in both peptides carrying it, **ca1** (KGMNYTVRL, position 3) and **ca2** (SAPENAVRM, position 9), and deepening the oxidation state of **ca1** from sulfoxide (**ca1_oxide_**) to sulfone (**ca1_sulfone_**) reduced display further. The largest effect was in **ca3** (INFDFPKL), where deamidation at position 2 lowered display by more than 100-fold, plausibly because it introduces a negative charge at a position that contributes to anchoring within the groove. Cysteinylation, by contrast, was well tolerated: adducts at position 1 of **ca4** (CGYEFTSKL) and position 7 of **ca5** (VAYEYLCHL) left stability largely unchanged, and HPLC confirmed that the cystine linkage remained intact in culture medium over the assay (**Supplementary Figs. S2,S3**), consistent with prior reports of cysteinylated peptides as immunodominant viral epitopes and H-Y minor histocompatibility antigens.^8, 37^ Modifications that lower surface pMHC could, in principle, function as an immune-evasion route by withdrawing immunogenic epitopes from display, a prediction that could be tested by comparing the modification burden of epitopes retained and lost during tumor immunoediting.

### Chemical enrichment of non-enzymatically modified MHC-I peptides

The defined epitopes above are tractable because each is a single pure species. On the surface of a cell, by contrast, such adducts exist as a vanishingly small fraction of a highly complex mixture. We therefore sought proof of concept that a non-enzymatically modified ligand could be captured directly from the cellular immunopeptidome. We focused on lysine acylation, a class that is endogenously abundant and, unlike the oxidative and carbonyl adducts above, addressable with an existing bioorthogonal probe. The central obstacle is abundance: non-enzymatic modifications are typically sub-stoichiometric, leaving the modified ligand rare against the bulk of unmodified MHC-eluted peptides.^38^ IMAC enrichment of MHC-I phosphopeptides established the key precedent for isolating such species, but it exploits an intrinsic, metal-coordinating handle unique to phosphopeptides,^39^ whereas stochastic non-enzymatic adducts carry no comparable feature and require that one be installed chemically.

To install such a handle, we employed a thioester-based probe (**1**, **Fig. 3a**) previously shown to label sites of non-enzymatic lysine acylation selectively.^40^ Probe **1** is itself a reactive thioester, the same acyl-donor chemistry through which acyl-CoA species acylate lysines non-enzymatically, so it reports intrinsic chemical reactivity rather than enzyme-directed modification and marks the lysines most prone to spontaneous acylation. Treating MDA-MB-231 cells with increasing concentrations of **1**, followed by CuAAC conjugation to fluorescein azide, produced a dose-dependent rise in cellular fluorescence (**Fig. 3b**), and in-gel fluorescence of the lysates confirmed robust labeling of protein targets (**Fig. 3c**). We then applied the strategy to the immunopeptidome (**Fig. 3d**). MHC-I-associated peptides from **1**-treated cells were recovered by immunoaffinity purification,^41^ conjugated by CuAAC to an acid-cleavable DADPS-biotin azide linker, captured on streptavidin beads, released under acid, and sequenced by LC-MS/MS. This workflow recovered two MHC-I peptides bearing the modification introduced by **1** (**Fig. 3e,f**). We present this as proof of concept that acylation-susceptible MHC-I ligands can be captured and sequenced from cells, not as evidence that any protein class is preferentially modified, a claim that two peptides cannot support. The workflow establishes a chemical-proteomic route to a ligand class that may play underappreciated roles in immune surveillance and tumor immunogenicity.

**Figure 3.**
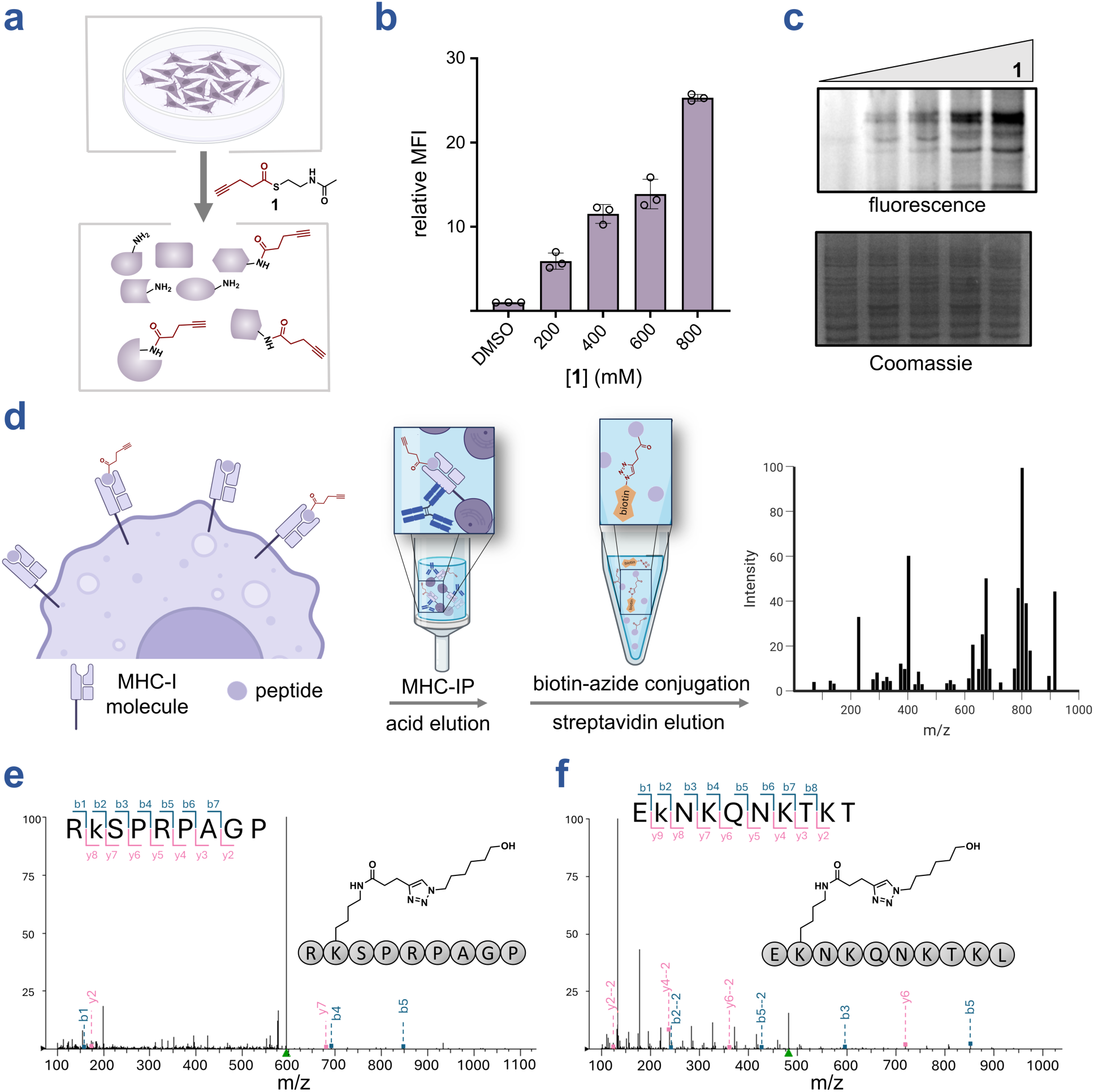
**(a)** Schematic representation of labeling sites of non-enzymatic modifications by probe **1**. **(b)** Flow cytometry analysis of MDA-MB-231 cells treated with the indicated concentration of **1** followed by a reaction with FAM-N_3_ in the presence of CuSO_4_, ascorbic acid, and THPTA. MFI is the mean fluorescence intensity of the level of fluorescence relative to the DMSO control. Data are represented as mean ± SD (*n* = 3) of technical replicates. **(c)** In-gel fluorescence analysis of **1**-treated MDA-MB-231 cells after labeling with FAM-N_3_ in the presence of CuSO_4_, ascorbic acid, and THPTA and resolved by SDS-PAGE. **(d)** Schematic of the chemical enrichment workflow using **1**. Peptides from the immunopeptidome of **1**-treated cells were reacted with DADPS-biotin azide and enriched on streptavidin beads prior to LC-MS/MS analysis. Representative LC-MS/MS spectra of modified peptides **(e)** RKSPRPAGP and **(f)** EKNKQNKTKL. Detected b+ ion fragments are shown in blue, and y+ ion fragments are shown in pink.

### Environmental electrophiles and pharmaceuticals modify antigenic peptides

The modifications considered thus far arise from endogenous metabolism. The exogenous burden is different: evolutionarily unfamiliar and developmentally unscheduled, and unlikely to be represented in the thymic peptide pool at the time the repertoire is selected. Yet these compounds react at the same nucleophilic side chains as endogenous adducts, and because exposure is often chronic, the resulting modifications can persist over a lifetime. Electrophilic pesticides, herbicides, and fungicides are a particularly understudied class in this regard. Owing to their inherent reactivity (**Fig. 4a**), these compounds covalently modify nucleophilic side chains in proteins,^10^ a process termed haptenation. Such modifications are not uniform; the local microenvironment dictates side-chain nucleophilicity, creating hyper-reactive hotspots that serve as the primary route to immunogenic neoepitopes.^42^ Prior chemoproteomic work has mapped where such compounds react across the proteome^10^ but has not asked whether the residues they modify are displayed to the immune system.

**Figure 4.**
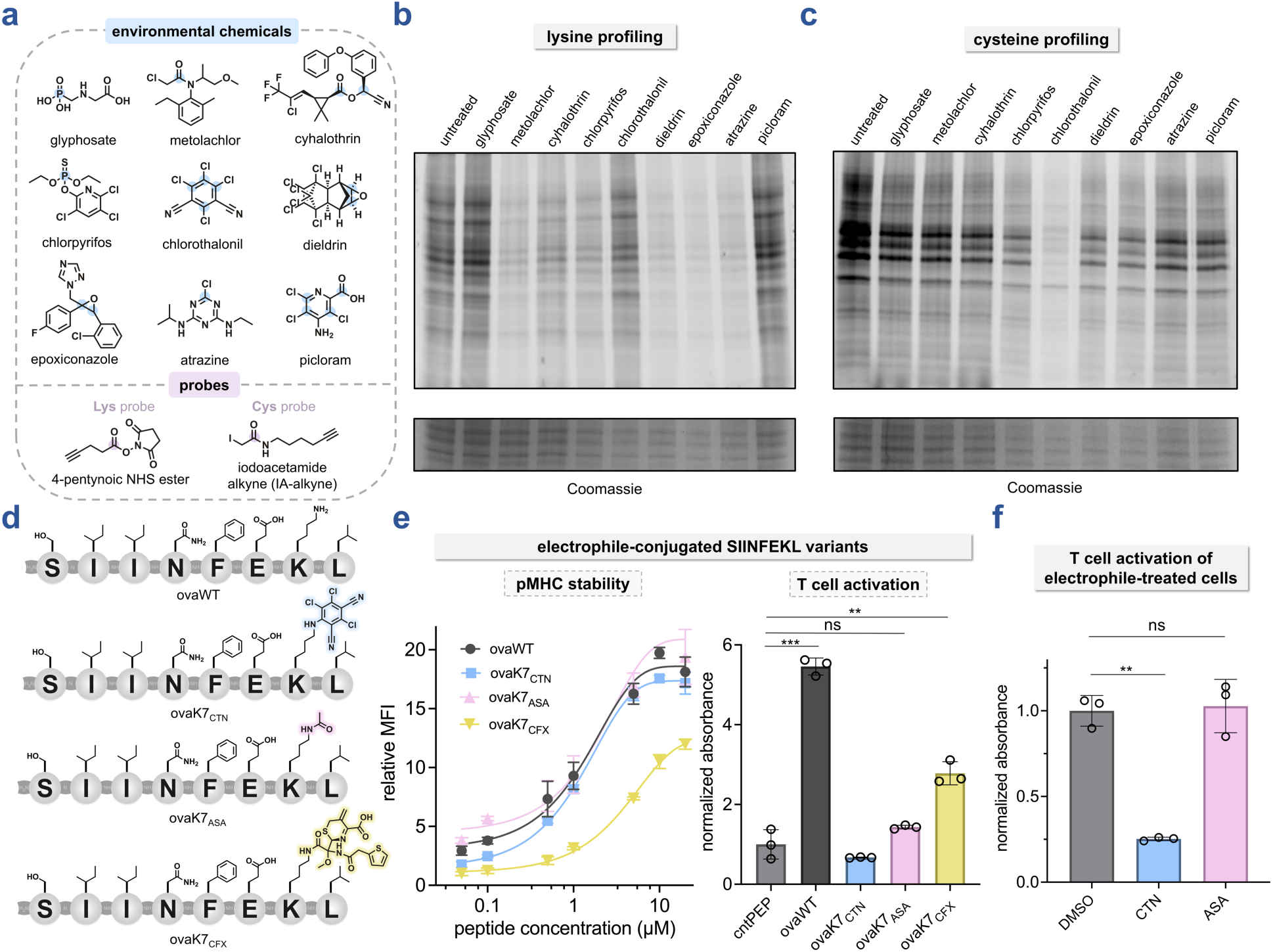
**(a)** Chemical structures of exogenous environmental chemical species and alkyne probes for both lysine- and cysteine-reactivity. Electrophilic centers are highlighted in blue (for environmental chemicals) and purple (for probes). **(b – c)** In-gel fluorescence analyses of environmental chemicals with **(b)** lysine- and **(c)** cysteine-reactive sites on proteins in cell lysates. MDA-MB-231 cell lysates were treated with electrophilic environmental chemicals (100 µM, 30 min) then competed with **(b)** 4-pentynoic NHS ester or **(c)** IA-alkyne (100 µM, 30 min) labeling, followed by a reaction with FAM-N_3_ in the presence of CuSO_4_, ascorbic acid, and THPTA. Reactivity was resolved by SDS-PAGE and visualized by in-gel fluorescence. **(d)** Chemical structures of **ovaWT**, **ovaK7_CTN_**, **ovaK7_ASA_**, and **ovaK7_CFX_**. **(e)** Dose-response curves from flow cytometry analysis of the RMA-S stabilization assay, and T cell activation from B3Z assay for electrophile-modified SIINFEKL variants. For the RMA-S assay, RMA-S cells were incubated with the indicated concentration of peptides and analyzed via flow cytometry for H-2K^b^ expression by APC anti-mouse H-2K^b^ antibody. MFI is the mean fluorescence intensity of the level of fluorescence relative to the negative control peptide (**cntPEP**). Boltzmann sigmoidal curves were fitted to the data using GraphPad Prism. For the B3Z T cell activation assay, RMA-S cells were incubated with 5 μM of indicated peptides for 1 h at 26°C. RMA-S cells were subsequently co-incubated with B3Z T cells for 6 h at 37°C. β-galactosidase expression was then measured via the conversion of the colorimetric reagent chlorophenol red-β-D-galactopyranoside (CPRG) on a plate reader at 570 nm. Data for both assays were normalized to the negative control peptide (**cntPEP**) and are represented as mean ± SD (*n* = 3) of technical replicates. *P*-values were determined by a two-tailed *t*-test (ns = not significant, ** *p* < 0.01, *** *p* < 0.001). **(f)** B3Z T cell activation assay with electrophile-treated B16-OVA cells. IFN-γ-pre-treated B16-OVA cells were incubated with 100 µM of CTN or 1 mM of ASA for 1.5 h, washed, then co-incubated with B3Z T cells for 6 h. β-galactosidase expression was then measured via the conversion of the colorimetric reagent chlorophenol red-β-D-galactopyranoside (CPRG) on a plate reader at 570 nm. The data presented have been normalized to the absorbance of the DMSO control. Data are represented as mean ± SD (*n* = 3) of technical replicates. *P*-values were determined by a two-tailed *t*-test (ns = not significant, ** *p* < 0.01).

We therefore used competitive activity-based protein profiling to rank the proteome-wide reactivity of a panel of cysteine- and lysine-reactive herbicides, pesticides, and fungicides. Chemicals were pre-incubated with lysates (for gel-based experiments) or intact cells (for flow cytometry-based assays), and the unoccupied cysteine and lysine sites were labeled with an iodoacetamide alkyne (IA-alkyne) or NHS-ester alkyne probe,^43^ so that reduced fluorescence reports increased reactivity of the competitor. Lysine-directed labeling produced weaker and less discriminating signals (**Fig. 4b** and **Supplementary Fig. S6**), likely reflecting competing hydrolysis of the NHS-ester probe and its reported reactivity toward serine and threonine.^44^ Cysteine-directed labeling was considerably more informative, with the largest reductions in fluorescence following chlorothalonil treatment (**Fig. 4c** and **Supplementary Fig. S10**). Chlorothalonil was at least as reactive as the positive control iodoacetamide toward cellular cysteines and was therefore selected for subsequent immunological studies. Acetylsalicylic acid (ASA; aspirin), which acetylates serine, lysine, and other side chains, produced no substantial reduction in lysine-directed labeling (**Supplementary Figs. S11,S12**), consistent with its slower acetylation kinetics and with the low site occupancy reported in the absence of deacetylase inhibition, which would otherwise allow acetyl marks to accumulate.^45^

Guided by these profiles, we synthesized SIINFEKL variants bearing exogenous adducts at lysine 7 (**Fig. 4d**): the chlorothalonil hapten (**ovaK7_CTN_**), the acetylated product of ASA (**ovaK7_ASA_**), and a β-lactam adduct from the antibiotic cefoxitin (**ovaK7_CFX_**). β-Lactams provide a clinically grounded comparator: they are among the best-characterized haptens, modify lysines on human serum albumin (HSA) to form conjugates that elicit drug-specific immune responses, and appear as naturally presented MHC-I ligands.^11, 46^ **ovaK7_CFX_** carries the rearranged adduct formed after ring opening and elimination of the C3 carbamoyloxy leaving group.^47^ Because the lysine-directed probe was insensitive to cefoxitin (**Supplementary Fig. S15**), we verified β-lactam reactivity orthogonally, treating human serum with the clickable analogue azidocillin and identifying HSA as the predominant labeled protein (**Supplementary Fig. S16**).

Using the RMA-S stabilization assay, both **ovaK7_CTN_** and **ovaK7_ASA_** stabilized H-2K^b^ to a similar extent as unmodified **ovaWT**, whereas **ovaK7_CFX_** produced a modest reduction (**Fig. 4e**). Despite preserved MHC binding, neither **ovaK7_CTN_** nor **ovaK7_ASA_** elicited detectable T cell activation, while **ovaK7_CFX_** retained a measurable, albeit substantially reduced, response. Notably, **ovaK7_CFX_** was the weakest stabilizer of the three yet the only one to support activation, so its retained response cannot be explained by higher surface display. The chemistry of a covalent lysine modification, rather than the modification site alone, is therefore a key determinant of T cell recognition.

Pre-conjugated peptides afford precise control but bypass endogenous processing. We therefore adapted the T cell activation assay to B16-OVA cells, a murine melanoma line expressing full-length ovalbumin that generates SIINFEKL intracellularly for display on H-2K^b^. Chlorothalonil treatment of B16-OVA cells reduced T cell activation to background levels, whereas ASA treatment did not (**Fig. 4f**), consistent with its slower acetylation kinetics. Cefoxitin was not evaluated in this model, since β-lactam hypersensitivity is associated with haptenation of extracellular proteins rather than intracellular antigens.

### Dietary isothiocyanates modify antigenic peptides

We next investigated ITCs, reactive electrophiles abundant in cruciferous vegetables that covalently modify nucleophilic residues across the proteome (**Fig. 5a**).^12^ Across both gel-based and flow cytometry-based assay formats, phenethyl isothiocyanate (PEITC) and allyl isothiocyanate (AITC) competed robustly with cysteine-reactive probes, consistent with their known electrophilicity toward thiols, whereas sulforaphane (SFN) competed comparatively weakly (**Fig. 5b,c** and **Supplementary Fig. S14**). The corresponding lysine-modified variants **ovaK7_PEITC_** and **ovaK7_AITC_** retained MHC binding comparable to **ovaWT** but showed markedly reduced T cell activation (**Fig. 5d,e**), indicating that covalent lysine modification by dietary ITCs also renders peptides immunologically distinct. In the B16-OVA system, PEITC and AITC reduced engagement of endogenously processed SIINFEKL, whereas SFN did not (**Fig. 5f**). The ranking established against cellular cysteines is therefore preserved at a peptide that contains no cysteine, which points to a determinant other than thiol selectivity. The greater polarity of SFN is expected to limit cellular accumulation, and its sulfinyl group adds steric bulk near the electrophilic carbon, which together may restrict access to a lysine ε-amine within the constrained pMHC environment. Bulk electrophilicity alone is therefore not predictive of antigen-modifying potential; cellular accumulation, residue selectivity, adduct stability, and molecular context together determine whether a covalent modification translates into altered immune recognition. Adduct stability is a key limitation here, as both the dithiocarbamate adducts formed with cysteine and the thiourea adducts formed with lysine by ITCs are reversible on physiologically relevant timescales.

**Figure 5.**
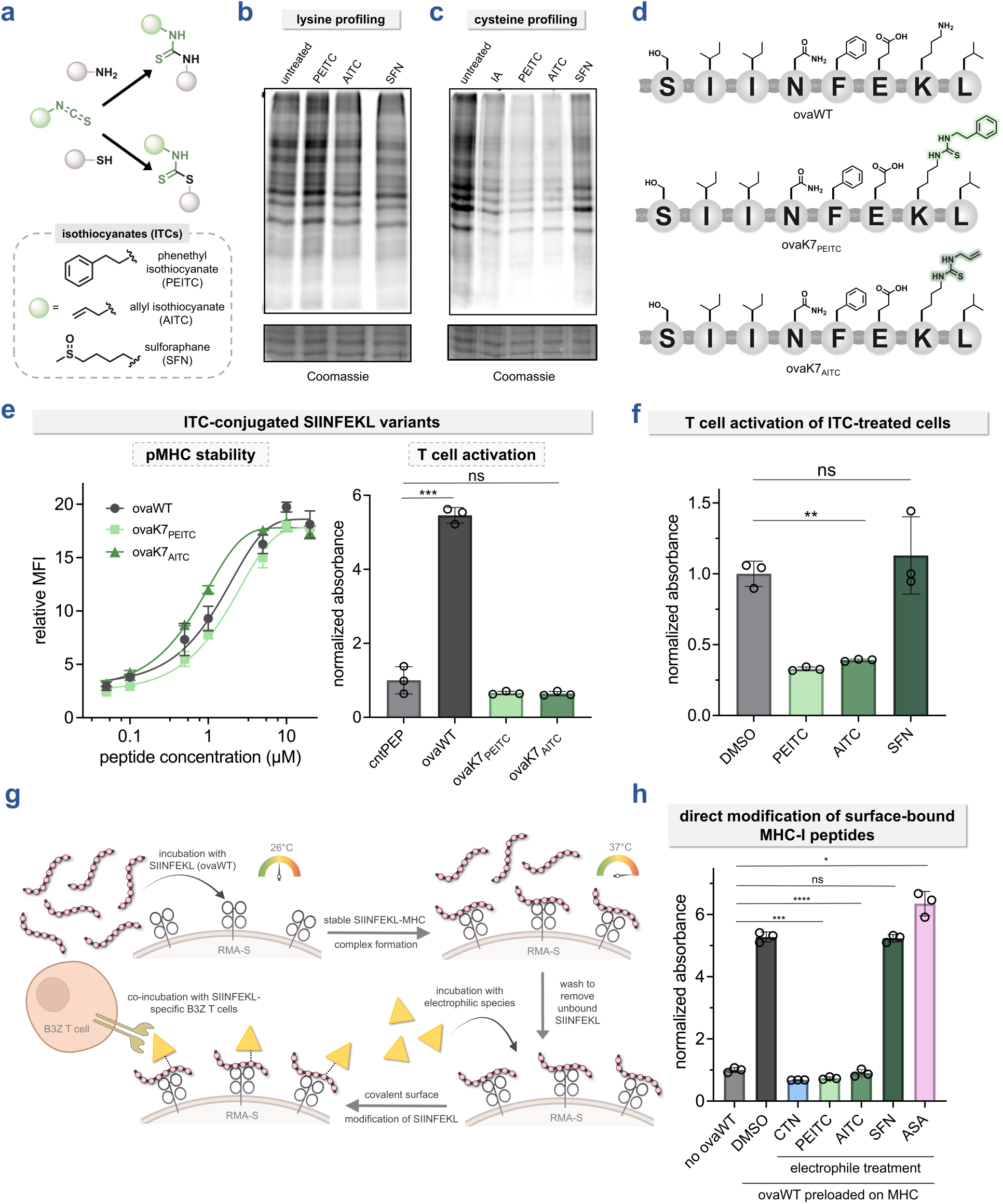
**(a)** Schematic depicting reactivity of isothiocyanate (ITC) with amines and thiols, and chemical structures of example ITCs. **(b – c)** In-gel fluorescence analyses of ITCs and aspirin (ASA) reactivity with **(b)** lysine- and **(c)** cysteine-reactive sites on proteins in cell lysates. MDA-MB-231 cell lysates were treated with positive control iodoacetamide and ITCs (100 µM, 30 min), then competed with **(b)** 4-pentynoic NHS ester or **(c)** IA-alkyne (100 µM, 30 min) labeling, followed by a reaction with FAM-N_3_ in the presence of CuSO_4_, ascorbic acid, and THPTA. Reactivity was resolved by SDS-PAGE and visualized by in-gel fluorescence. **(d)** Chemical structures of **ovaWT**, **ovaK7_PEITC_**, and **ovaK7_AITC_**. **(e)** Dose-response curves from flow cytometry analysis of the RMA-S stabilization assay, and T cell activation from B3Z assay for ITC-modified SIINFEKL variants. For the RMA-S assay, RMA-S cells were incubated with the indicated concentration of peptides and analyzed via flow cytometry for H-2K^b^ expression by APC anti-mouse H-2K^b^ antibody. MFI is the mean fluorescence intensity of the level of fluorescence relative to the negative control peptide (**cntPEP**). Boltzmann sigmoidal curves were fitted to the data using GraphPad Prism. For the B3Z T cell activation assay, RMA-S cells were incubated with 5 μM of indicated peptides for 1 h at 26°C. RMA-S cells were subsequently co-incubated with B3Z T cells for 6 h at 37°C. β-galactosidase expression was then measured via the conversion of the colorimetric reagent chlorophenol red-β-D-galactopyranoside (CPRG) on a plate reader at 570 nm. Data for both assays were normalized to the negative control peptide (**cntPEP**) and are represented as mean ± SD (*n* = 3) of technical replicates. *P*-values were determined by a two-tailed *t*-test (ns = not significant, *** *p* < 0.001). **(f)** B3Z T cell activation assay with ITC-treated B16-OVA cells. IFN-γ-pre-treated B16-OVA cells were incubated with 100 µM of ITCs for 1.5 h, washed, then co-incubated with B3Z T cells for 6 h. β-galactosidase expression was then measured via the conversion of the colorimetric reagent chlorophenol red-β-D-galactopyranoside (CPRG) on a plate reader at 570 nm. The data presented have been normalized to the absorbance of the DMSO control. Data are represented as mean ± SD (*n* = 3) of technical replicates. *P*-values were determined by a two-tailed *t*-test (ns = not significant, ** *p* < 0.01). **(g)** Schematic illustrating the B3Z T cell activation assay using RMA-S cells with SIINFEKL pre-loaded on MHC to isolate the impact of exclusively surface modification of MHC-I-presented peptides by electrophilic species. **(h)** B3Z T cell activation assay with electrophile-treated, SIINFEKL-pre-loaded RMA-S cells. RMA-S cells were incubated with 5 μM **ovaWT** for 1 h at 26°C in order to occupy empty MHC on the surface, then washed thoroughly to remove unbound **ovaWT** from the culture media. RMA-S cells were then treated with 100 µM of electrophilic compounds (except for ASA, which was 1 mM) for 1.5 h, washed, then co-incubated with B3Z T cells for 6 h at 37°C. β-galactosidase expression was then measured via the conversion of the colorimetric reagent chlorophenol red-β-D-galactopyranoside (CPRG) on a plate reader at 570 nm. The data presented have been normalized to the absorbance of the DMSO control. Data are represented as mean ± SD (*n* = 3) of technical replicates. *P*-values were determined by a two-tailed *t*-test (ns = not significant, * *p* < 0.05, *** *p* < 0.001, **** *p* < 0.0001).

### Surface-bound MHC-I peptides are directly modified

Because antigens in B16-OVA cells are generated intracellularly, the preceding experiments cannot separate modifications acquired during antigen processing from those installed directly on peptides already bound to MHC-I at the cell surface, and both routes may contribute to the observed loss of recognition. The two are differently constrained: intracellular modification requires the electrophile to accumulate and traverse the membrane, whereas surface modification could alter recognition even for a poorly permeable species. To isolate the surface route, we exploited the TAP deficiency of RMA-S cells, which restricts display to exogenously loaded peptides (**Fig. 5g**). Cells were first loaded with **ovaWT** to generate stable pMHC-I complexes, then washed extensively to remove free peptide before treatment with electrophilic compounds, confining any modification to peptides already bound to H-2K^b^. Treatment with chlorothalonil, PEITC, or AITC reduced activation to background levels, whereas SFN had no effect (**Fig. 5h**). Select electrophiles can therefore modify peptides already presented on MHC-I and disrupt TCR recognition independently of intracellular antigen processing.

### Chemically distinct cysteine adducts produce divergent recognition

Because many electrophiles converge on the same nucleophilic side chains, the immunological consequence of an exposure depends on which chemical state a residue ends up in. Occupation of a cysteine by an ITC, for example, would leave no free thiol for a later electrophile to engage. Rather than examining competing reactions *in situ*, we synthesized chemically defined cysteine adducts at position 7 of SIINFEKL and asked how each engages the SIINFEKL-specific TCR, isolating the chemistry from reaction kinetics and incomplete conversion.

Replacing lysine 7 with cysteine (**ovaK7C**, **Fig. 6a**) abolished activation, as expected for a residue acutely sensitive to charge and side-chain chemistry, while providing a uniquely reactive thiol (**Fig. 6b**). Conjugation of PEITC to this cysteine generated the corresponding dithiocarbamate (**ovaK7C_PEITC_**), which likewise produced no activation, establishing two chemically distinct yet functionally silent antigenic states against which subsequent remodeling could be assessed. We next examined chlormequat (CMQ), a quaternary ammonium plant growth regulator bearing a 2-haloethyl electrophile; human exposure is increasingly common, with recent biomonitoring detecting the compound in the majority of surveyed urine samples.^48^ The reaction of CMQ with cysteine introduces a permanently cationic 2-(trimethylammonio)ethyl substituent. This modification partially recapitulates trimethyllysine, which demonstrated appreciable activation in our previous work.^6^ Despite introducing a positively charged substituent, **ovaK7C_CMQ_** failed to activate B3Z cells (**Fig. 6b**), indicating that restoration of positive charge alone is insufficient to reproduce the structural determinants recognized by this TCR. We therefore sought a more faithful lysine surrogate: bromoethylamine converts cysteine to *S*-(2-aminoethyl)cysteine (thialysine), an established lysine analogue widely used in protein engineering.^49^ RMA-S cells pulsed with **ovaK7C_BrEA_** elicited robust T cell activation (**Fig. 6b**), establishing thialysine as a functional lysine mimic within this epitope. Chemically distinct electrophiles acting at a single residue therefore generate antigenic states with markedly different capacities to engage the same TCR, providing a framework for how prior occupation of a reactive residue could constrain the chemical states accessible to subsequent exposures.

**Figure 6.**
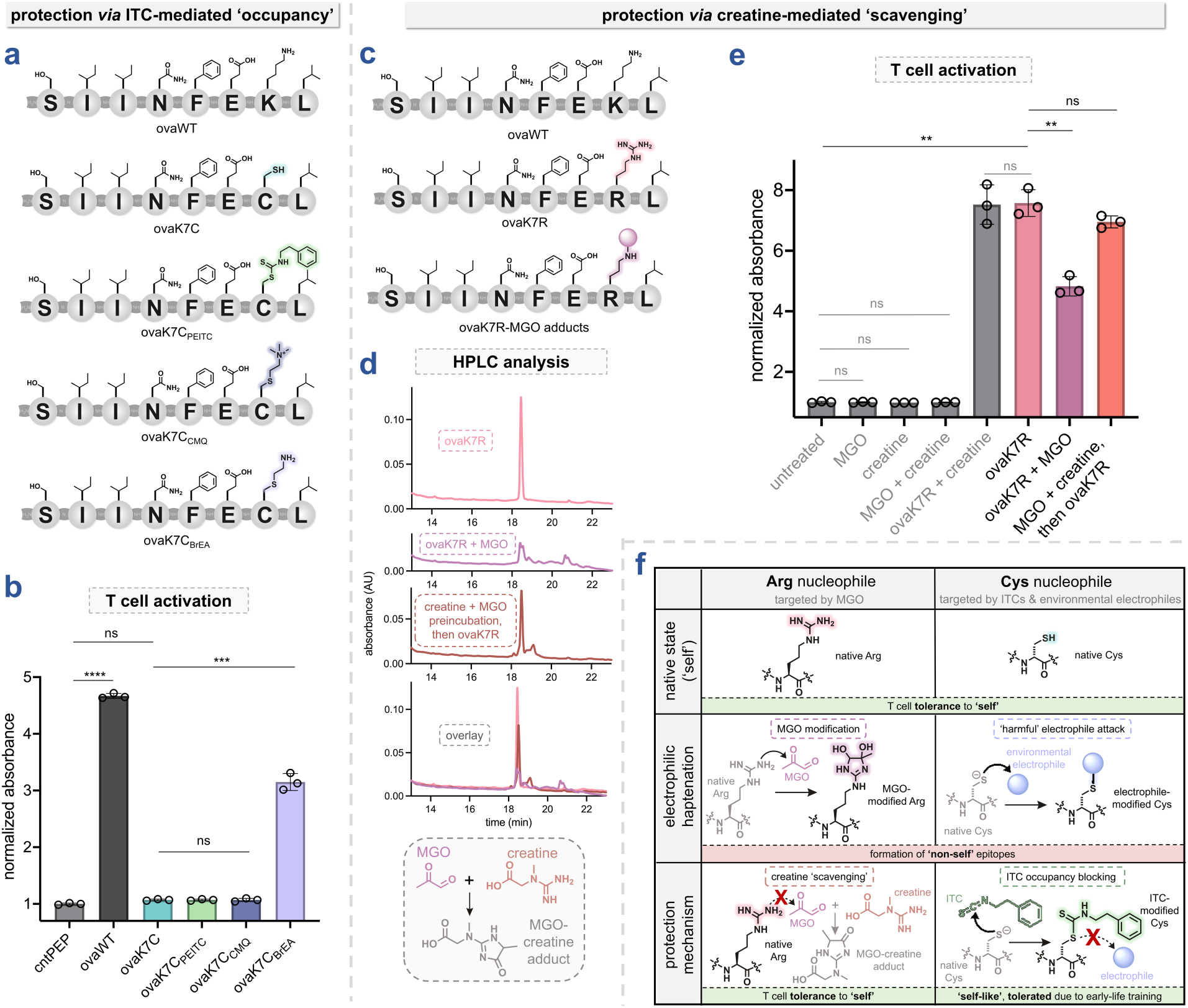
**(a – b)** ITC-mediated ‘occupancy’ protection mechanism. **(a)** Chemical structures of **ovaWT**, **ovaK7C**, **ovaK7C_PEITC_**, **ovaK7C_CMQ_**, and **ovaK7C_BrEA_**. **(b)** T cell activation from B3Z assay for cysteine-modified SIINFEKL variants. RMA-S cells were incubated with 5 μM of indicated peptides for 1 h at 26°C. RMA-S cells were subsequently co-incubated with B3Z T cells for 6 h at 37°C. β-galactosidase expression was then measured via the conversion of the colorimetric reagent chlorophenol red-β-D-galactopyranoside (CPRG) on a plate reader at 570 nm. The data presented have been normalized to the absorbance of the negative control peptide (**cntPEP**) and are represented as mean ± SD (*n* = 3) of technical replicates. *P*-values were determined by a two-tailed *t*-test (ns = not significant, *** *p* < 0.001, **** *p* < 0.0001). **(c – e)** Creatine-mediated ‘scavenging’ protection mechanism. **(c)** Chemical structures of **ovaWT**, **ovaK7R**, and **ovaK7R**-MGO adducts (representative). **(d)** HPLC analysis of **ovaK7R** alone, **ovaK7R** incubated with 50-fold excess MGO (24 h, 37°C), or incubated with MGO and creatine mixture (24 h, 37°C) that had been pre-reacted together (2 h, 37°C, 10-fold excess creatine to MGO). Reaction mixtures were analyzed using RP-HPLC monitoring absorbance at 220 nm. **(e)** B3Z T cell activation assay with reaction mixture of MGO-modified and creatine-scavenged **ovaK7R**. RMA-S cells were incubated with a 500-fold dilution (equivalent to 100 nM **ovaK7R**, 5 mM MGO, and 50 mM creatine) of various reaction mixtures for 1 h at 26°C, washed thoroughly, then co-incubated with B3Z T cells for 6 h at 37°C. β-galactosidase expression was then measured via the conversion of the colorimetric reagent chlorophenol red-β-D-galactopyranoside (CPRG) on a plate reader at 570 nm. The data presented have been normalized to the absorbance of the untreated control. Data are represented as mean ± SD (*n* = 3) of technical replicates. *P*-values were determined by a two-tailed *t*-test (ns = not significant, ** *p* < 0.01). **(f)** Proposed complementary mechanisms by which creatine and ITCs may limit electrophile-driven antigen remodeling. In the absence of protection, reactive electrophiles modify susceptible amino acid side chains, generating chemically altered epitopes with the potential to modify T cell recognition. Creatine limits arginine modification by scavenging MGO prior to protein adduction, whereas ITCs are proposed to limit cysteine modification by occupying reactive thiol sites and competing with other electrophiles. Despite acting through distinct mechanisms, both interventions reduce the formation of alternative electrophile-derived epitopes and may help maintain immunological recognition of ‘self’.

### MGO glycation impairs recognition and is prevented by creatine

Having shown that electrophiles can remodel antigens both during processing and at the cell surface, we returned to an endogenous reactive species to ask the converse question: if these adducts are the products of chemical reactions, can they be prevented by controlling the reactive species itself? MGO non-enzymatically glycates lysine and arginine side chains, along with *N*-terminal amines, to form advanced glycation end products (AGEs),^24^ which are strongly associated with aging and metabolic disease.^50^ The underlying chemistry is heterogeneous, with nearly 40 described AGEs that interconvert through successive condensation and rearrangement,^24^ so isolating a single representative adduct remains challenging.

To generate a defined MGO-reactive substrate, we replaced the position 7 lysine with arginine (**ovaK7R**, **Fig. 6c**), introducing a guanidinium acceptor while maintaining the native positive charge; **ovaK7R** retained appreciable activation, establishing a functional baseline (**Fig. 6e**). Rather than isolating a single product, we glycated **ovaK7R** with excess MGO and evaluated the resulting mixture directly. HPLC showed substantial depletion of the parent peptide peak and the emergence of several new peaks (**Fig. 6d**), and LC-MS identified masses consistent with previously reported MGO-derived AGEs, including dihydroxyimidazolidine, the hydroimidazolone isomers MGH-1 and MGH-2, carboxyethylarginine, and tetrahydropyrimidine (**Supplementary Figs. S18,S19)**.^51^ RMA-S cells pulsed with the glycated mixture produced a significant reduction in activation relative to unmodified **ovaK7R** (**Fig. 6e**). Because the mixture still contains a fraction of unmodified peptide, and because the enzymatic reporter amplifies even weak engagement, this reduction is a conservative measure of the loss caused by glycation.

We next asked whether interception of the reactive carbonyl could preserve recognition. Creatine, a guanidinium-containing metabolite and one of the most widely consumed dietary supplements, reacts directly with MGO under physiological conditions and scavenges more than half of ingested MGO in the human stomach.^52^ Western blotting with an anti-MGO adduct antibody confirmed that creatine co-treatment markedly reduced adduction at the proteome level (**Supplementary Fig. S20**). Pre-incubating MGO with excess creatine before adding **ovaK7R** substantially preserved the parent peptide, as observed by HPLC (**Fig. 6d**). In the activation assay, this preserved peptide yielded responses comparable to unmodified **ovaK7R** and significantly higher than those of the MGO-treated peptide (**Fig. 6e**). Control conditions (MGO alone, creatine alone, and MGO plus creatine without peptide) produced only background signals. Consequently, carbonyl scavenging preserves antigen recognition by restricting the formation of immunologically disruptive glycation products.

## DISCUSSION

We show that non-enzymatic chemical modifications, whether arising from endogenous metabolites or exogenous electrophiles, can substantially reshape MHC-I antigen presentation and T cell recognition. Across defined synthetic and cancer-derived epitopes, single modifications altered pMHC-I stability and frequently disrupted recognition even when MHC binding was preserved, and a chemical enrichment strategy allowed non-enzymatically acylated MHC-I ligands to be identified directly from cells. A single principle links these observations: antigenic identity is determined not only by peptide sequence but by the cumulative chemical history of the antigen, the sum of electrophilic exposures that install, alter, or prevent specific side-chain modifications. Our results further identify two levers on that history: the identity of the adduct formed and the availability of the species that forms it.

Modified peptides remain challenging to detect by conventional approaches, owing to their low stoichiometric abundance and the unpredictable effects of PTMs on fragmentation and ionization,^38^ which impose sensitivity and search-space limits on discovery pipelines. The KRAS(G12C) inhibitor ARS1620 illustrates the consequence: it acts as a hapten, generating a drug-peptide neoepitope presented on MHC-I that can serve as a direct target for cytotoxic T cell immunotherapy, yet the haptenated peptide has not been isolated from cultured cells by standard immunopeptidomic workflows, its existence established functionally instead.^53^ This case reflects a broader challenge for the field. Establishing that a modified peptide drives physiology, beneficially as in haptenated neoantigen vaccines or pathologically as in xenobiotic-induced hypersensitivity, requires linking a defined chemical modification to a specific immunological outcome. Where that causal chain has been completed, the immune response is acute and the modification effectively obligatory, as in celiac disease, in which transglutaminase-mediated deamidation of gliadin peptides generates negatively charged epitopes with enhanced binding to HLA-DQ2 and HLA-DQ8 and subsequent activation of pathogenic CD4+ T cells.^54^ Chronic, low-grade immunopathology is far less tractable: the modification may be sub-stoichiometric, the phenotype diffuse, and no single antigen-withdrawal experiment decisive. In such settings, functional immunological readouts become a necessary substitute for direct physical detection, which is the rationale for the bottom-up strategy taken here. Synthesizing homogeneous peptides bearing a single specified modification addresses the two questions that gate relevance, whether the modification is tolerated for stable presentation and whether the resulting complex still engages T cells, and identifies which marks merit the harder effort of locating them in the endogenous immunopeptidome.

The same logic explains why proteome-wide reactivity is a poor proxy for immunological impact. T cell responses are frequently driven by a small number of epitopes, so modification of one immunologically relevant residue could substantially alter recognition while producing minimal change across the bulk proteome. The modest shifts in our gel-based and flow cytometric analyses should therefore not be read as evidence that electrophile-induced remodeling is insignificant; these formats can overlook rare but consequential events within specific antigenic proteins.

An important caveat of this study is that activation was assessed using a single SIINFEKL-specific TCR reporter. We relied on the B3Z system because analytical tools for evaluating antigen-specific T cell recognition remain limited, and comparable high-throughput alternatives are not currently available. Although primary T cells could, in principle, provide a broader assessment of immunogenicity, they are highly impractical for systematically screening large panels of post-translationally modified peptides. Primary cell assays introduce substantial biological variability, contain exceedingly low frequencies of antigen-specific T cells, and are largely incompatible with the high-throughput formats required to evaluate diverse chemical adducts. Consequently, a loss of activation in our surrogate model should not be interpreted as a universal loss of immunogenicity. Instead, chemical modification may disrupt recognition by one TCR while simultaneously generating epitopes that remain recognizable, or are even preferentially recognized, by other TCRs. Against this backdrop, our findings place chemically distinct electrophiles on equal footing: compounds typically regarded as harmful (e.g., environmental pesticides) and those considered beneficial (e.g., isothiocyanates from cruciferous vegetables) can form covalent adducts at the same cysteine and lysine side chains within peptide antigens. The underlying chemistry is agnostic to whether an electrophile originates from a dietary metabolite or an environmental toxicant; instead, the immunological outcome is dictated by the identity of the resulting adduct and the context in which it is first encountered, not by the toxicological classification of the compound.

Our **ovaK7C** series further suggests that such modifications should not be considered in isolation. Because many electrophiles target the same side chains, prior modification by one can influence or preclude modification by another, and ITC modification may act not merely as a source of altered antigens but as a form of chemical occupancy that limits formation of alternative, electrophile-driven neoepitopes (**Fig. 6f**). One limitation qualifies this interpretation: in our surrogate model, the competing reaction restored near-native recognition rather than generating a distinct neoepitope. Consequently, the hypothesis that occupancy forecloses more hazardous states remains to be tested in a system where the blocked reaction is inherently immunogenic. Regarding exposure timing, modification during thymic education might simultaneously promote tolerance toward the occupied site and limit subsequent neoepitope formation by more hazardous electrophiles.

Our findings with MGO extend this concept to endogenous metabolic byproducts. Despite the heterogeneity of AGE chemistry, MGO exposure alone was sufficient to impair antigen-specific activation. This impairment was reversed by creatine, an endogenous metabolite that scavenges MGO prior to peptide modification; to our knowledge, its influence on antigen presentation has not been previously examined. Because AGEs accumulate with age and metabolic disease, they may represent a particularly relevant class of late-emerging, altered-self antigens that arise outside the window of thymic selection. More broadly, these findings establish chemically modified peptides as an underexplored dimension of antigenic diversity, shaped by the cumulative effects of metabolism, environmental exposure, diet, and therapeutics. Furthermore, our work provides both mechanistic evidence and experimental tools to define how such epitopes contribute to cancer immunity, inflammatory disease, and autoimmunity.

## METHODS

### Experimental Methods

#### Mammalian Cell Culture

RMA-S cells were a kind gift from Dr. John Sampson. B3Z cells were kindly provided by Dr. Aaron Esser-Kahn. B16-OVA cells were generously provided by Dr. Timothy Bullock. RMA-S and B3Z cells were maintained in RPMI 1640 media supplemented with 10% fetal bovine serum, 50 IU/mL penicillin, and 50 µg/mL streptomycin. B16-OVA cells were maintained in RPMI 1640 medium, supplemented with 10% fetal bovine serum, 50 IU/mL penicillin, 50 µg/mL streptomycin, and 1 mg/mL G418 (geneticin). MDA-MB-231 cells were maintained in Dulbecco’s Modified Eagle Medium (DMEM) supplemented with 10% fetal bovine serum (FBS), 50 IU/mL penicillin, and 50 µg/mL streptomycin. All cells were cultured in T75 flasks and maintained in a humidified atmosphere of 5% CO_2_ at 37°C.

#### RMA-S Stabilization Assay

10^5^ RMA-S cells were seeded in a treated 96 well plate at 37°C overnight. The next day, RMA-S cells were moved to a 26°C incubator for 24 hours. Following the incubation period, cells were incubated with peptides in culture media at indicated concentrations for 1 hour at 26°C before being moved to the 37°C incubator for 6 hours. Cells were removed from the well plate by vigorous pipetting and transferred to a round-bottom 96-well plate. Transferred cells were centrifuged (1100 x g, 5 min) in a Thermo Scientific Jouan C4i centrifuge and the supernatant was removed. Cell pellets were resuspended in a 1:100 dilution of APC-labeled anti-mouse H-2K^b^ antibody in culture media for 1 hour at 4°C. Following centrifugation and removal of the supernatant, cells were then fixed with 4% formaldehyde solution, and analyzed using the Attune NxT Flow Cytometer (Thermo Fisher) equipped with a 637 nm laser with 670/14 nm bandpass filter.

#### B3Z T Cell Activation Assays

##### Co-Incubation with RMA-S cells (for modified SIINFEKL peptides)

10^5^ RMA-S cells were seeded in a treated 96 well plate at 37°C overnight. The next day, RMA-S cells were moved to a 26°C incubator for 24 hours. Following the incubation period, cells were incubated with peptides in culture media at indicated concentrations for 1 hour at 26°C. After this time, 1.5 x 10^4^ B3Z cells in culture media were co-incubated with the RMA-S cells for 6 hours. Cells were centrifuged (1100 x g, 5 min) in a Thermo Scientific Jouan C4i centrifuge and the supernatant was removed. Lysis buffer containing 0.2% saponin, 500 mM CPRG reagent (Chlorophenol Red β-D-Galactopyranoside), 20 mM MgCl_2_, and 100 mM β-mercaptoethanol in 1X PBS was added to each well. After 1 hour, absorbance at 570 nm was recorded using a BioTek Synergy H1 Microplate Reader.

##### Co-Incubation with RMA-S cells (for electrophile treatment)

10^5^ RMA-S cells were seeded in a treated 96 well plate at 37°C overnight. The next day, RMA-S cells were moved to a 26°C incubator for 24 hours. Following the incubation period, cells were incubated with 5 µM **ovaWT** in culture media 1 hour at 26°C. After this time, cells were washed twice with 1X PBS, then treated with indicated concentration of electrophilic compounds in culture media (with no FBS) at 37°C for 1.5 hours. Cells were then washed twice with 1X PBS, and the culture media was replaced with media containing 1.5 x 10^4^ B3Z cells, which were co-incubated with the RMA-S cells for 6 hours. Cells were centrifuged (1100 x g, 5 min) in a Thermo Scientific Jouan C4i centrifuge and the supernatant was removed. Lysis buffer containing 0.2% saponin, 500 mM CPRG reagent (Chlorophenol Red β-D-Galactopyranoside), 20 mM MgCl_2_, and 100 mM β-mercaptoethanol in 1X PBS was added to each well. After 2 hours, absorbance at 570 nm was recorded using a BioTek Synergy H1 Microplate Reader.

##### Co-Incubation with RMA-S cells (for MGO-modified ovaK7R reaction treatment)

10^5^ RMA-S cells were seeded in a treated 96 well plate at 37°C overnight. The next day, RMA-S cells were moved to a 26°C incubator for 24 hours. Following the incubation period, cells were incubated with a 500x dilution (equivalent to 100 nM of **ovaK7R**, 5 mM MGO, and 50 mM creatine) of various reaction mixtures (see “MGO Modification of ovaK7R and Creatine Scavenging Reactions” section below) in culture media 1 hour at 26°C. After this time, cells were washed twice with 1X PBS, and the culture media was replaced with media containing 1.5 x 10^4^ B3Z cells, which were co-incubated with the RMA-S cells for 6 hours. Cells were centrifuged (1100 x g, 5 min) in a Thermo Scientific Jouan C4i centrifuge and the supernatant was removed. Lysis buffer containing 0.2% saponin, 500 mM CPRG reagent (Chlorophenol Red β-D-Galactopyranoside), 20 mM MgCl_2_, and 100 mM β-mercaptoethanol in 1X PBS was added to each well. After 2 hours, absorbance at 570 nm was recorded using a BioTek Synergy H1 Microplate Reader.

##### Co-Incubation with B16-OVA Cells (for electrophile treatment)

5 x 10^4^ B16-OVA cells were seeded in a treated 96 well plate and pretreated with 10 ng/mL IFN-γ. The next day, the culture media was replaced with culture media (with no FBS) containing electrophilic compounds at indicated concentrations at 37°C for 1.5 hours. Cells were washed twice with 1X PBS, and the culture media was replaced with media containing 1.5 x 10^4^ B3Z cells, which were co-incubated with the B16-OVA cells for 6 hours. As before, cells were centrifuged (1100 x g, 5 min) in a Thermo Scientific Jouan C4i centrifuge and the supernatant was removed. Lysis buffer containing 0.2% saponin, 500 mM CPRG reagent (Chlorophenol Red β-D-Galactopyranoside), 20 mM MgCl_2_, and 100 mM β-mercaptoethanol in 1X PBS was added to each well. After 2 hours, absorbance at 570 nm was recorded using a BioTek Synergy H1 Microplate Reader.

#### Flow Cytometry Analysis of Non-Enzymatic Acylation with 1

MDA-MB-231 cells were seeded in a treated 96 well plate at 37°C until reaching ∼80-90% confluency. Cells were incubated with **1** at various concentrations overnight. Cells were washed once with 1X PBS, removed using TrypLE^TM^ Express Enzyme (Thermo Fisher), and transferred to a round-bottom 96-well plate. Transferred cells were centrifuged (1000 x g, 5 min) in a Thermo Scientific Jouan C4i centrifuge, and the cell pellets were fixed in 4% formaldehyde solution for 20 minutes and permeabilized with methanol. The plate was centrifuged (1000 x g, 5 min) and pelleted cells were resuspended with 20 mM FAM-N_3_ with 1 mM CuSO_4_, 128 mM THPTA, and 1.2 mM ascorbic acid for 30 min. Cells were analyzed using an Attune NxT Flow Cytometer (Thermo Fisher) equipped with a 488 nm laser with 530/30 nm bandpass filter.

#### In-Gel Fluorescence Analysis of Non-Enzymatic Acylation with 1

MDA-MB-231 cells were seeded in a treated 6 well plate at 37°C until reaching ∼80-90% confluency. Cells were washed thrice with 1X PBS, then incubated with varying concentrations of **1** at 37°C overnight. Cells were washed thrice, scraped to remove, and centrifuged (1.5 x g, 5 min) to pellet. After removing the supernatant, cell pellets were then lysed on ice with RIPA buffer. Cell lysates were subsequently reacted with 150 mM FAM-N_3_ with 1 mM CuSO_4_, 128 mM THPTA, and 1.2 mM ascorbic acid for 30 min at 37°C. Reactions were resolved by 12% SDS-PAGE (200 V, 1 h) and scanned for fluorescence using an Azure Biosystems 300Q Imaging System.

#### Competitive Flow Cytometry Analysis of Proteome-Wide Reactivity

MDA-MB-231 cells were seeded in treated 96 well plates at 37°C until reaching ∼80-90% confluency. Cells were washed thrice with 1X PBS, then incubated with varying concentrations of electrophilic compounds in DMEM containing no FBS for 1 h. Cells were again washed thrice, then incubated with 100 µM of probe (IA alkyne for cysteine reactivity, 4-pentynoic acid succinimidyl ester for lysine reactivity) in DMEM containing no FBS for 1 h. Cells were washed thrice, removed using TrypLE^TM^ Express Enzyme (Thermo Fisher), and transferred to a round-bottom 96-well plate. Transferred cells were centrifuged (1000 x g, 5 min) in a Thermo Scientific Jouan C4i centrifuge, and the cell pellets were fixed in 4% formaldehyde solution for 20 minutes. The plate was centrifuged (1000 x g, 5 min) and pelleted cells were resuspended with 20 mM coumarin azide with 1 mM CuSO_4_, 128 mM THPTA, and 1.2 mM ascorbic acid in a solution of 0.2% saponin in PBS (in order to allow permeabilization of the fluorophore) for 30 min. Cells were analyzed using an Attune NxT Flow Cytometer (Thermo Fisher) equipped with a 405 nm laser with 440/50 nm bandpass filter.

#### Competitive In-Gel Fluorescence Analysis of Proteome-Wide Reactivity

MDA-MB-231 cells were seeded in treated 6 well plates at 37°C until reaching 100% confluency. Cells were washed thrice with 1X PBS, scraped to remove, and centrifuged (1.5 x g, 5 min) to pellet. After removing the supernatant, the harvested cell pellets were then lysed by sonication in PBS pH 7.4 containing 1 mM EDTA and 1% Triton X-100. Cell lysates were incubated with 100 µM of electrophilic compounds for 30 min at 37°C, then 100 µM of probe (IA alkyne for cysteine reactivity, 4-pentynoic acid succinimidyl ester for lysine reactivity) for 30 min at 37°C. Cell lysates were subsequently reacted with 150 mM FAM-N_3_ with 1 mM CuSO_4_, 128 mM THPTA, and 1.2 mM ascorbic acid for 30 min at 37°C. Reactions were resolved by 12% SDS-PAGE (200 V, 1 h) and scanned for fluorescence using an Azure Biosystems 300Q Imaging System.

#### Competitive In-Gel Fluorescence Analysis of Cefoxitin Reactivity in Human Serum

Human serum was diluted to 0.5 mg/mL in PBS and incubated with 200 µM cefoxitin for 30 min at 37°C, then indicated concentrations (either 10 µM or 1 µM) of 4-pentynoic acid succinimidyl ester for 30 min at 37°C. Serum was subsequently reacted with 20 mM FAM-N_3_ with 1 mM CuSO_4_, 128 mM THPTA, and 1.2 mM ascorbic acid for 30 min at 37°C. Reactions were resolved by 12% SDS-PAGE (200 V, 1 h) and scanned for fluorescence using an Azure Biosystems 300Q Imaging System.

#### In-Gel Fluorescence Analysis of Azidocillin-Modified HSA in Human Serum

Human serum was diluted to 0.5 mg/mL in PBS and incubated with 200 µM of azidocillin for 30 min at 37°C. Serum was subsequently reacted with 200 µM FAM-alkyne with 1 mM CuSO_4_, 128 mM THPTA, and 1.2 mM ascorbic acid for 30 min at 37°C. Reactions were resolved by 12% SDS-PAGE (200 V, 1 h) and scanned for fluorescence using an Azure Biosystems 300Q Imaging System.

#### MGO Modification of ovaK7R and Creatine Scavenging Reactions

Glycation reactions with MGO were carried out as previously described,^51^ with minor modifications. Reactions were carried out in Eppendorf tubes with a final volume of 150 µL. For reactions only containing peptide and MGO, 50 µM **ovaK7R** in DMF and 50 equiv. (2.5 mM) MGO in water were added, along with 30 µL of 10X PBS and the remaining volume with water, and allowed to shake at 37°C for 24 h. For creatine “scavenge” reactions, 10 equiv. creatine (50 mM) was incubated with MGO (5 mM) for 2 h at 37°C brought to a final volume of 75 µL with water, after which time 50 µM **ovaK7R** in DMF, 30 µL of 10X PBS, and the remaining volume to 150 µL total was added. This reaction proceeded at 37°C for 24 h as before. Reactions were diluted 500x for B3Z assay (see “Co-Incubation with RMA-S cells for MGO-modified **ovaK7R** reaction treatment” section above under “B3Z T Cell Activation Assays”).

#### Western Blotting

MDA-MB-231 cells were seeded in a treated 6 well plate at 37°C until reaching ∼80-90% confluency. Cells were washed with 1X PBS, then incubated with 800 µM MGO, 10 mM creatine, or both, in culture media (with no FBS) at 37°C for 2 h. Cells were washed thrice, scraped to remove, and centrifuged (1.5 x g, 5 min) to pellet. After removing the supernatant, cell pellets were then lysed on ice with RIPA buffer. Cell lysates were normalized to 1 mg/mL (BCA protein assay), followed by the addition of 4X SDS gel loading buffer and boiling for 5 min. Proteins were resolved by 12% SDS-PAGE (200 V, 1 h). Proteins were then blotted onto a nitrocellulose membrane (100 V, 1 h) and stained with Ponceau S staining solution. After destaining, membranes blocked with 5% milk in Tris-buffered saline with Tween-20 (TBST) for 1 h at room temperature, then incubated with a 1:900 dilution of MG-H1 monoclonal antibody (clone 1H7G5) at 4°C overnight. Membranes were washed three times with TBST before treatment with a 1:1000 dilution of secondary antibody (HRP goat anti-mouse IgG) in 5% milk in TBST for 1 h at room temperature. Membranes were washed five times with TBST, then developed with Western Blotting Luminol Reagent (Santa Cruz Biotechnology) and visualized by chemiluminescence scan on a BioRad ChemiDoc XRS+ Gel Imaging System with Image Lab Software.

#### Immunoaffinity Pulldown of MHC-I-Bound Peptides Acylated with 1

Immunoaffinity purification was performed as described previously.^41^ First, NHS-Sepharose beads were washed twice with PBS. W6/32 antibody was conjugated to NHS-Sepharose beads at a ratio of 3 mg of antibody to 300 mL of NHS-Sepharose bead solution by rotating overnight at 4°C in 0.2 M NaHCO_3_, 0.5 M NaCl pH 8.3. Beads were then quenched in a solution of 100 mM Tris pH 8.0 by rotating for 1 h at 4°C. Beads were washed twice with alternating solutions of 100 mM ammonium acetate, 500 mM NaCl pH 5.0; and 100 mM Tris pH 8.0. If necessary, during subsequent preparation of cell lysates, beads were stored in 20 mM Tris, 150 mM NaCl (include 0.02% sodium azide if long-term) at 4°C. Next, 2 x 10^8^ MDA-MB-231 cells (that had been incubated with 800 µM of **1** overnight) were thoroughly washed with 1X PBS pH 7.4. Cells were centrifuged (1000 x g, 5 min) and the cell pellet was lysed on ice in lysis buffer (50 mM Tris, 150 mM NaCl, 20 mM iodoacetamide, 1% CHAPS, 1 protease inhibitor tablet, pH 7.4), 2 mL lysis buffer per 100 million cells. Lysed cells were then centrifuged (10,000 x g for 1 h at 4°C). The supernatant was then rotated with W6/32-conjugated beads at 4°C overnight. Beads were washed with the 10 column volumes (CV) each of the following series of buffers: 50 mM Tris, 150 mM NaCl pH 8.0; 50 mM Tris, 450 mM NaCl pH 8.0; 50 mM Tris, 150 mM NaCl pH 8.0; 50 mM Tris pH 8.0. Peptides were eluted with 500 mL of a 10% acetic acid solution, and this elution step was repeated to obtain all eluted peptides fractions. The solution was then lyophilized overnight to obtain a dry powder. The lyophilized sample was reacted with 100 µM DADPS-biotin azide in 1.2 mM sodium ascorbate, 1 mM CuSO_4_, and 100 µM TBTA at pH 7.4 in 1X PBS at 4°C rotating overnight. The following day, sodium ascorbate was replenished with the addition of another 1.2 mM, and the solution was allowed to continue incubating for another day. The reaction mixture was spun through a 3 kDa Amicon Ultra centrifugal filter column, and the filtrate was lyophilized overnight. Finally, the lyophilized powder was resuspended in 1X PBS pH 7.4 and loaded onto an Oasis HLB column following manufacturer’s protocol and eluted with 60% acetonitrile. The flowthrough was lyophilized and stored at -80°C until further analysis.

#### Mass Spectrometry Data Acquisition

Data were acquired using an Orbitrap Astral mass spectrometer in a DDA mode. For MS1 scans, the Orbitrap resolution was set at 120,000 with an AGC target of 100%. MS1 spectra were recorded over an m/z range of 350–1350, with a maximum injection time of 50 ms. The isolation window for MS2 precursor selection was set to 1.2 m/z. Up to 30 scans were acquired per MS1 cycle for precursor ions with intensities greater than 5.0 x 10^3^ and charge states ranging from 2 to 6. MS2 fragmentation was performed with HCD at 25% collision energy and a maximum injection time of 25 ms. Dynamic exclusion was turned on with a duration of 20 seconds.

## SYNTHESIS AND CHARACTERIZATION

### General procedure for the solid-phase synthesis of peptides

All peptides were prepared by standard Fmoc-based solid-phase chemistry using the appropriate resins. Briefly, to a 25 mL peptide synthesis vessel, an appropriate amount of resin was added, followed by 20% piperidine in N, N-Dimethylformamide (DMF, 15 mL) (if resin was Fmoc-protected). Briefly, to a 25 mL peptide synthesis vessel, an appropriate amount of resin was added, followed by 20% piperidine in N, N-Dimethylformamide (DMF, 15 mL) (if resin was Fmoc-protected). This was followed by shaking at room temperature for 30 min. The resin was then washed with methanol (CH_3_OH) and dichloromethane (DCM) three times. After the last wash, 4 equiv. of amino acid was added along with 4 equiv. of ethyl cyanohydroxyiminoacetate (Oxyma) and 4 equiv. of N, N’-Diisopropylcarbodiimide (DIC). The resin was shaken at room temperature for 2 h, then washed with CH_3_OH/DCM. The remainder of the amino acids were coupled in the same manner. Peptides were cleaved from the resin using a TFA/TIPS/H_2_O mixture (95:2.5:2.5, v/v/v) shaking at room temperature for 2 h. The solution was filtered and concentrated prior to precipitation by the addition of cold diethyl ether to yield crude peptide. Crude peptides were purified by reverse-phase preparative high-performance liquid chromatography (RP-HPLC) equipped with Waters 1525 with 2489 UV/Visible Detector on a Phenomenex Luna 10 μm C8(2) 100 Å (250 x 21.2 mm) column using gradient elution with H_2_O/CH_3_OH with 0.1% TFA at 10 mL/min. The HPLC fractions of the desired purified compounds were first concentrated under reduced pressure using a rotary evaporator. The final concentrated aqueous solutions were lyophilized to dryness using Labconco Freezone 4.5 L (-84°C) lyophilizer. The peptides were analyzed for purity using Phenomenex Luna 5 µm C8(2) on the same RP-HPLC; gradient elution in H_2_O/CH_3_OH with 0.1% TFA at 1 mL/min. Peptide identities were analyzed using high resolution electrospray ionization mass spectrometry (HRMS, ESI/MS) analyses obtained on an Agilent 6545B Q-TOF LC/MS equipped with 1260 infinity II LC system with auto sampler, or matrix-assisted laser desorption ionization time-of-flight (MALDI-TOF) mass spectrometry (Shimadzu 8020).

### Solid-phase synthesis of modified peptides

Synthesis was performed in accordance with established procedures for the resin, following the general procedure above (4 equiv. amino acid, 4 equiv. Oxyma, 4 equiv. DIC in DMF, shaking at room temperature for 2 h), but using respective commercially available unnatural amino acids. Fmoc-L-a-aminoadipic acid d-tert-butyl for **ovaK7_aad_**, Fmoc-L-homocitrulline-OH for **ovaK7_hcit_**, Fmoc-*S*-sulfo-L-cysteine disodium for **ovaE6C_SO3_** (Chem Impex catalog #11179), Fmoc-3-nitro-L-tyrosine (Chem Impex catalog #04992) for **ovaF5Y_NO2_**, Fmoc-3-chloro-L-tyrosine (Chem Impex catalog # 15121) for **ovaF5Y_Cl_**, Fmoc-L-methionine sulfoxide (Chem Impex catalog # 03724) for **ca1_oxide_** and **ca2_oxide_**, Fmoc-L-methionine sulfone (Chem Impex catalog # 03725) for **ca1_oxone_** and c**a2_oxone_**, and Fmoc-acetyl-L-lysine (Chem Impex catalog # 02430) for **ovaK7_ASA_**. For other lysine-modified variants, Fmoc-L-lysine(Mtt)-OH was coupled in the seventh position of SIINFEKL in same manner as above (4 equiv. of amino acid along with 4 equiv. of Oxyma and 4 equiv. of DIC) in DMF. The resin was shaken at room temperature for 2 h, then washed with CH_3_OH/DCM, and the remainder of the amino acids were coupled in the same manner. The MTT protecting group was removed by the addition of 1% TFA, 2.5% TIPS, in 10 mL DCM for 15 min, washed and repeated 5 more times. For coupling onto the deprotected lysine residue, 4 equiv. of the respective chemical, 4 equiv. of Oxyma, and 4 equiv. of DIC were added in DMF to the reaction vessel and coupled at room temperature for 2 h. For the following, 4 equiv. of the respective chemical and 4 equiv. of DIEA were added: 4-nitrophenyl formate (Thermo Scientific # L00316.06) for **ovaK7_formyl_**, chlorothalonil for **ovaK7_CTN_**, phenethyl isothiocyanate for **ovaK7_PEITC_**, and allyl isothiocyanate for **ovaK7_AITC_**. Following coupling of all amino acids and modifications, the final Fmoc-deprotection was performed, and cleavage and purification were carried out as described before.

### Solution-phase modification of ovaE6C_2sc_

Synthesis of the parent **ovaE6C** was performed in accordance with the solid-phase synthesis protocol above. Following cleavage from the resin and precipitation in diethyl ether, 5 mM of crude **ovaE6C** was stirred with 2 equiv. of Tris(2-carboxyethyl)phosphine) (TCEP) in 1X PBS (pH adjusted to 8.0) for 30 min at room temperature in a round-bottom flask. 5 equiv. of maleic acid (Sigma-Aldrich catalog #M0375) was then added, and the reaction mixture continued to stir at room temperature for 24 h. The reaction mixture was then purified by reverse-phase preparative high-performance liquid chromatography (RP-HPLC) as described before.

### Solution-phase modification of ovaK7C_PEITC_

Synthesis of the parent **ovaK7C** was performed in accordance with the solid-phase synthesis protocol above. Following cleavage from the resin and precipitation in diethyl ether, 5 mM of crude **ovaK7C** was stirred with 2 equiv. of Tris(2-carboxyethyl)phosphine) (TCEP) in 1X PBS (pH adjusted to 8.0) for 30 min at room temperature in a round-bottom flask. 2 equiv. of 2-phenylethyl isothiocyanate (AK Scientific catalog #J53261) was then added (along with DMF and DMSO added dropwise for solubility) and the reaction mixture continued to stir at room temperature for 4 h. The reaction mixture was then purified by reverse-phase preparative high-performance liquid chromatography (RP-HPLC) as described before.

### Solution-phase modification of ovaK7C_BrEA_ and ovaK7C_CMQ_

Synthesis of the parent **ovaK7C** was performed in accordance with the solid-phase synthesis protocol above. Following cleavage from the resin and precipitation in diethyl ether, 5 mM of crude **ovaK7C** was stirred with 10 equiv. of DL-Dithiothreitol (DTT) in 0.1 M sodium bicarbonate pH 8.4 (along with DMF dropwise for solubility) for 1 h at 37°C in a round-bottom flask. 100 equiv. of 2-bromoethylamine (AK Scientific catalog #J91302) for **ovaK7C_BrEA_** or chlormequat (AK Scientific catalog #J51995) for ovaK7C**_CMQ_** were then added and the reaction mixture continued to stir at 37°C for 3 h. 10 equiv. more of DTT was added and stirred for 30 min at room temperature, followed by 100 equiv. more of bromoethylamine or chlormequat added and stirred overnight (∼16 h) at room temperature. The reaction mixtures were then purified by reverse-phase preparative high-performance liquid chromatography (RP-HPLC) as described before.

### Characterization of Peptides

The peptides were analyzed for purity using Phenomenex Luna 5 µm C8(2) on the same RP-HPLC; gradient elution in H_2_O/CH_3_CN with 0.1% TFA at 1 mL/min. Peptides were analyzed via high resolution electrospray ionization mass spectrometry (HRMS, ESI/MS) on an Agilent 6545B Q-TOF LC/MS equipped with 1260 infinity II LC system with auto sampler, or matrix-assisted laser desorption ionization time-of-flight (MALDI-TOF) mass spectrometry (Shimadzu 8020). All peptides exhibited a purity greater than 93%. Characterization data are provided below.

## ACKNOWLEDGEMENTS

This study was supported by the NIH grant R35GM124893 (M.M.P.).

## SUPPLEMENTARY INFORMATION

Additional figures, tables, and materials/methods are included in the supporting information file.

