## Supplementary Information for "Chemical History Shapes the Antigenic Identity of MHC-I Epitopes"

### Table of Contents


| SUPPLEMENTARY FIGURES | S5 – S24 |
|---|---|
| **Figure S1.** Non-enzymatic PTMs on cancer-associated peptides | S5 |
| **Figure S2.** Analytical HPLC of **ca4** and **ca4**$_{\mathbf{cystine}}$ in cell culture media | S6 |
| **Figure S3.** Analytical HPLC of **ca5** and **ca5**$_{\mathbf{cystine}}$ in cell culture media | S7 |
| **Figure S4.** Uncropped in-gel fluorescence analysis of concentration-dependent labeling of proteins in nonenzymatic acylation probe **1**-treated cells | S8 |
| **Figure S5.** Uncropped in-gel fluorescence analysis of proteome-wide interactions of environmental chemicals using lysine-reactive probe | S9 |
| **Figure S6.** Flow cytometry analysis of proteome-wide interactions of environmental chemicals using lysine-reactive probe | S10 |
| **Figure S7.** Uncropped in-gel fluorescence analysis of proteome-wide interactions of environmental chemicals using 100 μM cysteine-reactive probe | S11 |
| **Figure S8.** Uncropped in-gel fluorescence analysis of proteome-wide interactions of environmental chemicals using 50 μM cysteine-reactive probe | S12 |
| **Figure S9.** Uncropped in-gel fluorescence analysis of proteome-wide interactions of chlorothalonil and picloram (concentration scan) using cysteine-reactive probe | S13 |
| **Figure S10.** Flow cytometry analysis of proteome-wide interactions of environmental chemicals using cysteine-reactive probe | S14 |
| **Figure S11.** Uncropped in-gel fluorescence analysis of proteome-wide interactions of ASA and isothiocyanates using lysine-reactive probe | S15 |
| **Figure S12.** Flow cytometry analysis of proteome-wide interactions of ASA and isothiocyanates using lysine-reactive probe | S16 |
| **Figure S13.** Uncropped in-gel fluorescence analysis of proteome-wide interactions of ASA and isothiocyanates using cysteine-reactive probe | S17 |
| **Figure S14.** Flow cytometry analysis of proteome-wide interactions of ASA and isothiocyanates using cysteine-reactive probe | S18 |
| **Figure S15.** In-gel fluorescence analysis of cefoxitin reactivity using lysine-reactive probe in human serum | S19 |
| **Figure S16.** In-gel fluorescence analysis of azidocillin-modified HSA in human serum | S20 |

| | |
|---|---|
| **Figure S17.** Uncropped in-gel fluorescence analyses of β-lactam interactions in human serum (**Figs. S15, S16** uncropped) | S21 |
| **Figure S18**. Ion chromatogram for (M+54) adduct of MGO modification reaction with **ovaK7R** | S22 |
| **Figure S19**. Ion chromatogram for (M+72) and (M+144) adducts of MGO modification reaction with **ovaK7R** | S23 |
| **Figure S20.** Uncropped Western blot analysis of MGO-treated cells with creatine scavenging | S24 |
| **MATERIALS AND METHODS** | **S25 – S33** |
| **Materials** | S25 – S27 |
| **Experimental Methods** | S28 – S33 |
| Mammalian Cell Culture | S28 |
| RMA-S Stabilization Assay | S28 |
| B3Z T Cell Activation Assays | S28 |
| Flow Cytometry Analysis of Non-Enzymatic Acylation with **1** | S30 |
| In-Gel Fluorescence Analysis of Non-Enzymatic Acylation with **1** | S30 |
| Competitive Flow Cytometry Analysis of Proteome-Wide Reactivity | S30 |
| Competitive In-Gel Fluorescence Analysis of Proteome-Wide Reactivity | S31 |
| Competitive In-Gel Fluorescence Analysis of Cefoxitin Reactivity in Human Serum | S31 |
| In-Gel Fluorescence Analysis of Azidocillin-Modified HSA in Human Serum | S31 |
| MGO Modification of **ovaK7R** and Creatine Scavenging Reactions | S31 |
| Western Blotting | S32 |
| Immunoaffinity Pulldown of MHC-I-Bound Peptides | S32 |
| Mass Spectrometry Data Acquisition | S33 |
| **SYNTHESIS AND CHARACTERIZATION** | **S34 – S36** |
| General procedure for solid-phase synthesis of peptides | S34 |
| Solid-phase synthesis of modified peptides | S34 |
| Solution-phase modification of **ovaE6C$_{2sc}$** | S35 |
| Solution-phase modification of **ovaK7C$_{PEITC}$** | S35 |
| Solution-phase modification of **ovaK7C$_{BrEA}$** | S36 |
| **Characterization of Peptides** | S37 – S75 |
| **ovaWT** (SIINFEKL) | S38 |
| **cntPEP** (SNFVSAGI) | S39 |
| **ovaK7$_{formyl}$** | S40 |
| **ovaK7$_{aad}$** | S41 |
| **ovaK7$_{hcit}$** | S42 |
| **ovaK7$_{lac}$** | S43 |
| **ovaK7$_{suc}$** | S44 |

| | |
|---|---|
| **ovaE6C** (SIINFCKL) | S45 |
| **ovaE6C$_{SO3}$** | S46 |
| **ovaE6C$_{2sc}$** | S47 |
| **ovaN4D** (SIIDFEKL) | S48 |
| **ovaN4D$_{iso}$** | S49 |
| **ovaF5Y** (SIINYEKL) | S50 |
| **ovaF5Y$_{NO2}$** | S51 |
| **ovaF5Y$_{Cl}$** | S52 |
| **ovaK7$_{CTN}$** | S53 |
| **ovaK7$_{ASA}$** | S54 |
| **ovaK7$_{CFX}$** | S55 |
| **ovaK7$_{PEITC}$** | S56 |
| **ovaK7$_{AITC}$** | S57 |
| **ovaK7C** (SIINFECL) | S58 |
| **ovaK7C$_{PEITC}$** | S59 |
| **ovaK7C$_{BrEA}$** | S60 |
| **ovaK7C$_{CMQ}$** | S61 |
| **ovaK7R** (SIINFERL) | S62 |
| **ca1** | S63 |
| **ca1$_{oxide}$** | S64 |
| **ca1$_{oxone}$** | S65 |
| **ca2** | S66 |
| **ca2$_{oxide}$** | S67 |
| **ca2$_{oxone}$** | S68 |
| **ca3** | S69 |
| **ca3$_{asp}$** | S70 |
| **ca3$_{isoasp}$** | S71 |
| **ca4** | S72 |
| **ca4$_{cystine}$** | S73 |
| **ca5** | S74 |
| **ca5$_{cystine}$** | S75 |
| **REFERENCES** | **S76** |

#### SUPPLEMENTARY FIGURES

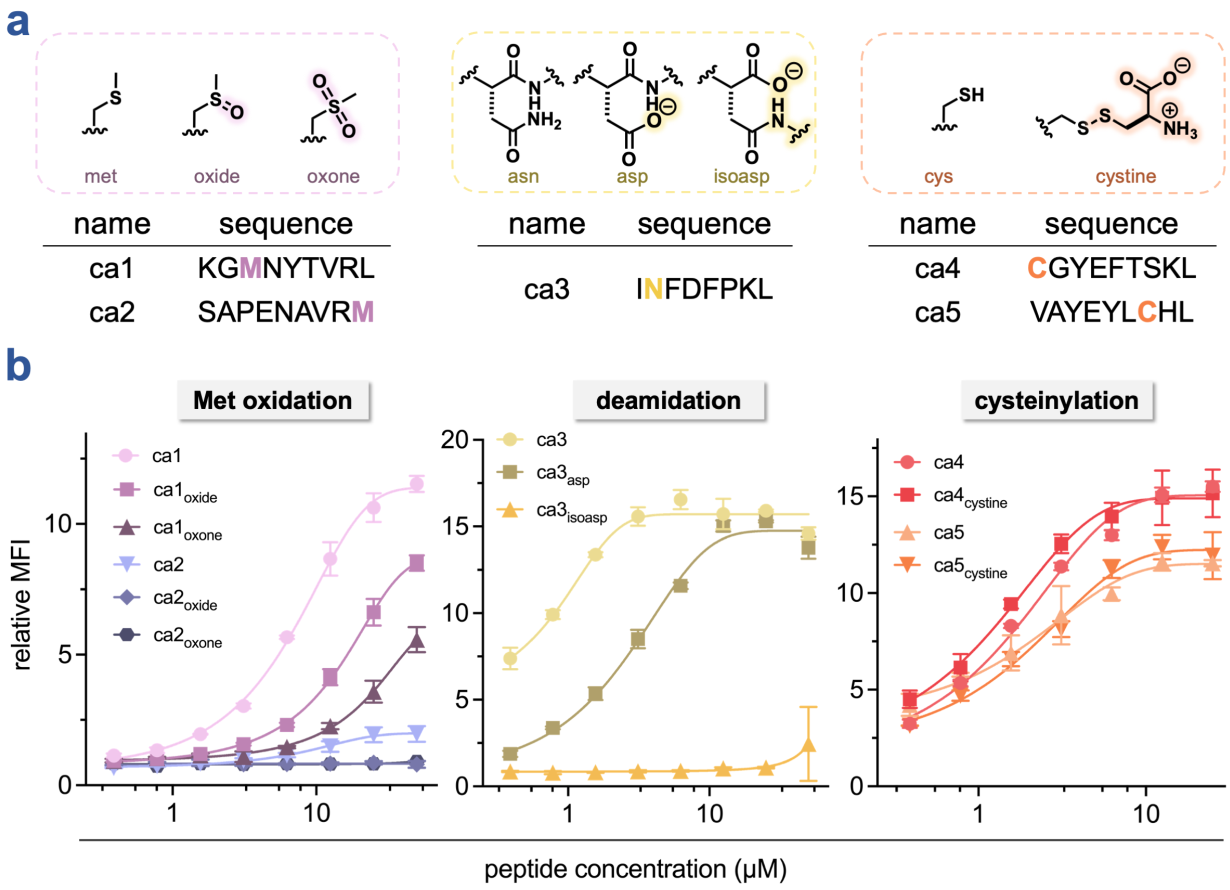


**Figure S1. (a)** Sequences of cancer-associated peptides and respective non-enzymatic PTMs of methionine oxidation (pink), deamidation (yellow), and cysteinylation (orange). **(b)** Dose-response curves from flow cytometry analysis of the RMA-S stabilization assay for non-enzymatic PTM cancer-associated peptides. RMA-S cells were incubated with the indicated concentration of peptides and analyzed via flow cytometry for H-2K$^b$ expression by APC anti-mouse H-2K$^b$ antibody. MFI is the mean fluorescence intensity of the level of fluorescence relative to the negative control peptide (**cntPEP**). Data are represented as mean ± SD (*n* = 3) of technical replicates, and Boltzmann sigmoidal curves were fitted to the data using GraphPad Prism.

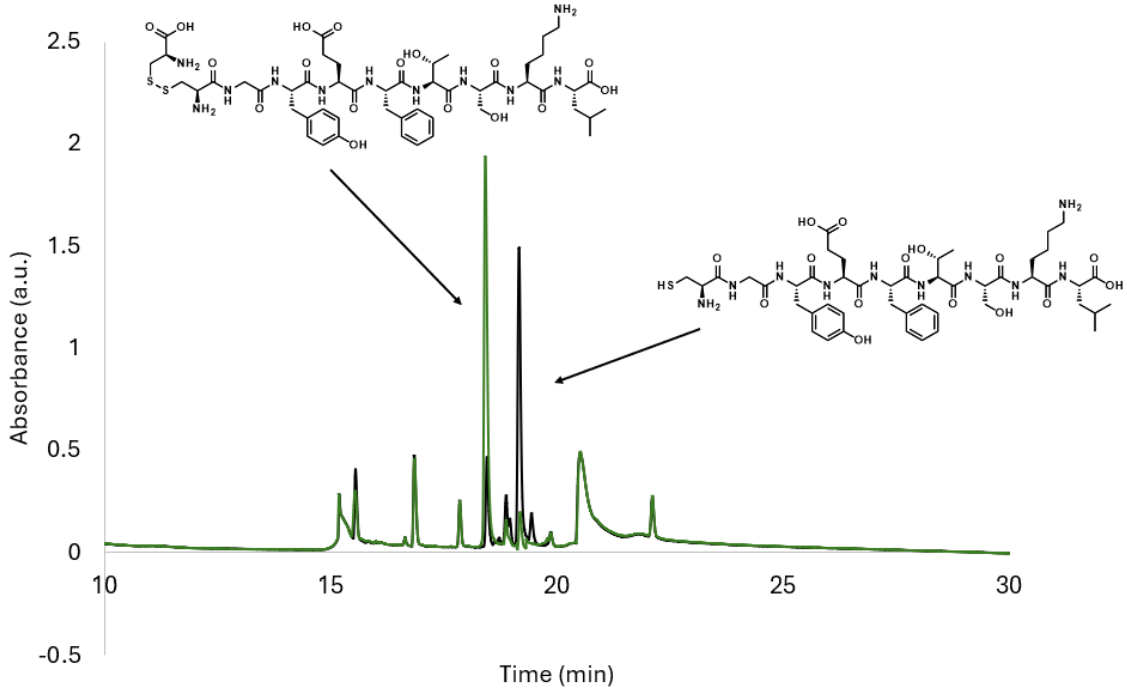


**Figure S2.** Analytical HPLC of **ca4** and **ca4**$_{\textbf{cystine}}$. Peptides were incubated in media containing 10% FBS for 6 hours at 37°C. Overlaid HPLC chromatograms of the mixture are shown for **ca4** (black) and **ca4**$_{\textbf{cystine}}$ (green).

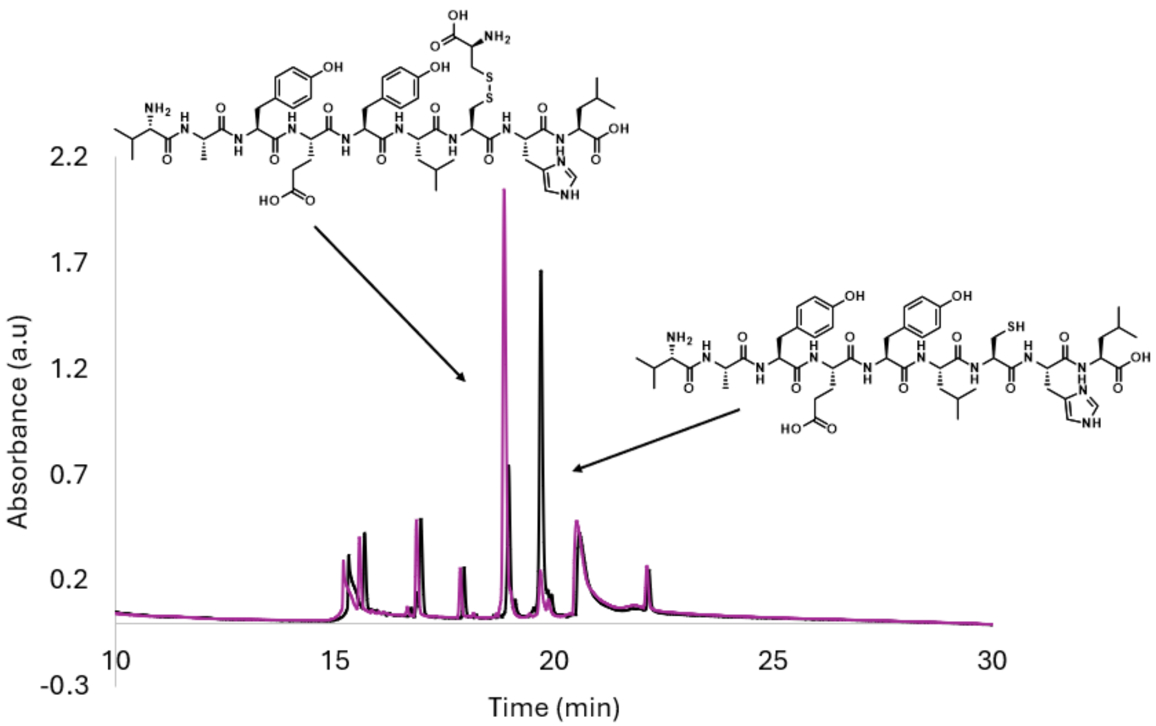


**Figure S3.** Analytical HPLC of **ca5** and **ca5$_{\text{cystine}}$**. Peptides were incubated in media containing 10% FBS for 6 hours at 37$^{o}$C. Overlaid HPLC chromatograms of the mixture are shown for **ca5** (black) and **ca5$_{\text{cystine}}$** (purple).

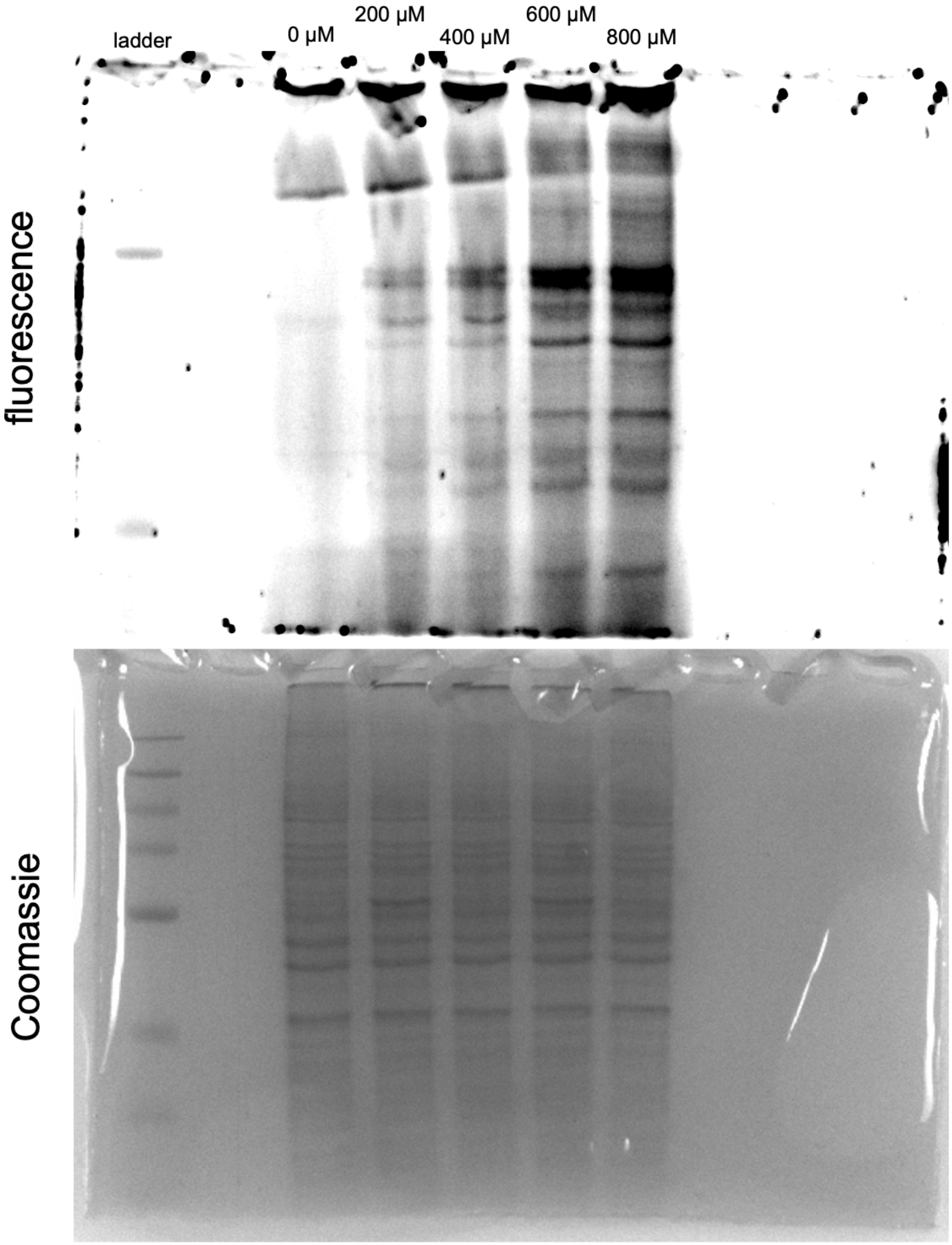


**Figure S4.** Uncropped in-gel fluorescence analysis of concentration-dependent labeling of proteins in nonenzymatic acylation probe **1**-treated cells. MDA-MB-231 cells were treated with varying concentrations of **1**. Cells were then lysed and reacted with FAM-$N_3$ in the presence of $CuSO_4$, ascorbic acid, and THPTA. Acylation sites of probe **1** were resolved by SDS-PAGE and visualized by in-gel fluorescence.

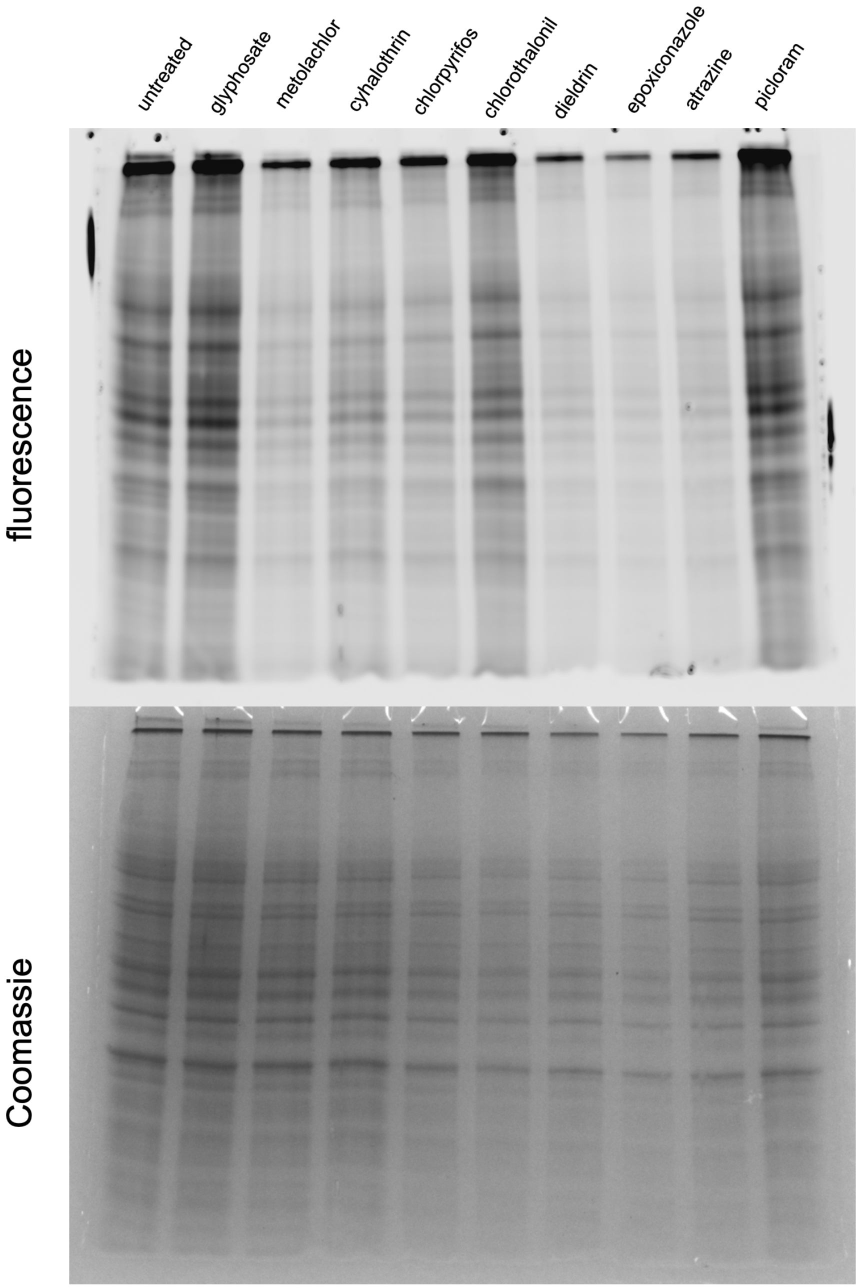


**Figure S5.** Uncropped in-gel fluorescence analysis of proteome-wide interactions of environmental chemicals using lysine-reactive probe. MDA-MB-231 cell lysates were treated with environmental chemicals (200 μM, 30 min) then competed with 4-pentynoic acid succinimidyl ester labeling (200 μM, 30 min), followed by a reaction with FAM-$N_3$ in the presence of $CuSO_4$, ascorbic acid, and THPTA. Lysine reactivity was resolved by SDS-PAGE and visualized by in-gel fluorescence.

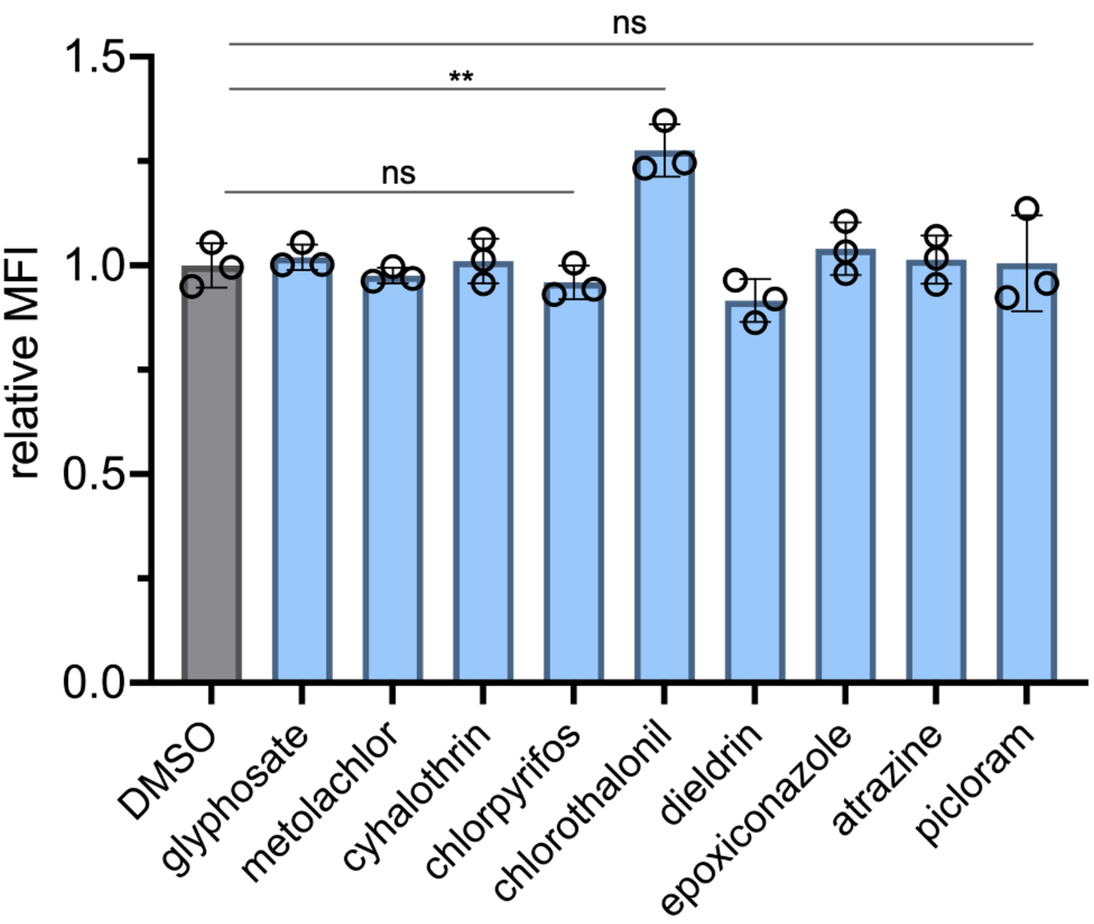


**Figure S6.** Flow cytometry analysis of proteome-wide interactions of environmental chemicals using lysine-reactive probe. MDA-MB-231 cells were treated with environmental chemicals (200 μM, 1 h) then competed with 4-pentynoic acid succinimidyl ester labeling (100 μM, 1 h), followed by a reaction with 3-azido-7-hydroxycoumarin in the presence of $CuSO_4$, ascorbic acid, and THPTA. Lysine reactivity was assessed by flow cytometry, where a decrease in MFI (mean fluorescence intensity) relative to the DMSO control is representative of an increase in reactivity. Data are represented as mean ± SD ($n$ = 3). $P$-values were determined by a two-tailed $t$-test (ns = not significant, ** $p < 0.01$).

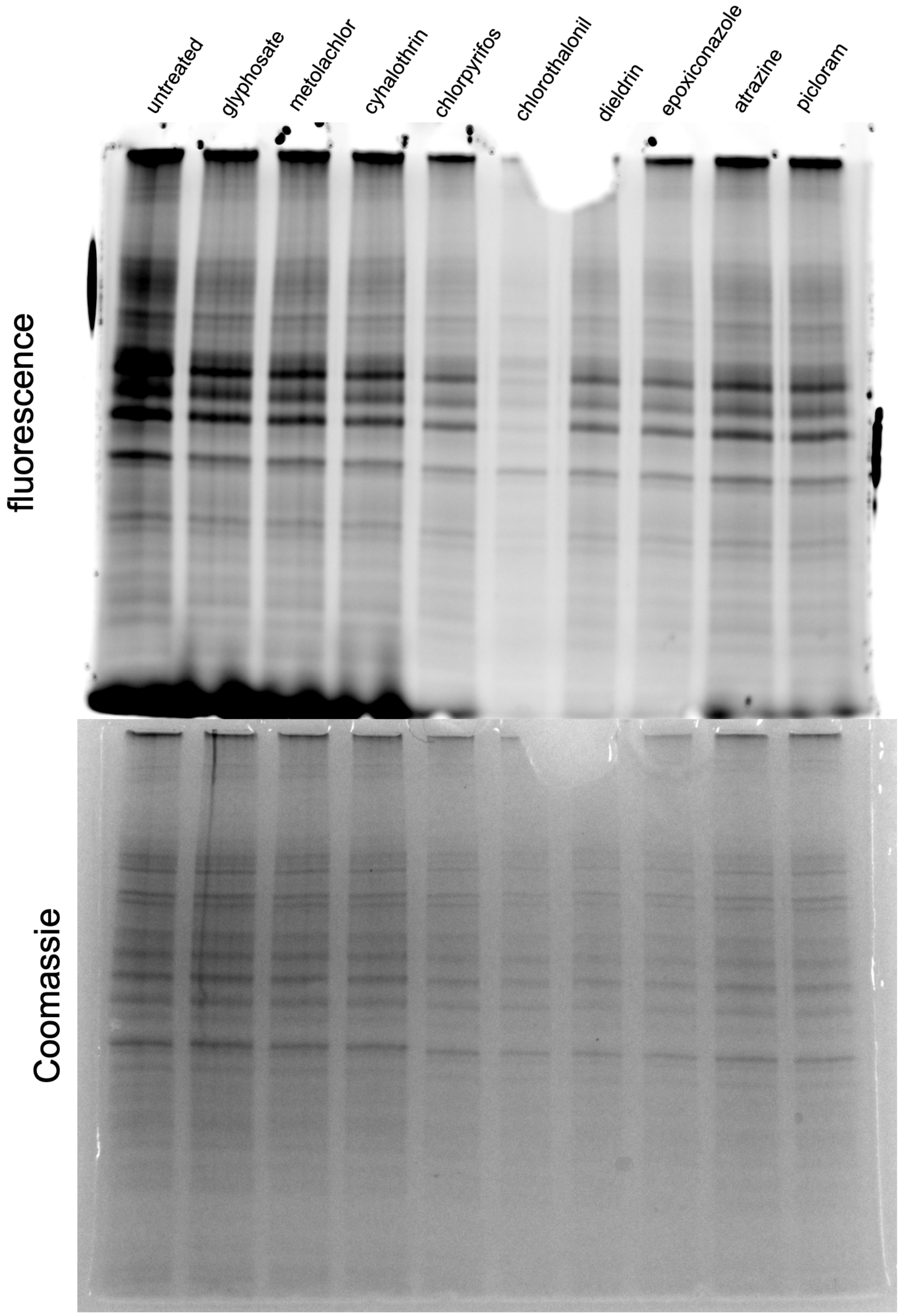


**Figure S7.** Uncropped in-gel fluorescence analysis of proteome-wide interactions of environmental chemicals using 100 µM cysteine-reactive probe. MDA-MB-231 cell lysates were treated with environmental chemicals (100 µM, 30 min), then competed with IA-alkyne labeling (100 µM, 30 min), followed by a reaction with FAM-$N_3$ in the presence of $CuSO_4$, ascorbic acid, and THPTA. Cysteine reactivity was resolved by SDS-PAGE and visualized by in-gel fluorescence.

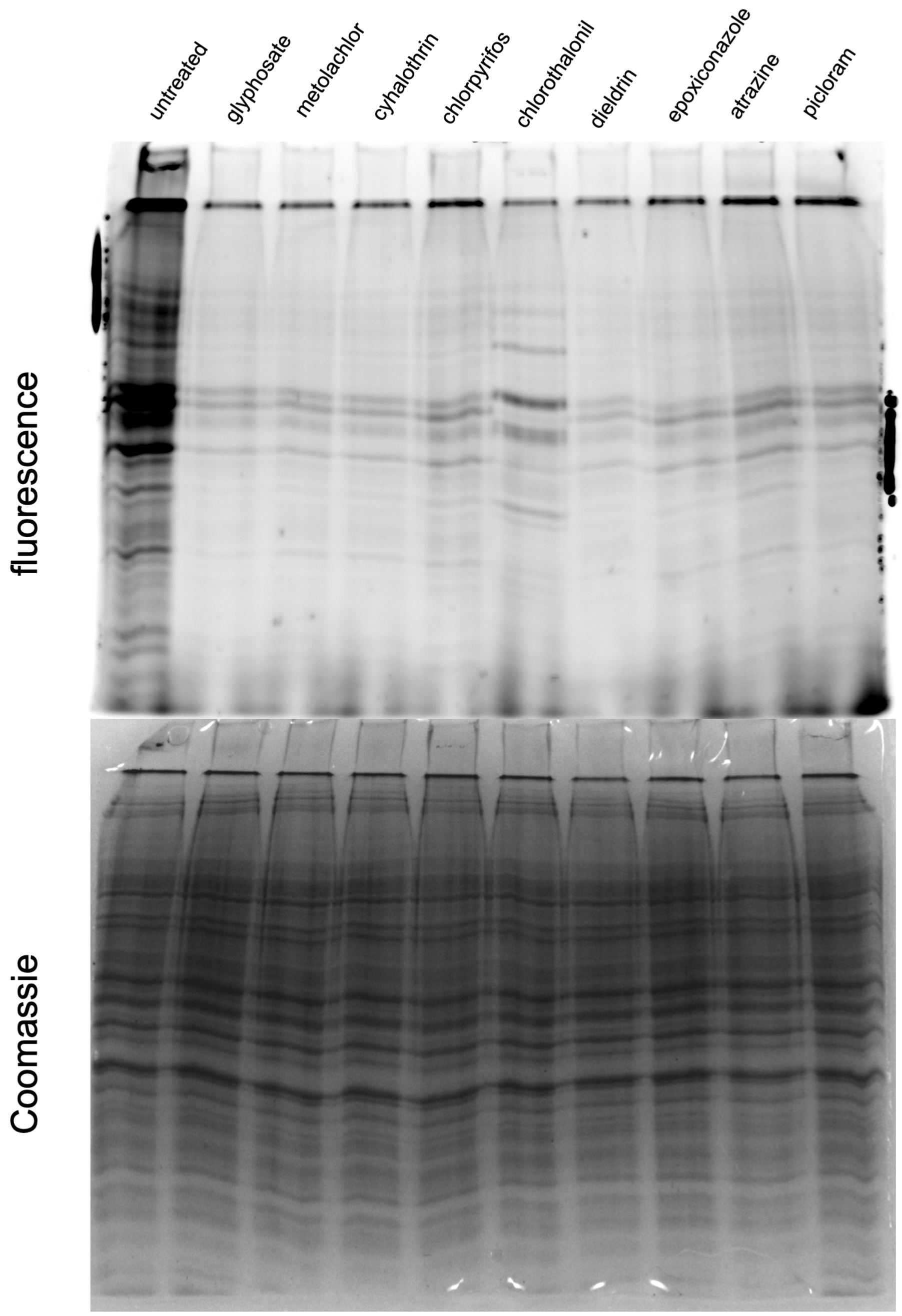


**Figure S8.** Uncropped in-gel fluorescence analysis of proteome-wide interactions of environmental chemicals using 50 µM cysteine-reactive probe. MDA-MB-231 cell lysates were treated with environmental chemicals (100 µM, 30 min), then competed with IA-alkyne labeling (50 µM, 30 min), followed by a reaction with FAM-$N_3$ in the presence of $CuSO_4$, ascorbic acid, and THPTA. Cysteine reactivity was resolved by SDS-PAGE and visualized by in-gel fluorescence.

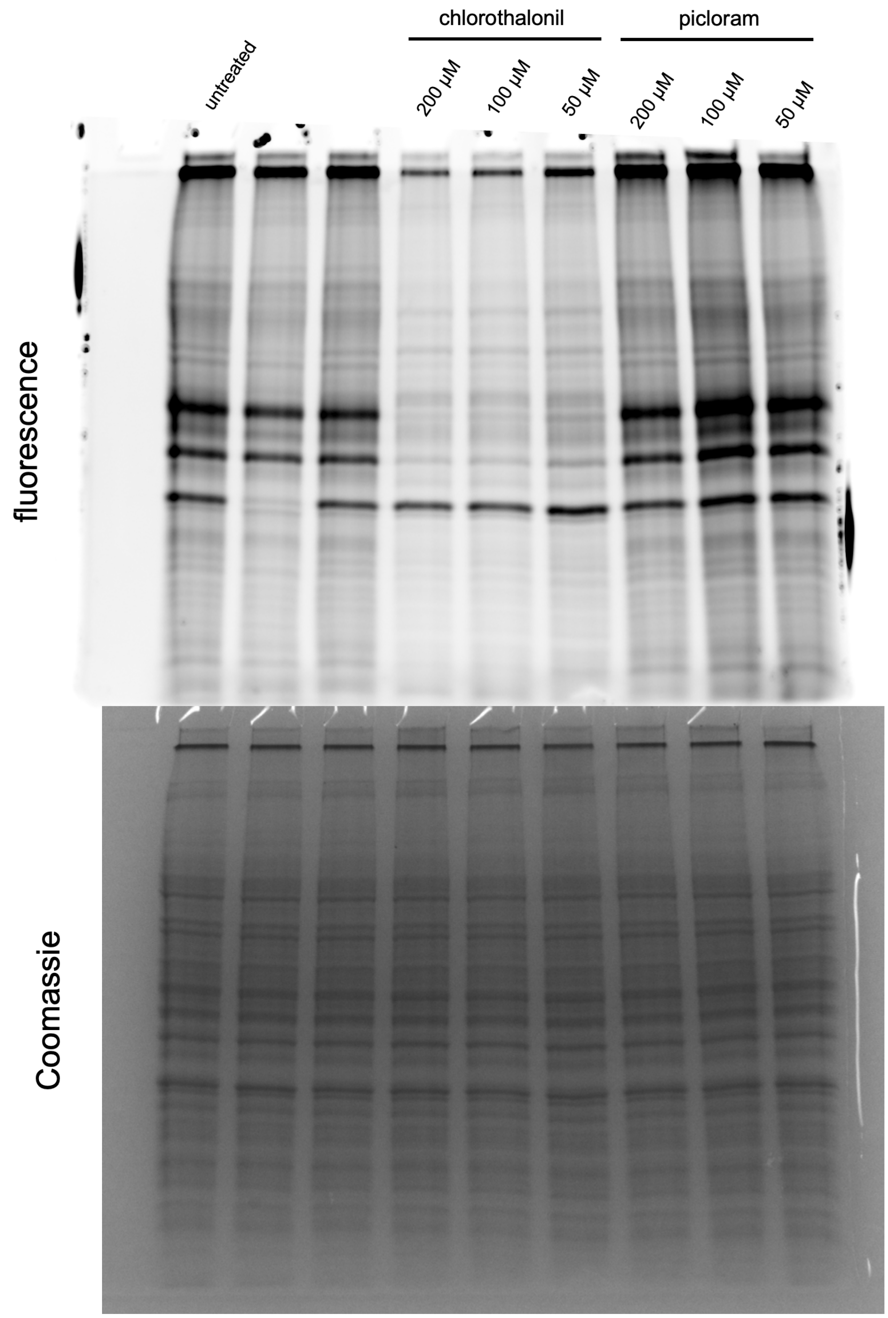


**Figure S9.** Uncropped In-gel fluorescence analysis of proteome-wide interactions of chlorothalonil and picloram (concentration scan) using cysteine-reactive probe. MDA-MB-231 cell lysates were treated with indicated concentrations of chlorothalonil and picloram (30 min), then competed with IA-alkyne labeling (100 µM, 30 min), followed by a reaction with FAM-$N_3$ in the presence of $CuSO_4$, ascorbic acid, and THPTA. Cysteine reactivity was resolved by SDS-PAGE and visualized by in-gel fluorescence.

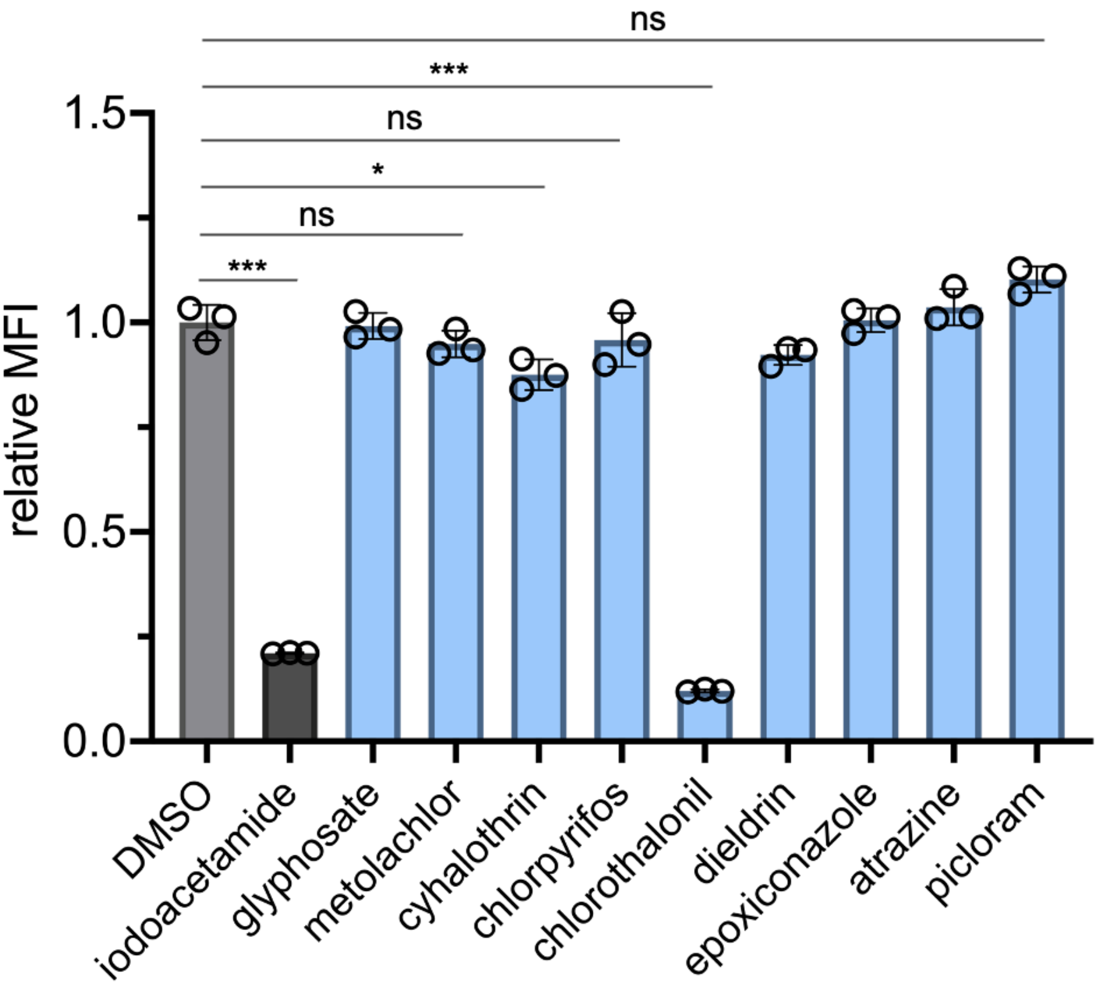


**Figure S10.** Flow cytometry analysis of proteome-wide interactions of environmental chemicals using cysteine-reactive probe. MDA-MB-231 cells were treated with positive control iodoacetamide or environmental chemicals (200 μM, 1 h), then competed with IA-alkyne (100 μM, 1 h), followed by a reaction with 3-azido-7-hydroxycoumarin in the presence of $CuSO_4$, ascorbic acid, and THPTA. Cysteine reactivity was assessed by flow cytometry, where a decrease in MFI (mean fluorescence intensity) relative to the DMSO control is representative of an increase in reactivity. Data are represented as mean ± SD ($n$ = 3). $P$-values were determined by a two-tailed $t$-test (ns = not significant, * $p$ < 0.05, ** $p$ < 0.01).

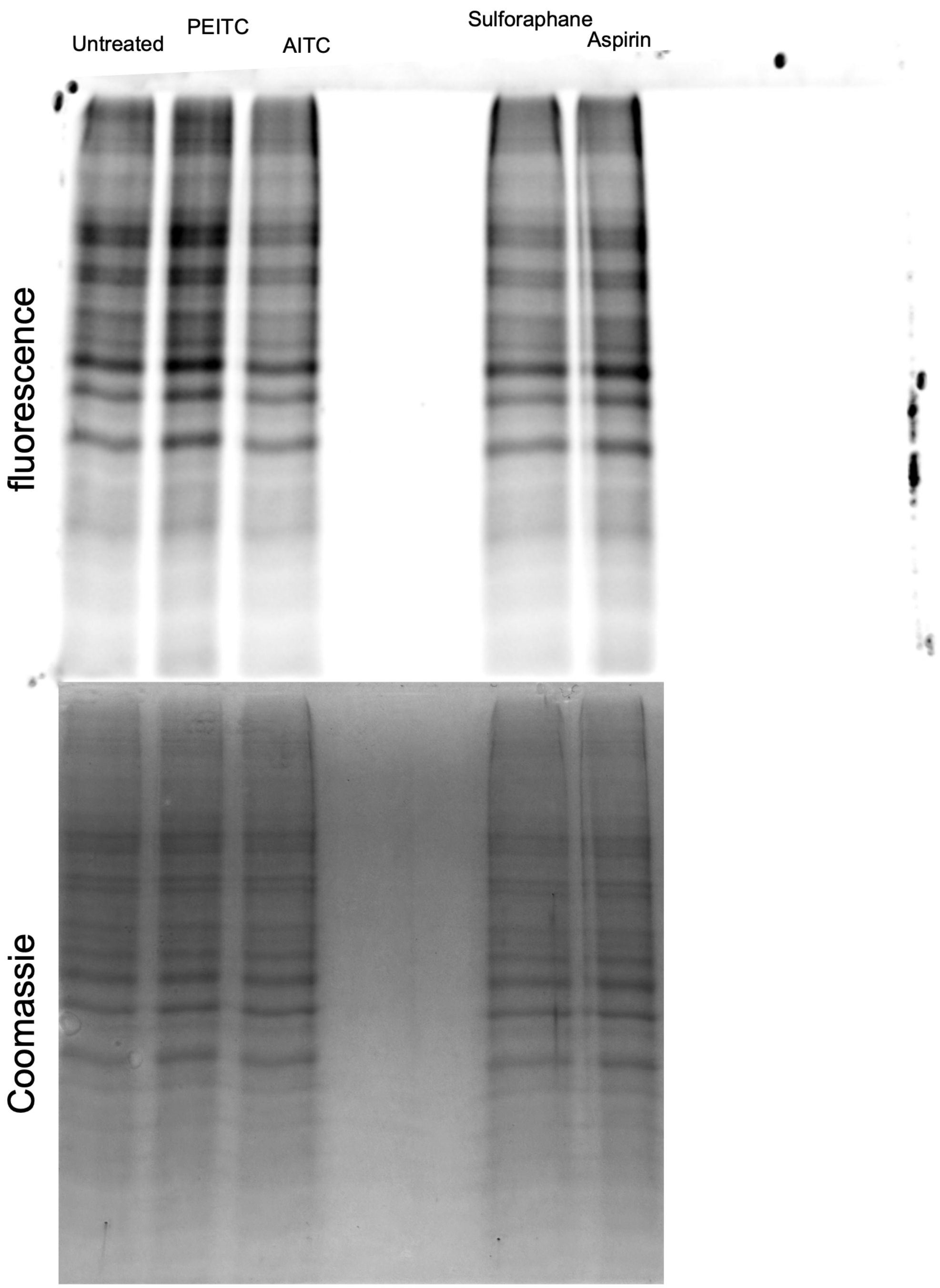


**Figure S11.** Uncropped In-gel fluorescence analysis of proteome-wide interactions of isothiocyanates and aspirin using lysine-reactive probe. MDA-MB-231 cell lysates were treated with isothiocyanates (100 µM, 30 min) or aspirin (1 mM, 30 min) then competed with 4-pentynoic acid succinimidyl ester labeling (100 µM, 30 min), followed by a reaction with FAM-$N_3$ in the presence of $CuSO_4$, ascorbic acid, and THPTA. Lysine reactivity was resolved by SDS-PAGE and visualized by in-gel fluorescence.

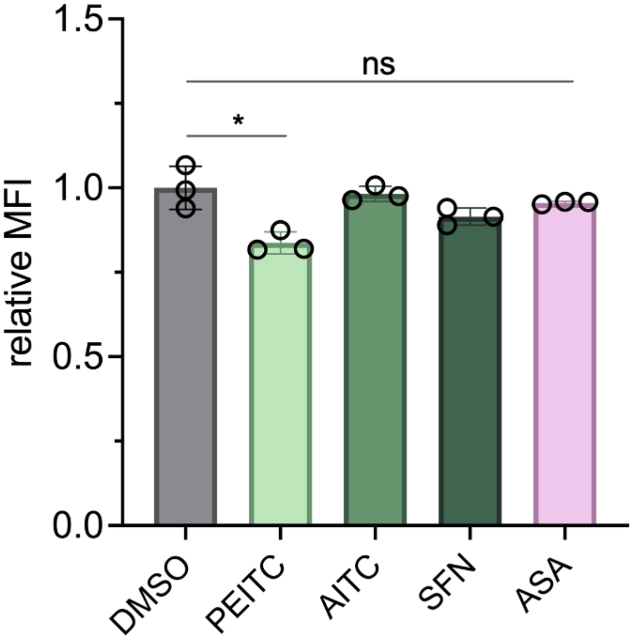


**Figure S12.** Flow cytometry analysis of proteome-wide interactions of isothiocyanates and aspirin using lysine-reactive probe. MDA-MB-231 cells were treated with ITCs (200 μM, 1 h) or aspirin (1 mM, 1 h) then competed with 4-pentynoic acid succinimidyl ester labeling (100 μM, 1 h), followed by a reaction with 3-azido-7-hydroxycoumarin in the presence of $CuSO_4$, ascorbic acid, and THPTA. Lysine reactivity was assessed by flow cytometry, where a decrease in MFI (mean fluorescence intensity) relative to the DMSO control is representative of an increase in reactivity. Data are represented as mean ± SD ($n$ = 3). $P$-values were determined by a two-tailed $t$-test (ns = not significant, * $p < 0.05$).

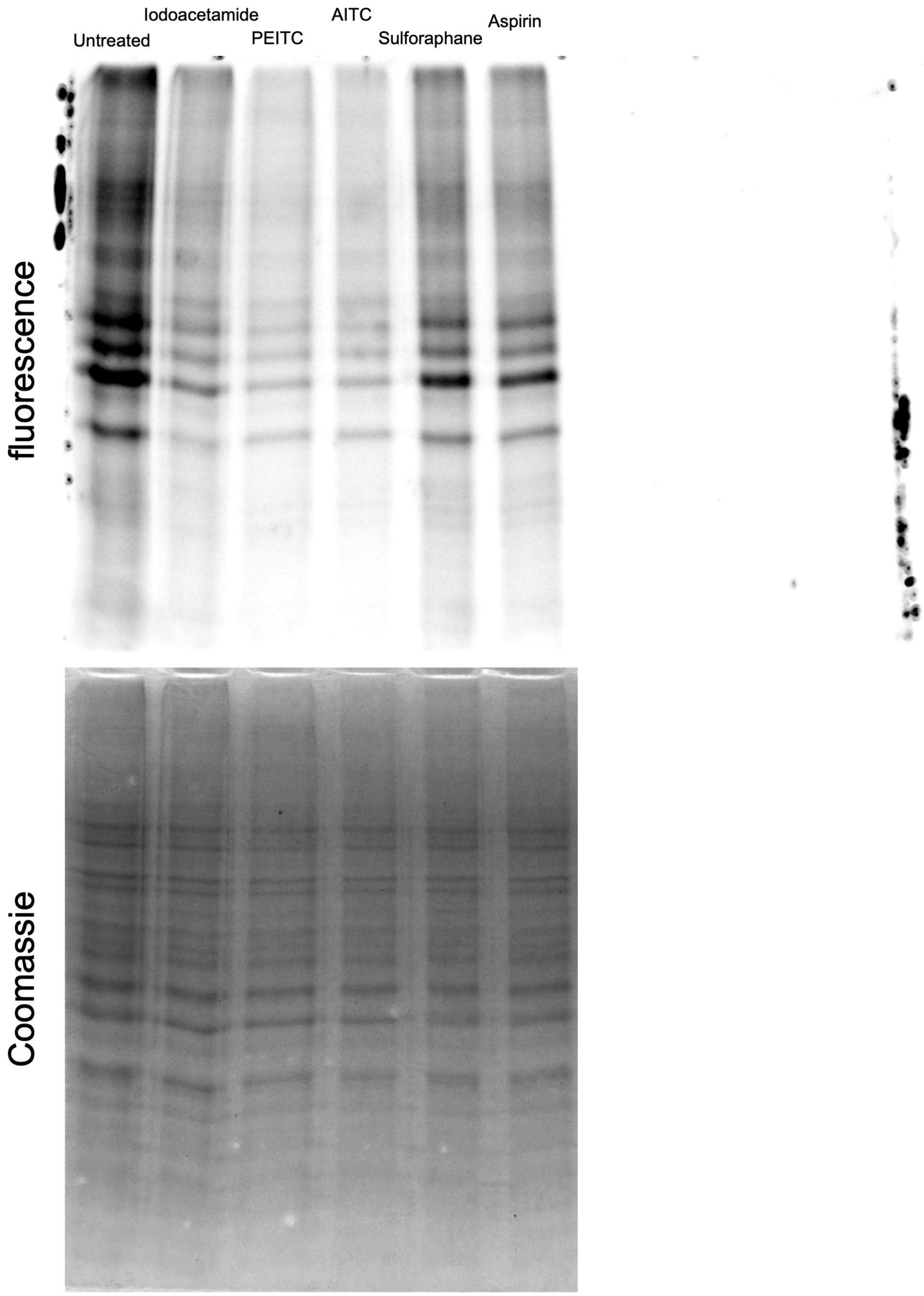


**Figure S13.** Uncropped in-gel fluorescence analysis of proteome-wide interactions of isothiocyanates and aspirin using cysteine-reactive probe. MDA-MB-231 cell lysates were treated with positive control iodoacetamide, isothiocyanates (100 µM, 30 min), or aspirin (1 mM, 30 min), then competed with IA-alkyne labeling (100 µM, 30 min), followed by a reaction with FAM-$N_3$ in the presence of $CuSO_4$, ascorbic acid, and THPTA. Cysteine reactivity was resolved by SDS-PAGE and visualized by in-gel fluorescence.

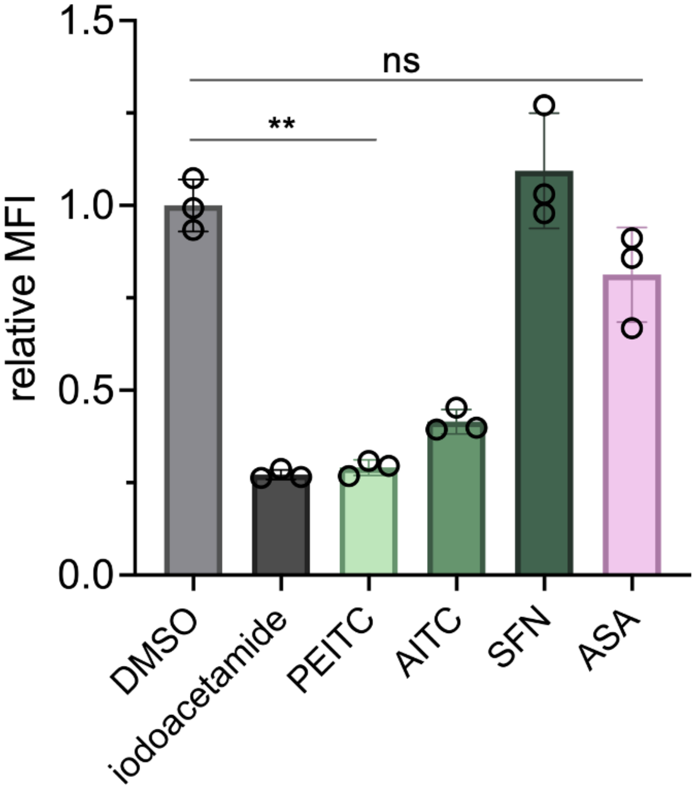


**Figure S14.** Flow cytometry analysis of proteome-wide interactions of isothiocyanates and aspirin using lysine-reactive probe. MDA-MB-231 cells were treated with positive control iodoacetamide, ITCs (200 µM, 1 h), or aspirin (1 mM, 1 h), then competed with IA-alkyne labeling (100 µM, 1 h), followed by a reaction with 3-azido-7-hydroxycoumarin in the presence of $CuSO_4$, ascorbic acid, and THPTA. Cysteine reactivity was assessed by flow cytometry, where a decrease in MFI (mean fluorescence intensity) relative to the DMSO control is representative of an increase in reactivity. Data are represented as mean ± SD ($n$ = 3). $P$-values were determined by a two-tailed $t$-test (ns = not significant, ** $p < 0.01$).

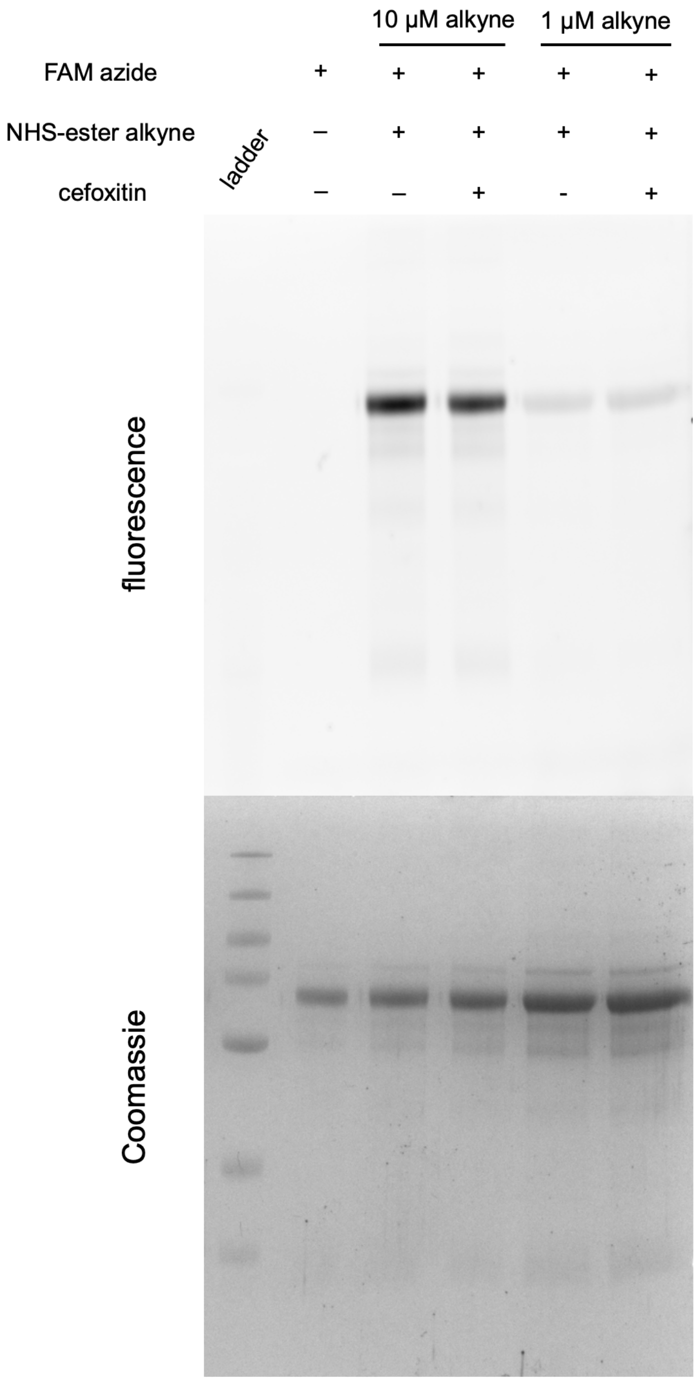


**Figure S15.** In-gel fluorescence analysis of cefoxitin reactivity using lysine-reactive probe in human serum. Human serum (0.5 mg/mL) was treated with cefoxitin (200 μM, 30 min), then competed with 4-pentynoic acid succinimidyl ester labeling (200 μM, 30 min), followed by a reaction with FAM-$N_3$ in the presence of $CuSO_4$, ascorbic acid, and THPTA. Lysine reactivity was resolved by SDS-PAGE and visualized by in-gel fluorescence.

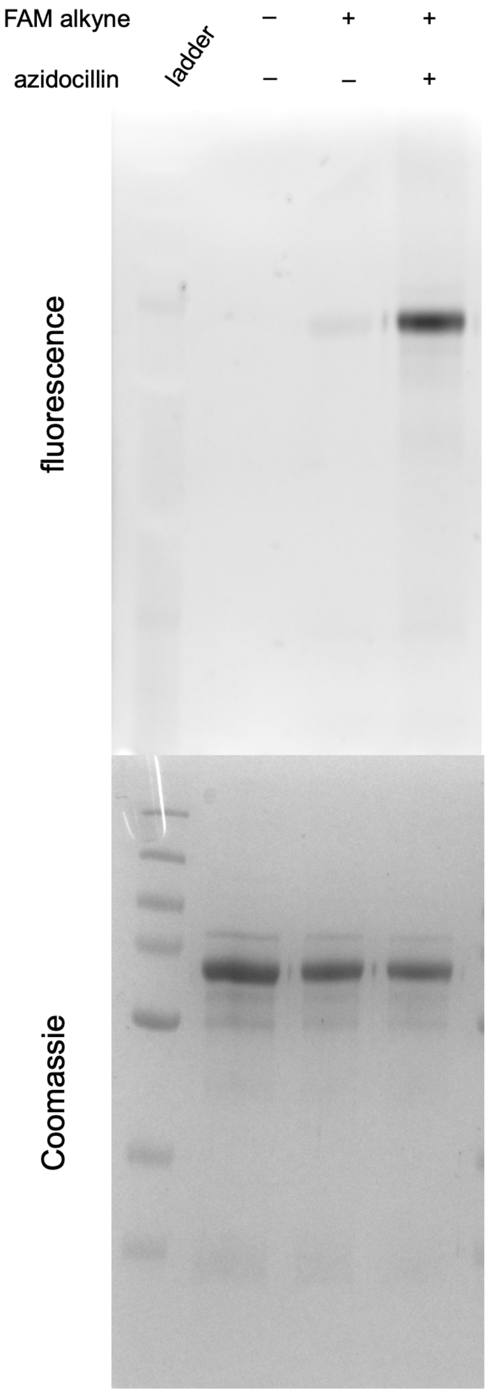


**Figure S16.** In-gel fluorescence analysis of azidocillin-modified HSA in human serum. Human serum (0.5 mg/mL) was treated with azidocillin (200 μM, 30 min), followed by a reaction with FAM-alkyne in the presence of $CuSO_4$, ascorbic acid, and THPTA. Reactivity of azidocillin was resolved by SDS-PAGE and visualized by in-gel fluorescence.

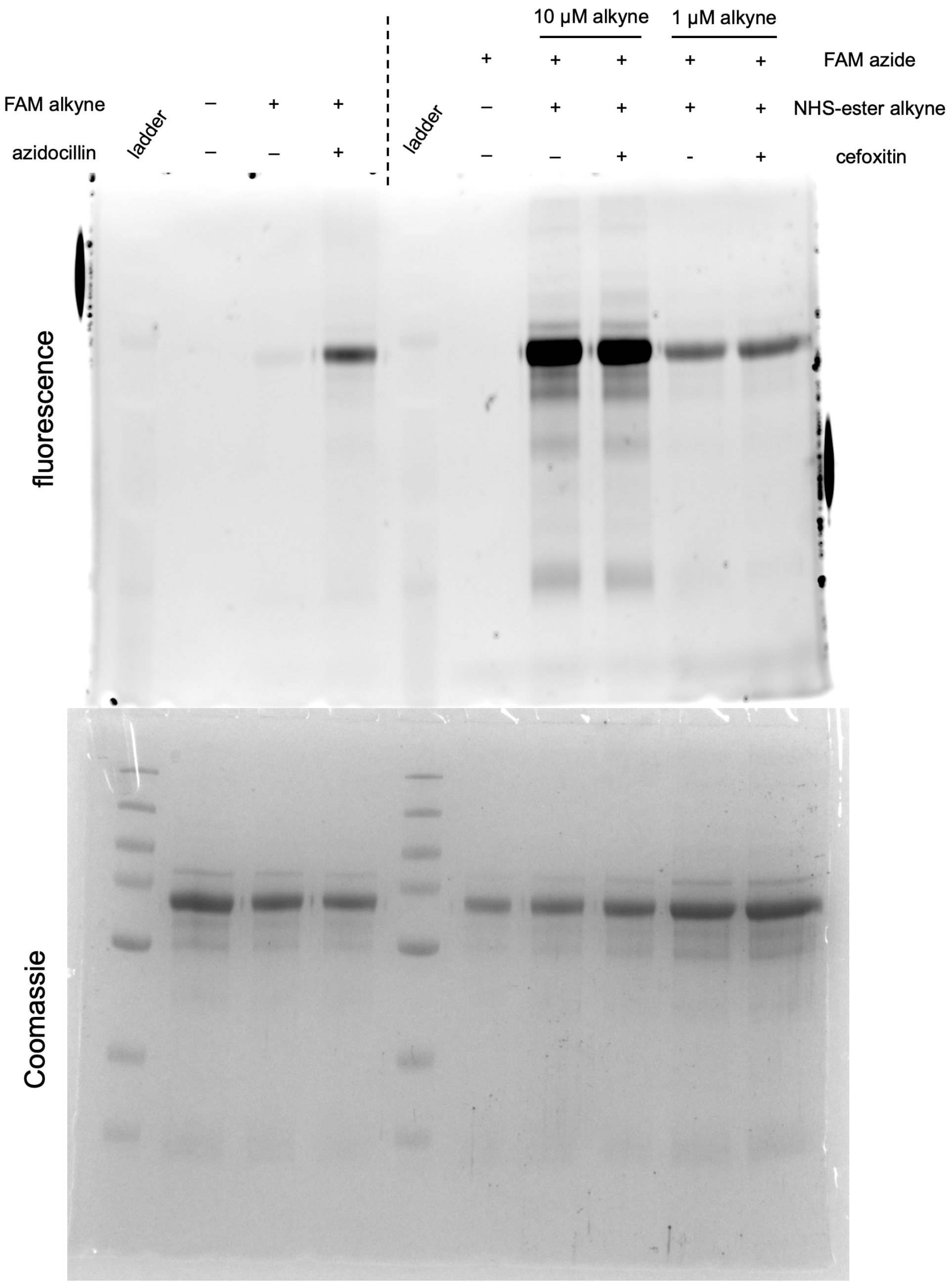


**Figure S17.** Uncropped in-gel fluorescence analyses of β-lactam interactions in human serum (corresponding to **Figs. S10, S11**). Lanes 2 – 4: Human serum (0.5 mg/mL) was treated with cefoxitin (200 μM, 30 min), then competed with 4-pentynoic acid succinimidyl ester labeling (200 μM, 30 min), followed by a reaction with FAM-$N_3$ in the presence of $CuSO_4$, ascorbic acid, and THPTA. Lanes 6 – 10: Human serum (0.5 mg/mL) was treated with azidocillin (200 μM, 30 min), followed by a reaction with FAM-alkyne in the presence of $CuSO_4$, ascorbic acid, and THPTA. Reactivity was resolved by SDS-PAGE and visualized by in-gel fluorescence.

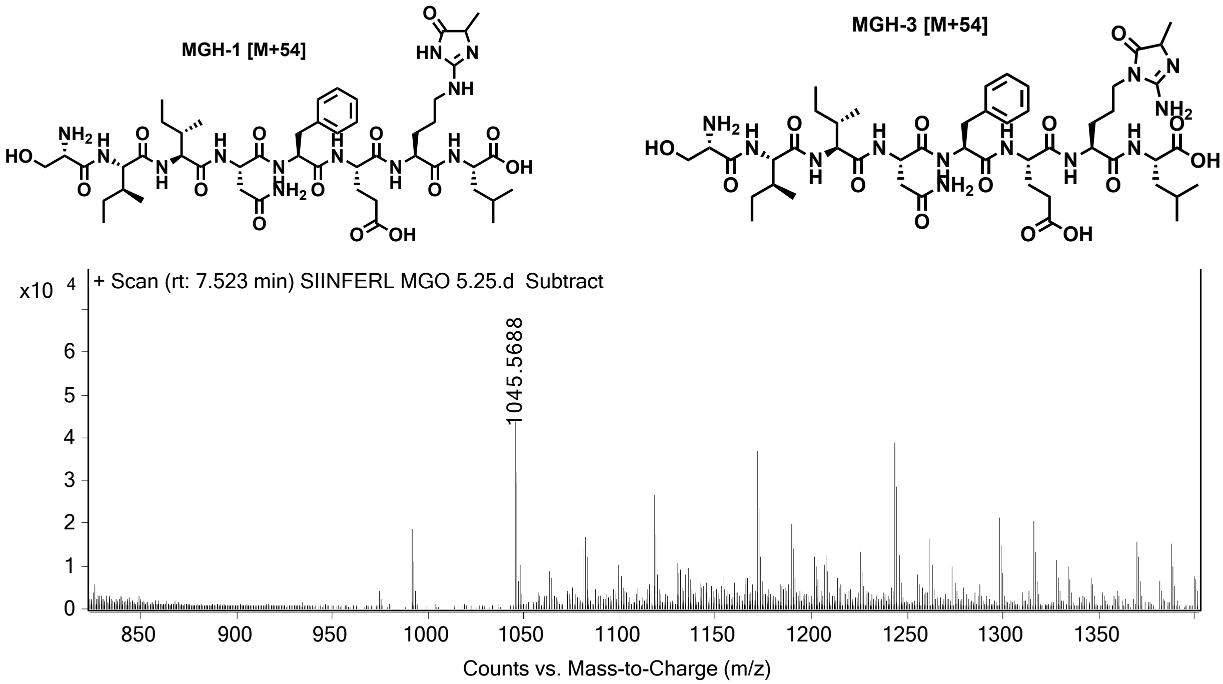


**Figure S18.** Extracted ion chromatogram (EIC) and corresponding chemical structures for (M+54) adduct of MGO modification reaction with **ovaK7R**. MGO-modified peptide identities were confirmed via high-resolution electrospray ionization mass spectrometry (HRMS, ESI/MS). Analyses were obtained on an Agilent 6545B Q-TOF LC/MS equipped with 1260 infinity II LC system with auto sampler. Samples were dissolved in $CH_3CN$ and eluted with a $CH_3CN/H_2O$ solution containing 0.1% formic acid.

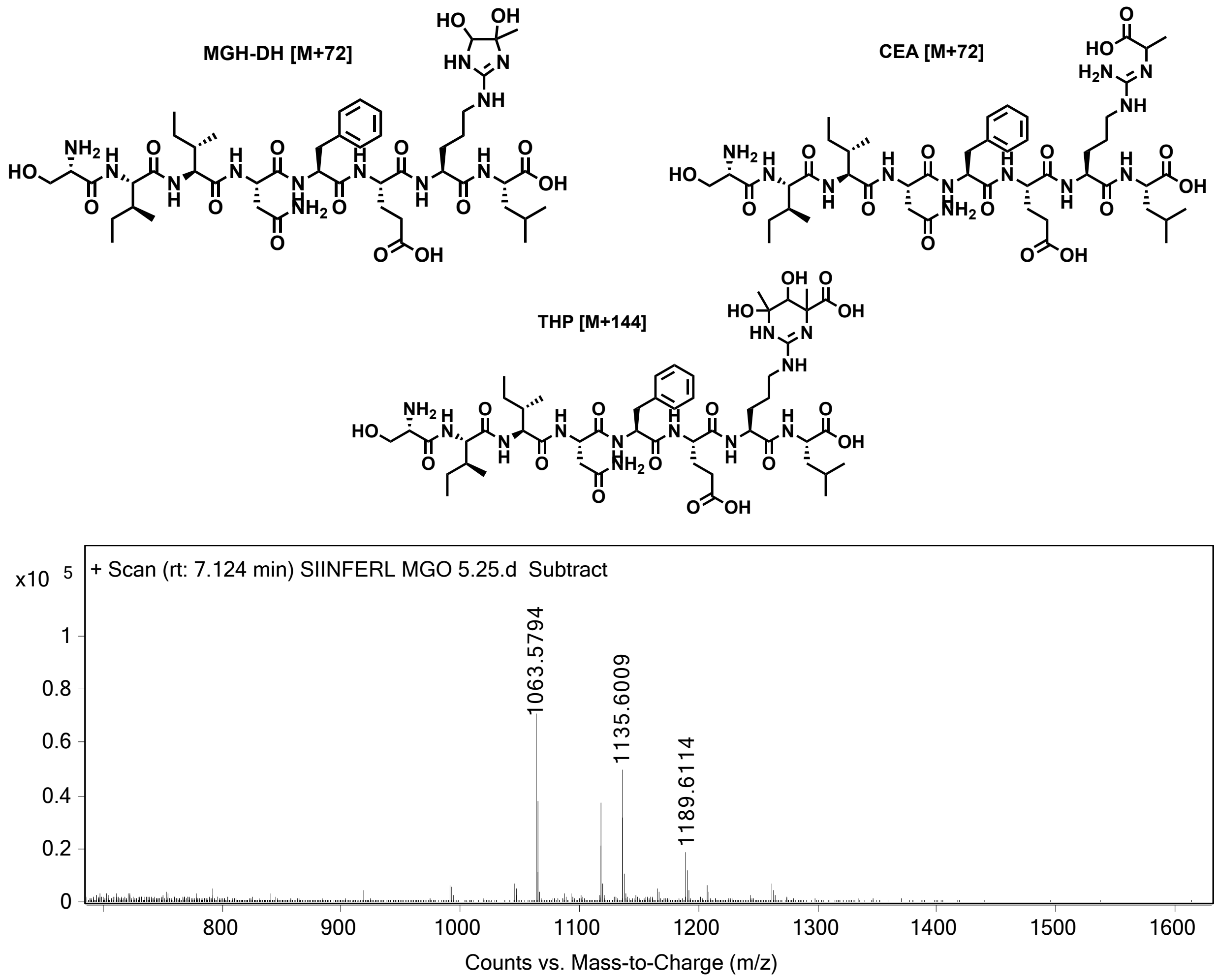


**Figure S19.** Extracted ion chromatogram (EIC) and corresponding chemical structures for (M+72) and (M+144) adducts of MGO modification reaction with **ovaK7R**. MGO-modified peptide identities were confirmed via high-resolution electrospray ionization mass spectrometry (HRMS, ESI/MS). Analyses were obtained on an Agilent 6545B Q-TOF LC/MS equipped with 1260 infinity II LC system with auto sampler. Samples were dissolved in $CH_3CN$ and eluted with a $CH_3CN/H_2O$ solution containing 0.1% formic acid.

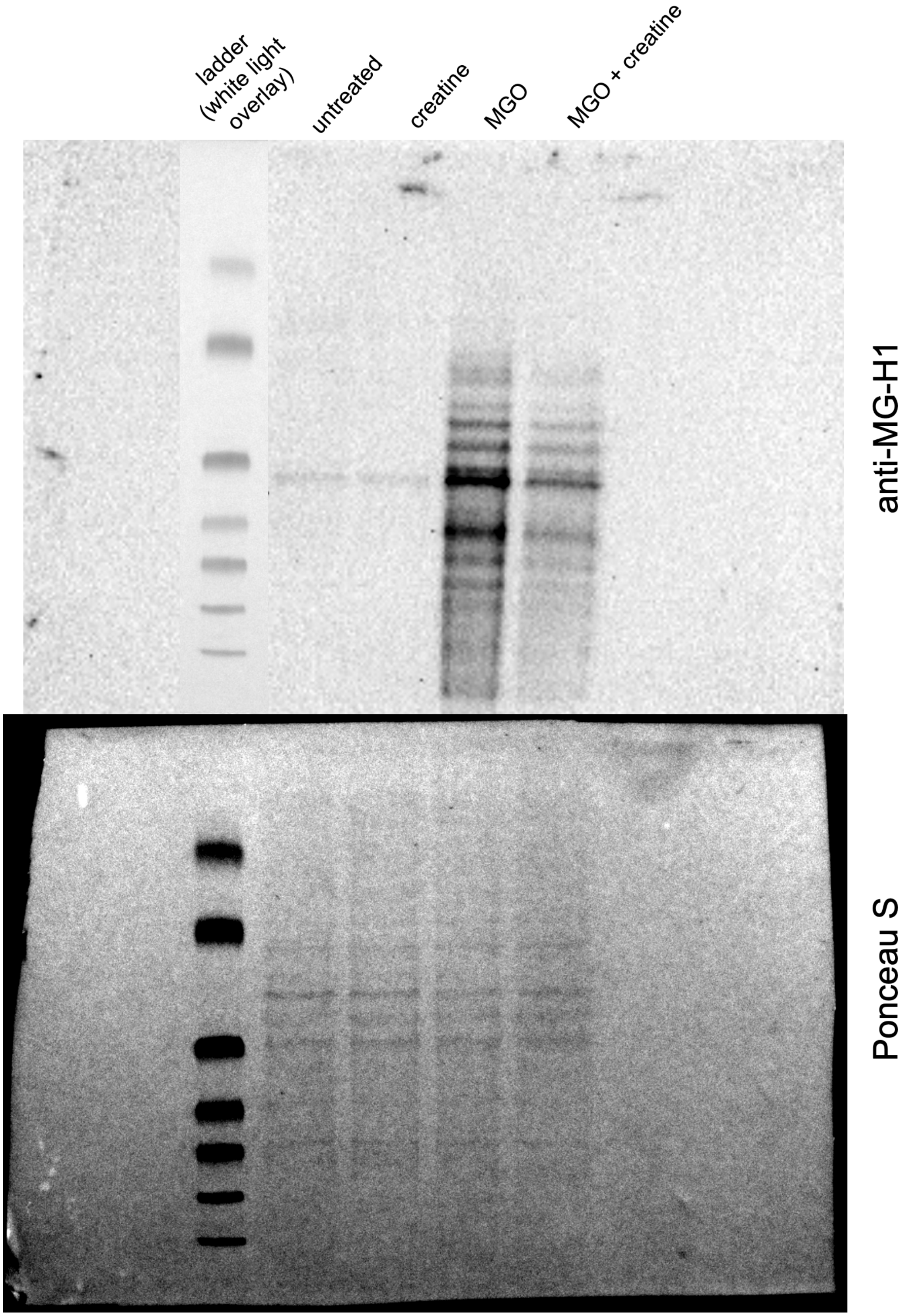


**Figure S20.** Uncropped Western blot analysis showing MGO-dependent labeling and creatine-mediated “scavenging” of MGO. MDA-MB-231 cells were treated for 1 h with MGO (800 μM), creatine (10 mM), or co-treatment with both, and probed with anti-MG-H1 antibody (clone 1H7G5).

#### MATERIALS AND METHODS

##### Materials

| REAGENTS | VENDOR SOURCE | CATALOG # |
|---|---|---|
| **Reagents for biological methods** | | |
| RPMI 1640 Medium | Sigma-Aldrich | R8758 |
| DMEM, high glucose, pyruvate | Sigma-Aldrich | 11995065 |
| Fetal Bovine Serum | Sigma-Aldrich | A52568-01 |
| Penicillin-Streptomycin | Sigma-Aldrich | P4333 |
| G418 disulfate, 50 mg/mL solution | Fisher Scientific | AAJ63871AB |
| TrypLE™ Express Enzyme (1X), phenol red | Fisher Scientific | 12605028 |
| Human Serum | Sigma-Aldrich | H4522 |
| APC anti-mouse H-2K$^b$ antibody | BioLegend | 116518 |
| Recombinant Murine IFN-γ | PeproTech | 315-05 |
| Ultra-LEAF Purified anti-human HLA-A,B,C antibody | BioLegend | 311441 |
| MG-H1 Monoclonal Antibody (1H7G5) | Invitrogen | MA5-49069 |
| HRP Goat anti-mouse IgG (minimal x-reactivity) Antibody | BioLegend | 405306 |
| Saponin | Thermo Scientific | A18820.14 |
| Chlorophenol Red β-D-Galactopyranoside (CPRG) | Cayman Chemical | 29707 |
| 2-Mercaptoethanol | Sigma-Aldrich | M3148 |
| Magnesium chloride | Sigma-Aldrich | 208337 |
| Phosphate buffered saline | Sigma-Aldrich | P3813 |
| 40% Acrylamide/Bis Solution, 19:1 | Bio-Rad | 1610144 |
| Ammonium Persulfate (APS) | Bio-Rad | 1610700 |
| TEMED | Bio-Rad | 1610800 |
| Sodium Dodecyl Sulfate (SDS) | Thermo Scientific | J18220 |
| N-[2-(pent-4-ynoylsulfanyl)ethyl]acetamide (**1**) | Enamine | EN300-37325821 |
| Acetic acid, glacial, ≥99% | Sigma-Aldrich | A6283 |
| DADPS Biotin Azide | Vector Laboratories | CCT-1330-1 |
| CHAPS | Thermo Scientific | 28300 |
| Ethylenediaminetetraacetic acid (EDTA) | Sigma-Aldrich | E9884 |

| | | |
|---|---|---|
| Triton X-100 | Sigma-Aldrich | 93443 |
| Tween-20 | Sigma-Aldrich | P1379 |
| NHS Act Sepharose 4 Fast Flow | Sigma-Aldrich | GE17-0906-01 |
| FAM azide, 6-isomer | Lumiprobe | A5130 |
| FAM alkyne, 6-isomer | BroadPharm | BP-22531 |
| 3-azido-7-hydroxy Coumarin | Cayman Chemical | 10596 |
| Cupric sulfate pentahydrate | Fisher Scientific | C493 |
| THPTA | BroadPharm | BP-26340 |
| L-Ascorbic acid | Cayman Chemical | 14656 |
| IA-Alkyne | AmBeed | A150286 |
| 4-pentynoic acid succinimidyl ester | BroadPharm | BP-28420 |
| 2-Iodoacetamide | AmBeed | A573639 |
| Glyphosate | MedChemExpress | HY-B0863 |
| Metolachlor | MedChemExpress | HY-B1871 |
| Cyhalothrin | MedChemExpress | HY-B0836 |
| Chlorpyrifos | Cayman Chemical | 21412 |
| Chlorothalonil | MedChemExpress | HY-N6625 |
| Dieldrin | Cayman Chemical | 24042 |
| Epoxiconazole | MedChemExpress | HY-119683 |
| Atrazine | MedChemExpress | HY-N7091 |
| 4-Amino-3,5,6-trichloropyridine-2-carboxylic acid, 95% (Picloram) | AK Scientific | J51358 |
| Paraquat (chloride) | Cayman Chemical | 38521 |
| Chloremquat (chloride) | AK Scientific | J51995 |
| Allyl isothiocyanate | Thermo Scientific | L02901.06 |
| 2-Phenylethyl isothiocyanate | AK Scientific | J53261 |
| DL-Sulforaphane, 98% | AK Scientific | Y0196 |
| 2-Acetoxybezoic acid, 98% | AK Scientific | J52104 |
| Cefoxitin | AK Scientific | E827 |
| Azidocillin | MedChemExpress | HY-B2091 |
| Pyruvic aldehyde, 35-45%, w/w aq. soln (MGO) | AK Scientific | H451 |
| Creatine anhydrous | AK Scientific | J51321 |
| Piece BCA Protein Assay Kit | Thermo Scientific | A55865 |

| | | |
|---|---|---|
| Western Blotting Luminol Reagent | Santa Cruz Biotechnology | sc-2048 |
| Dry Milk Powder | RPI Corp | M17200-500.0 |
| Ponceau S Staining Solution | Thermo Scientific | A40000279 |
| **Reagents for synthesis and characterization** | | |
| Fmoc-L-leucine 4-alkoxybenzyl alcohol resin (100-200 mesh, 0.3-0.8 meq/g) | ChemImpex | 01911 |
| 2-Chlorotrityl chloride resin (1.0-2.0 meq/g, 100-200 mesh) | ChemImpex | 03498 |
| α-*N*-Fmoc-amino acids | Chem Impex | Various |
| N, N'-Diisopropylcarbodiimide (DIC) | Chem Impex | 00100 |
| Ethyl Cyano(hydroxyimino)acetate (Oxyma) | TCI Chemicals | E0847 |
| Tris-(carboxyethyl)phosphine hydrochloride (TCEP) | Chem Impex | 23004 |
| DL-Dithiothreitol | Sigma-Aldrich | D0632 |
| 2-Bromoethylamine HBr | AK Scientific | J91302 |
| N, N-Dimethylformamide (DMF) | Sigma-Aldrich | 319937 |
| Dichloromethane (DCM) ACS-grade | Sigma-Aldrich | D65100 |
| Methanol ACS-grade | Sigma-Aldrich | 179337 |
| Acetonitrile HPLC-grade | Sigma-Aldrich | 34851 |
| Piperidine | Sigma-Aldrich | 104094 |
| Trifluoroacetic acid (TFA) reagent-grade | ChemImpex | 00289 |
| Trifluoroacetic acid (TFA) HPLC-grade | Sigma-Aldrich | 302031 |
| Triisopropylsilane | ChemImpex | 01966 |
| **General materials and equipment** | | |
| Greiner Bio-One CELLSTAR TC Treated Cell Culture Flasks | Fisher Scientific | 07-000-225 |
| Corning™ Costar™ 96-Well, Cell Culture-Treated, Flat-Bottom Microplate | Fisher Scientific | 09-761-145 |
| Greiner Bio-One 96-well Non-treated Polystyrene Microplates | Fisher Scientific | 07-000-124 |
| Oasis HLB 1cc (30mg) Extraction Cartridges | Waters | WAT094225 |
| Amicon Ultra Centrifugal Filter, 3 kDa MWCO | Sigma-Aldrich | UFC9003 |
| Nitrocellulose Membranes, 0.45 µm | Thermo Scientific | 88018 |

**ovaWT** (SIINFEKL)

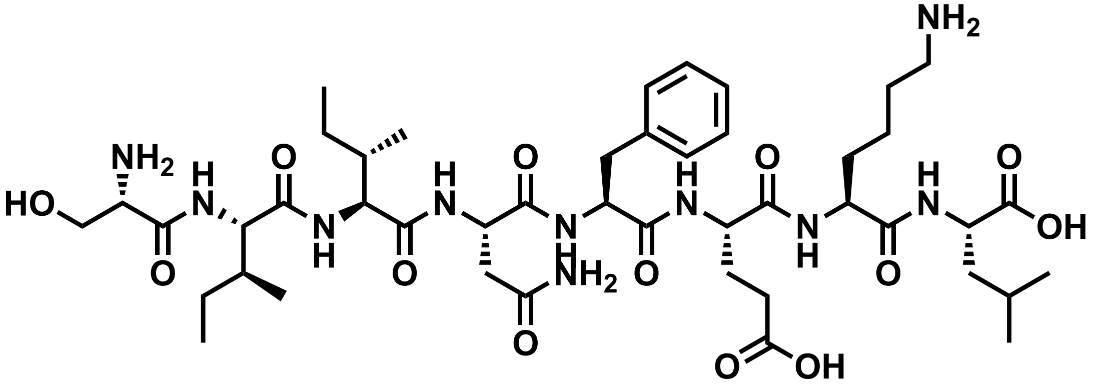

Chemical structure of **ovaWT**.

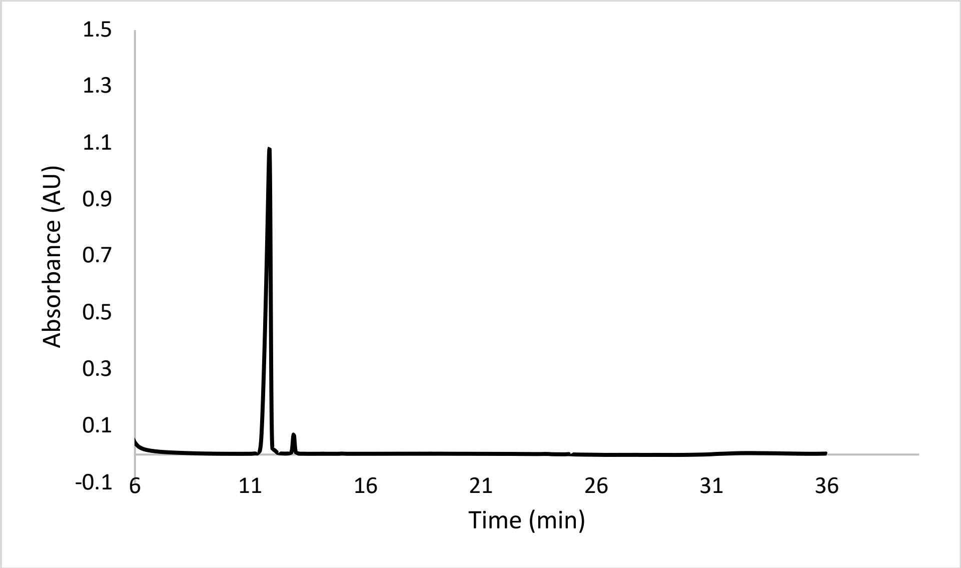


Analytical HPLC Chromatogram of **ovaWT**

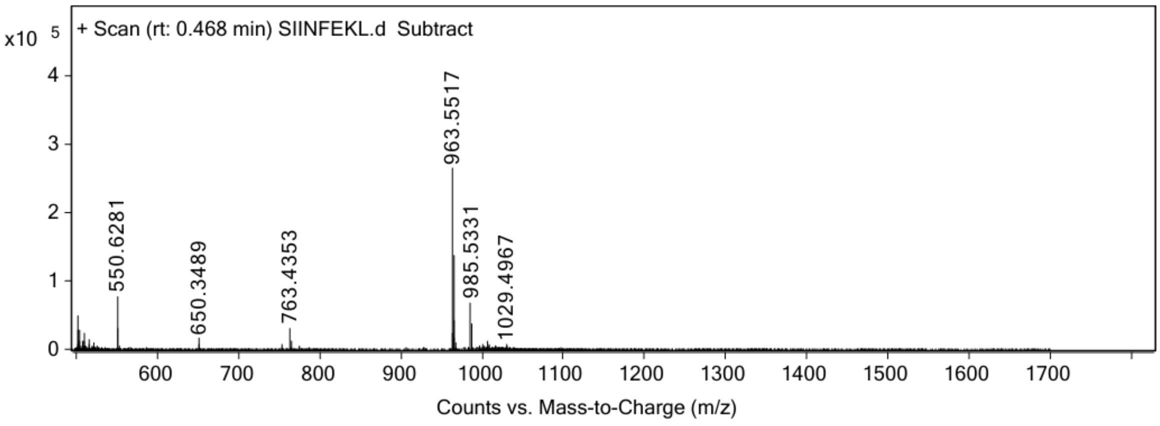


ESI-MS for **ovaWT** (calculated [M+H$^+$]: 963.5515, found 963.5517)

**cntPEP** (SNFVSAGI)

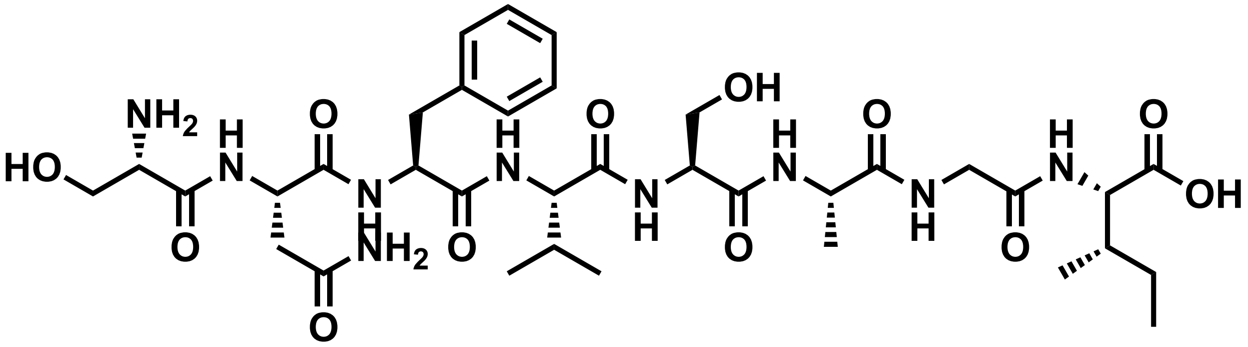


Chemical structure of **cntPEP**.

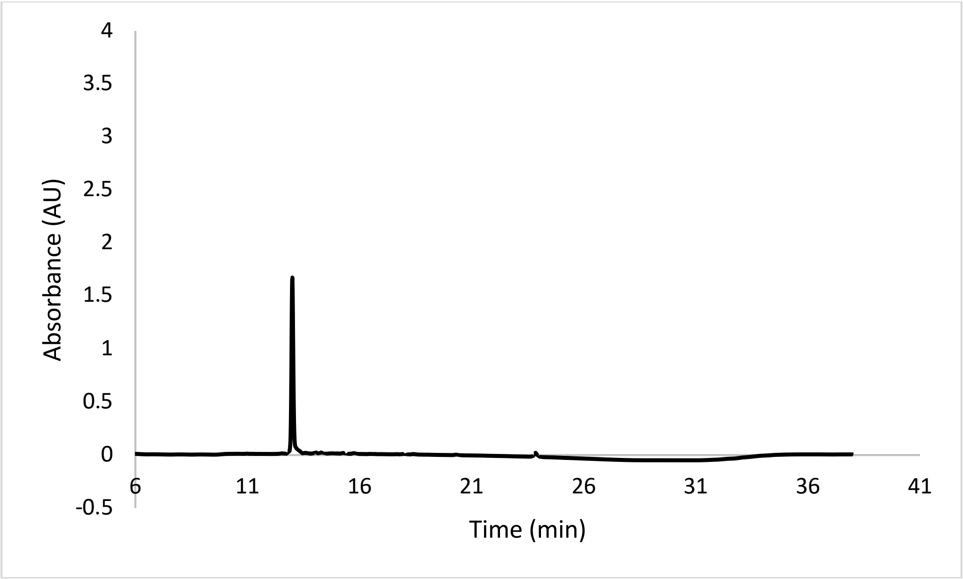


Analytical HPLC Chromatogram of **cntPEP**.

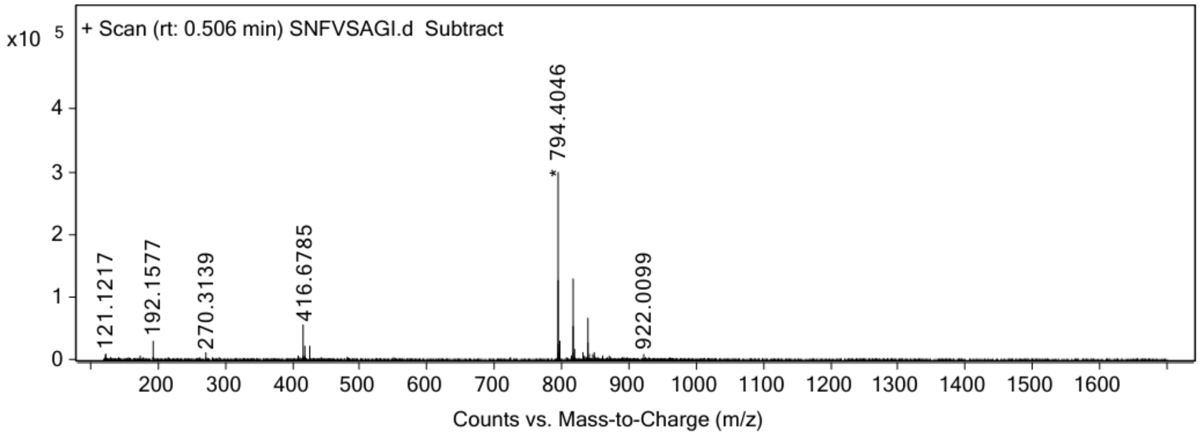


ESI-MS for **cntPEP** (calculated [$M+H^+$]: 794.4043, found 794.4046).

**ovaK7$_{\mathbf{formyl}}$**

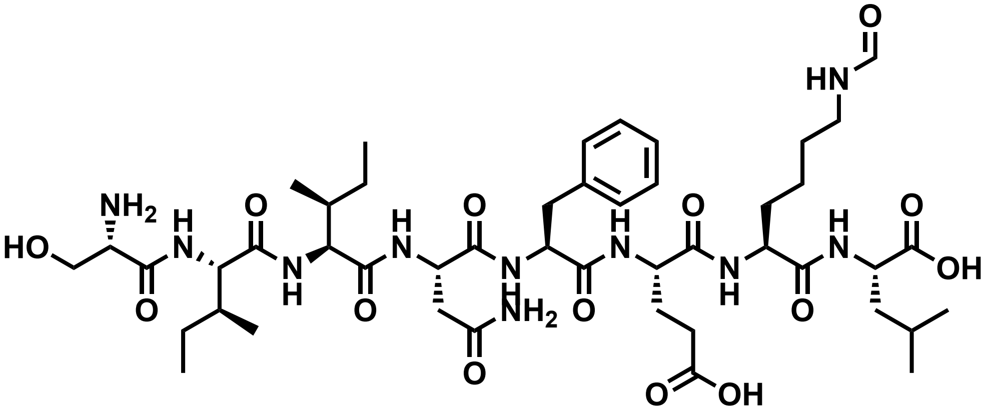

Chemical structure of **ovaK7$_{\mathbf{formyl}}$.**

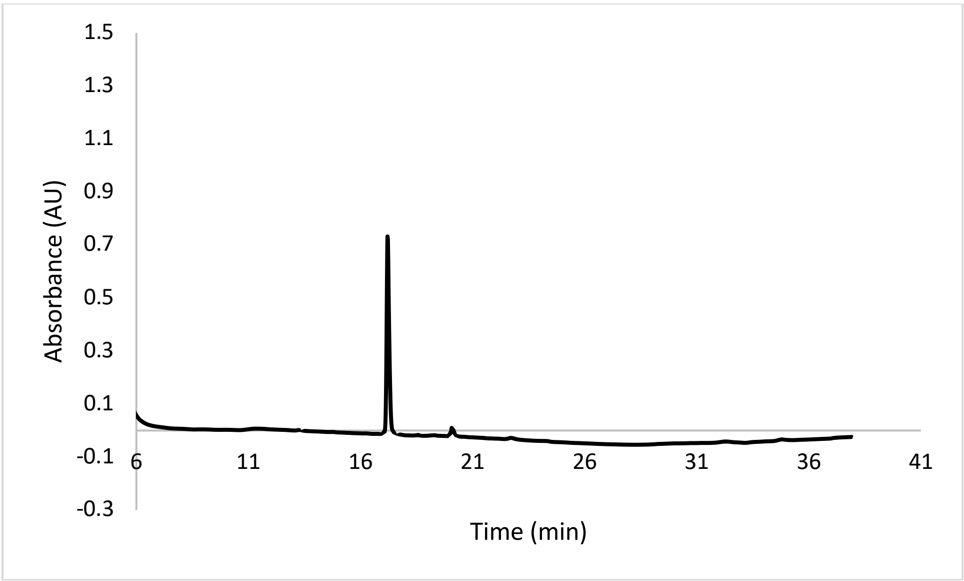


Analytical HPLC Chromatogram of **ovaK7$_{\mathbf{formyl}}$**.

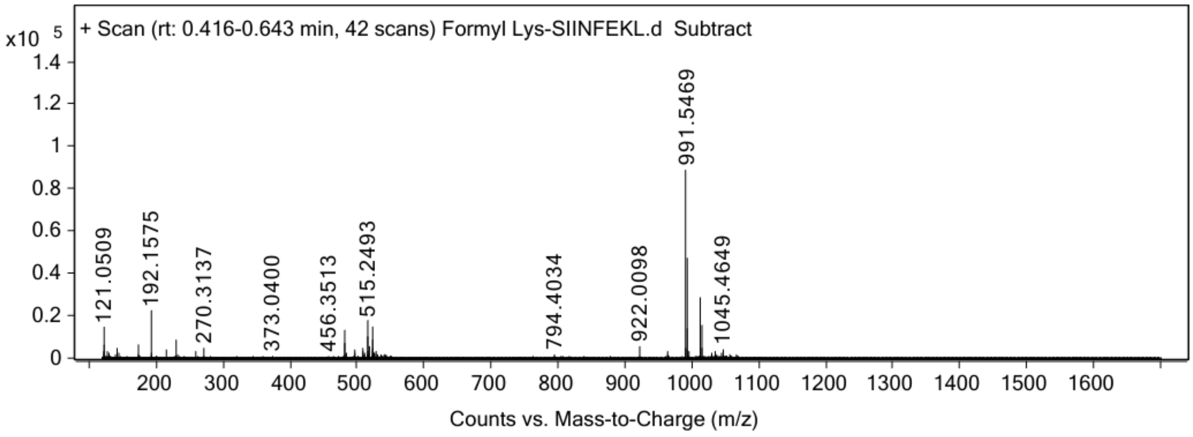


ESI-MS for **ovaK7$_{\mathbf{formyl}}$** (calculated [M+H$^+$]: 991.5459, found 991.5469).

**ovaK7$_{aad}$**

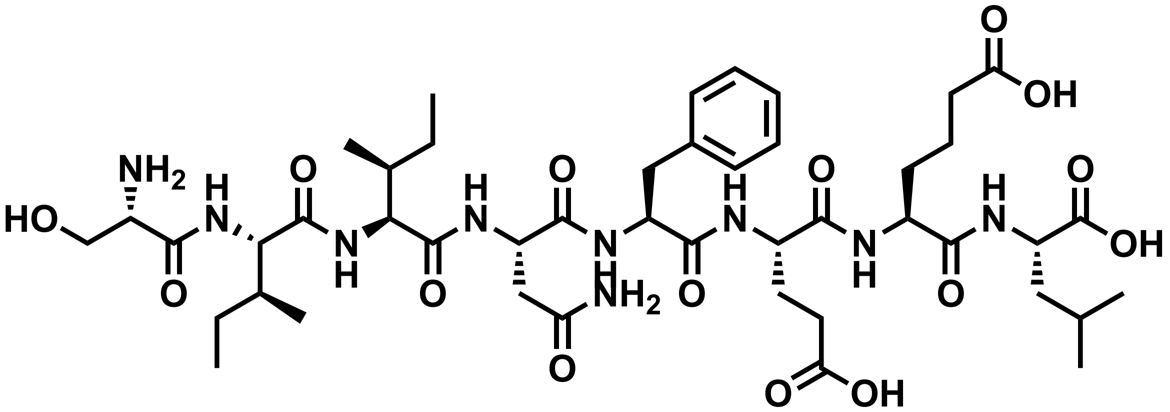


Chemical structure of **ovaK7$_{aad}$.**

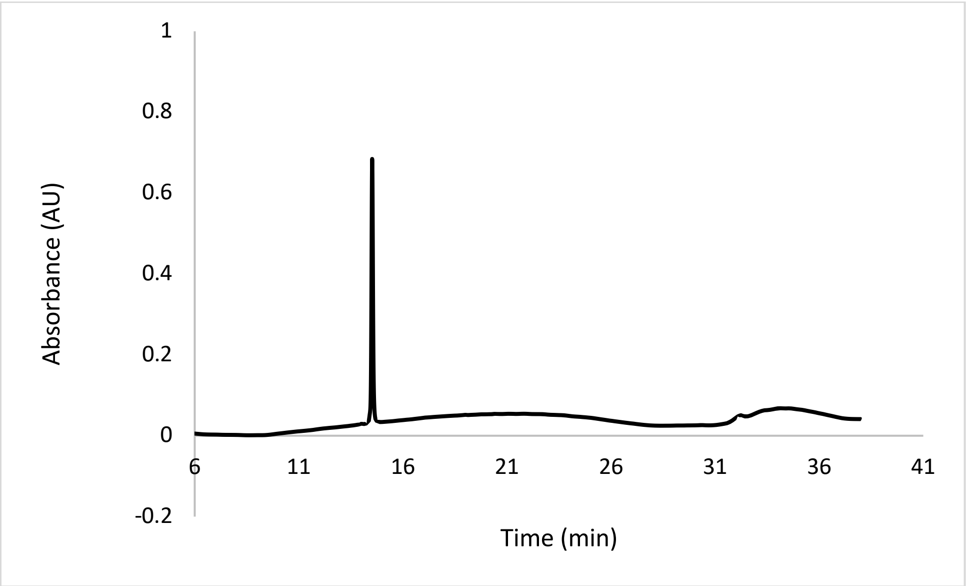


Analytical HPLC Chromatogram of **ovaK7$_{aad}$**

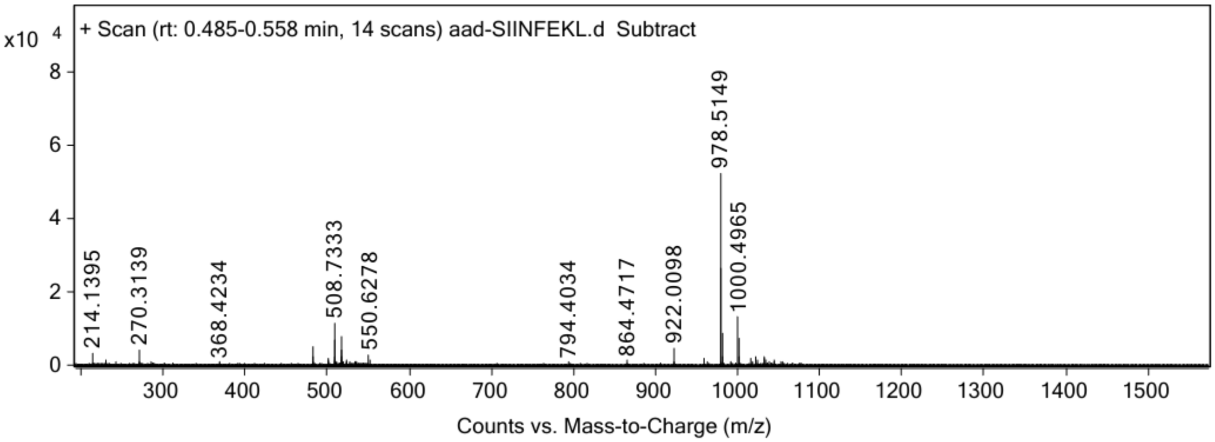


ESI-MS for **ovaK7$_{aad}$** (calculated [$M+H^+$]: 978.5142, found 978.5149).

**ovaK7$_{hcit}$**

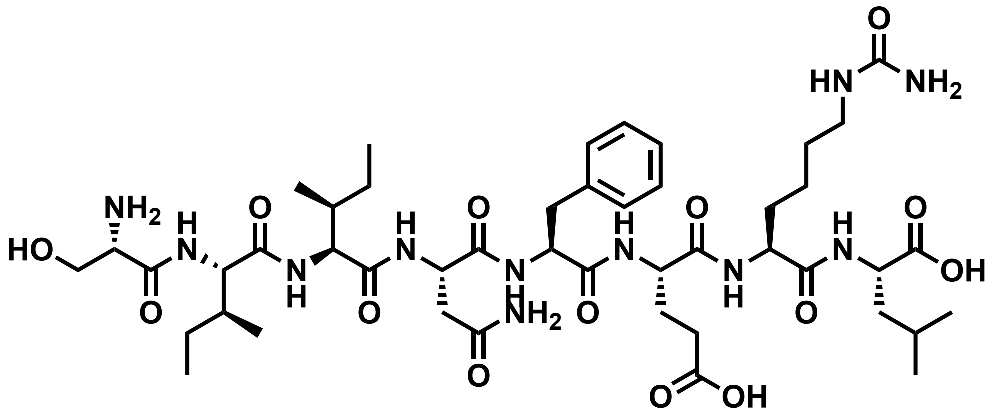


Chemical structure of **ovaK7$_{hcit}$.**

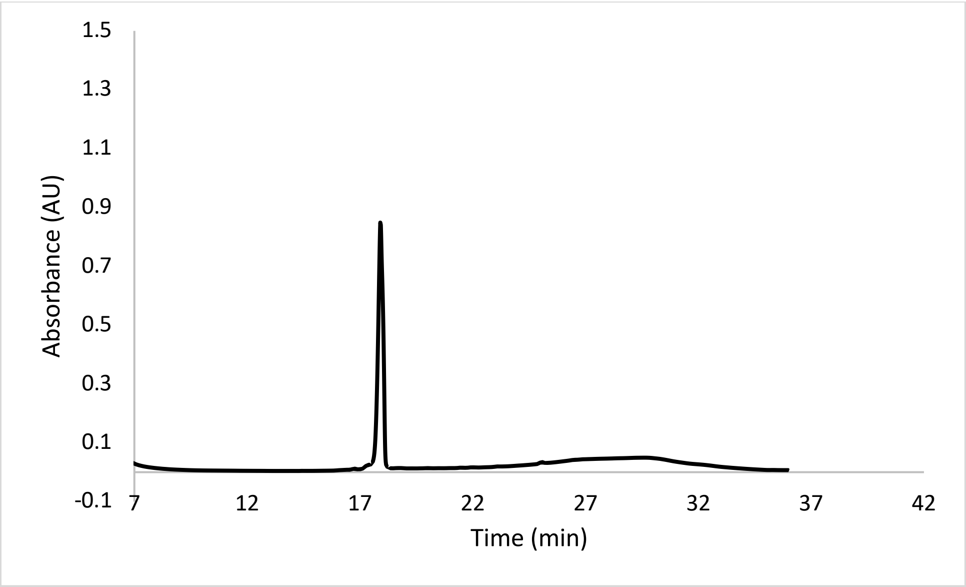


Analytical HPLC Chromatogram of **ovaK7$_{hcit}$.**

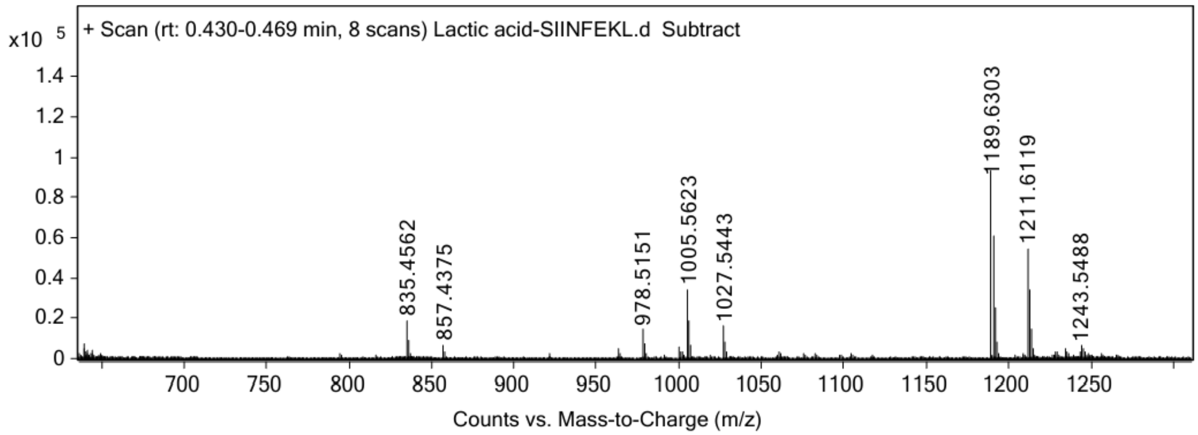


ESI-MS for of **ovaK7$_{hcit}$** calculated [M+H$^+$]: 1005.5494, found 1005.5623

**ovaK7$_{lac}$**

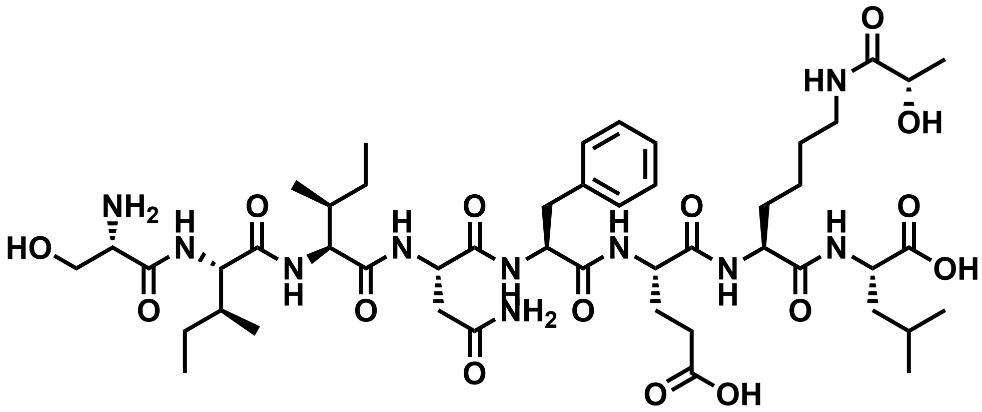

Chemical structure of **ovaK7$_{lac}$**.

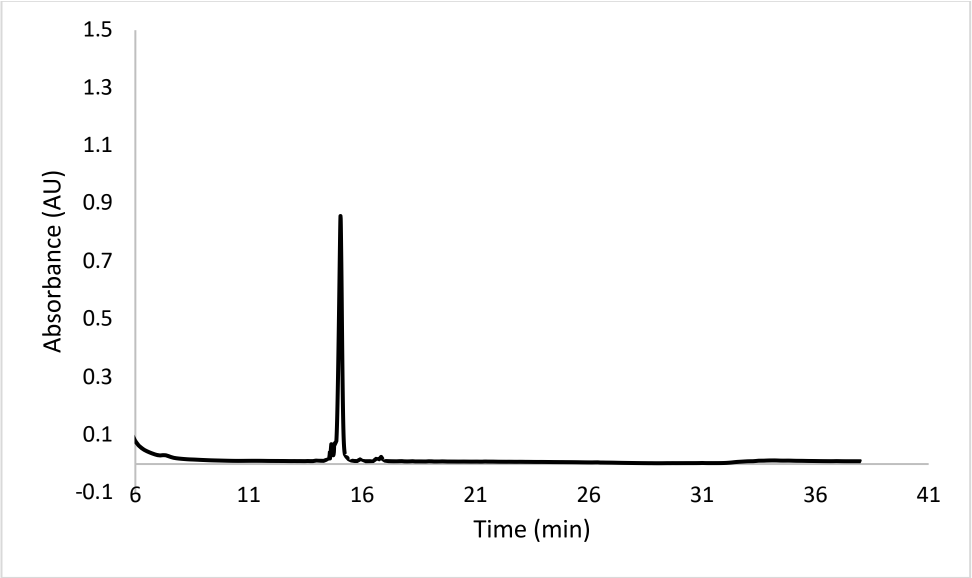


Analytical HPLC Chromatogram of **ovaK7$_{lac}$**.

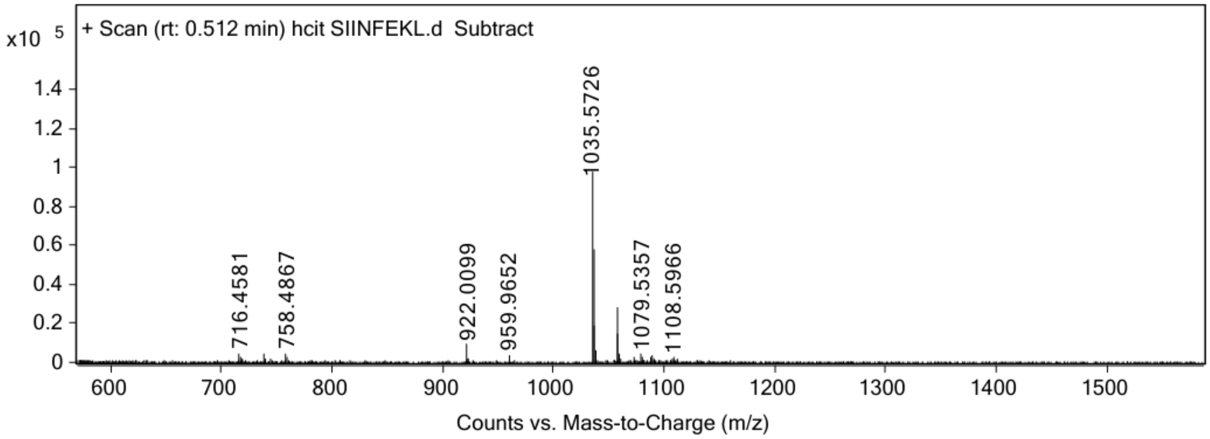


ESI-MS for **ovaK7$_{lac}$** (calculated [M+H$^+$]: 1035.5721, found 1035.5726).

**ovaK7$_{suc}$**

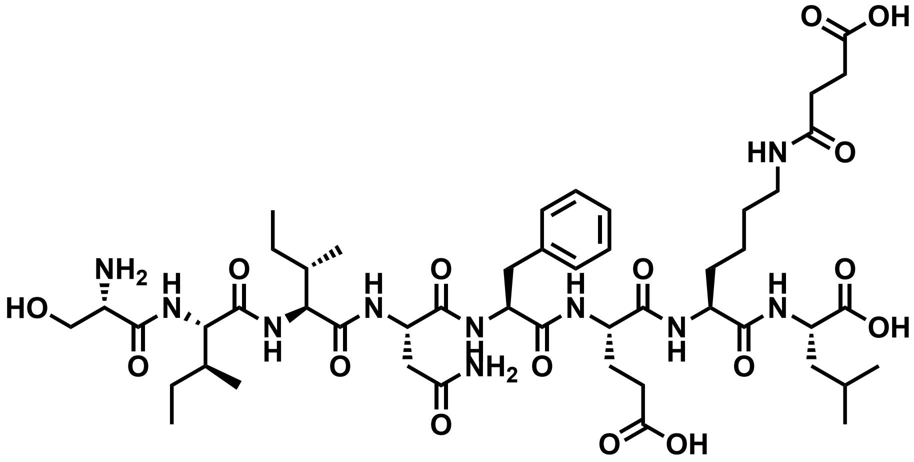

Chemical structure of **ovaK7$_{suc}$.**

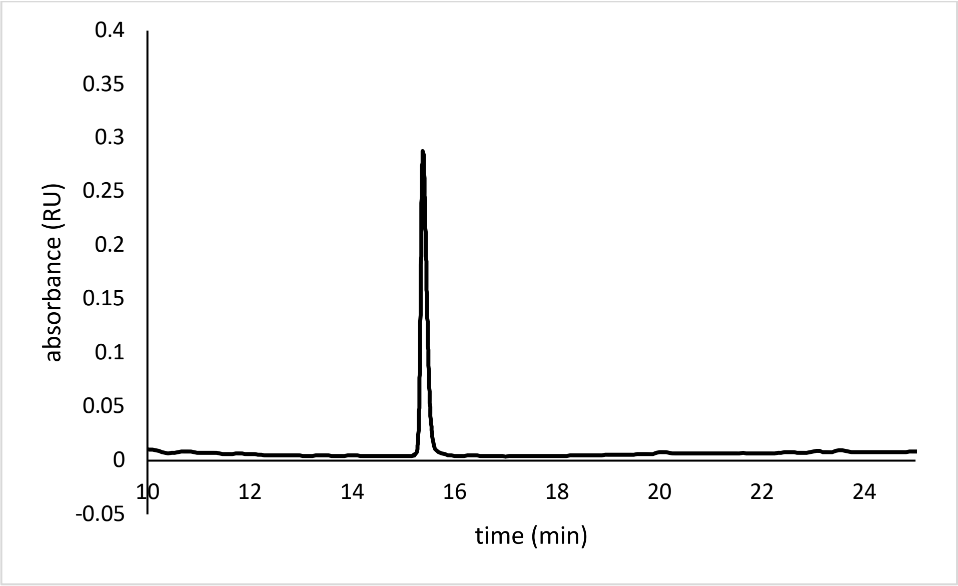


Analytical HPLC Chromatogram of **ovaK7$_{lac}$.**

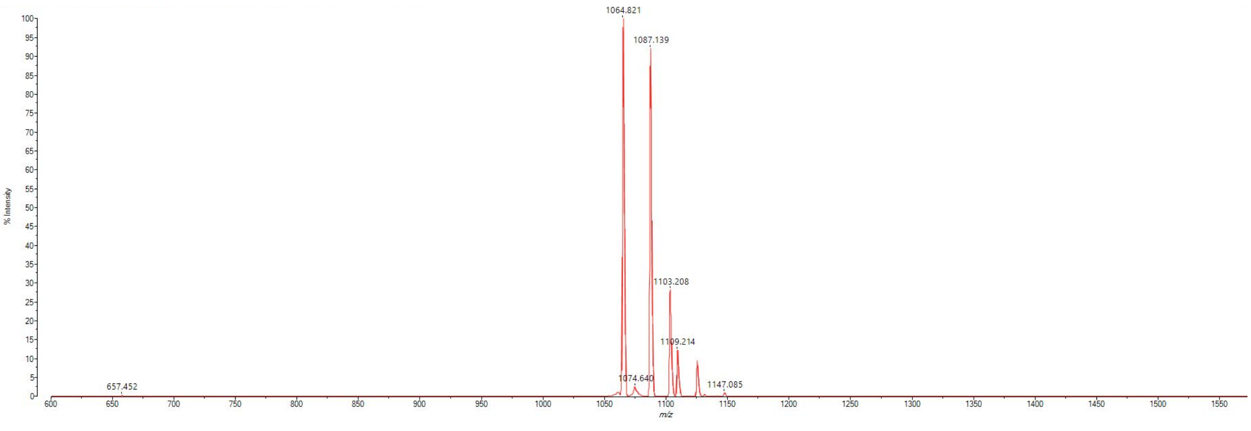


MALDI-TOF mass spectrum for **ovaK7$_{lac}$** (m/z 1063.567 found 1064.821).

**ovaE6C** (SIINFCKL)

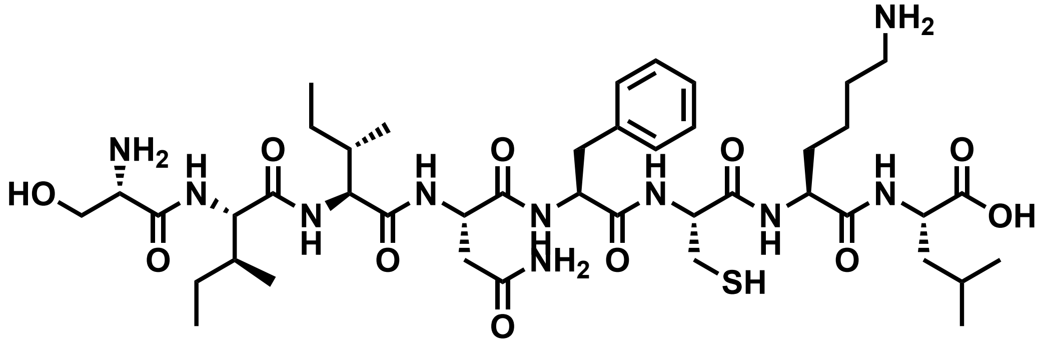


Chemical structure of **ovaE6C.**

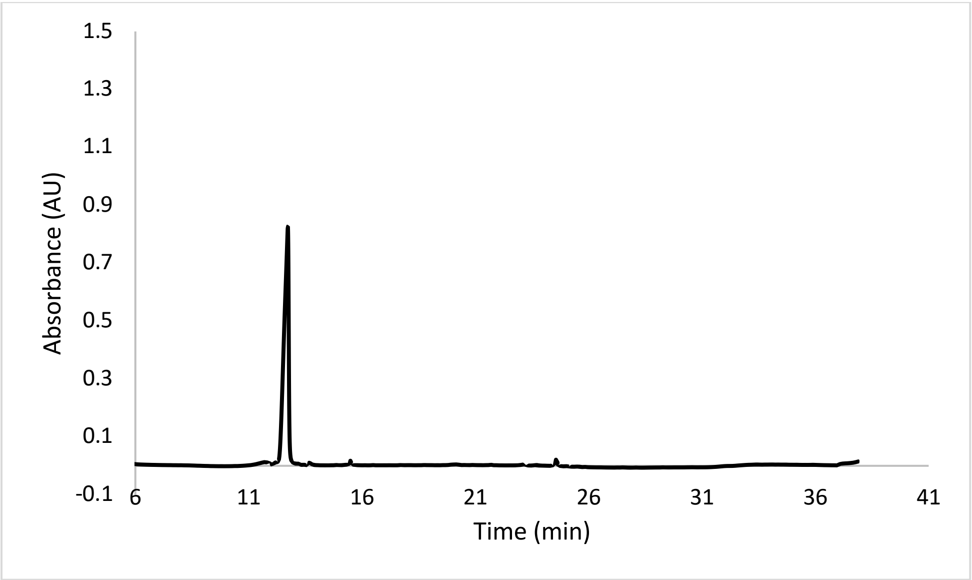


Analytical HPLC Chromatogram of **ovaE6C**.

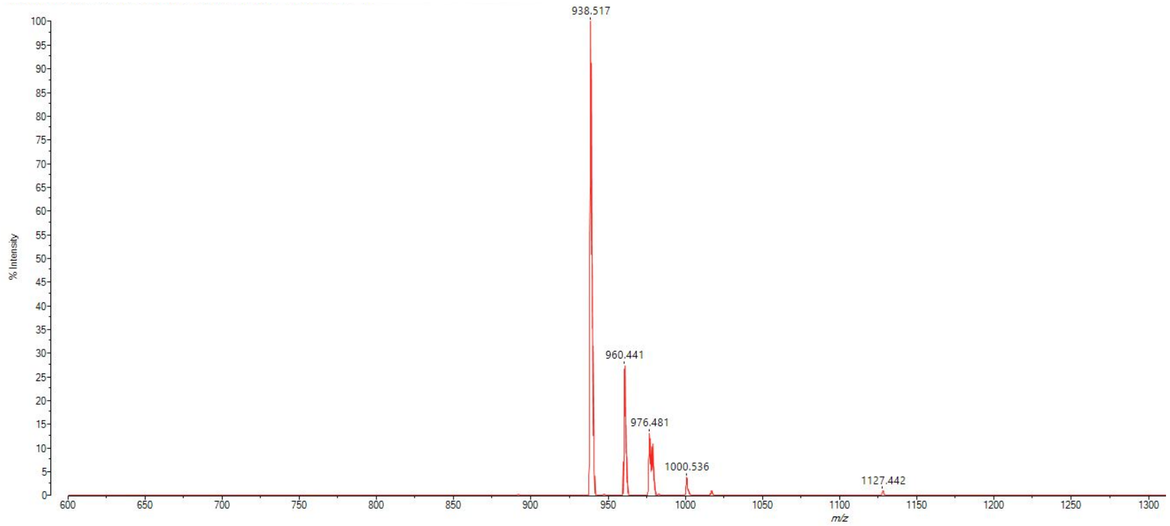


MALDI-TOF mass spectrum for **ovaE6C** (m/z 937.518 found 938.517).

**ovaE6C$_{SO3}$**

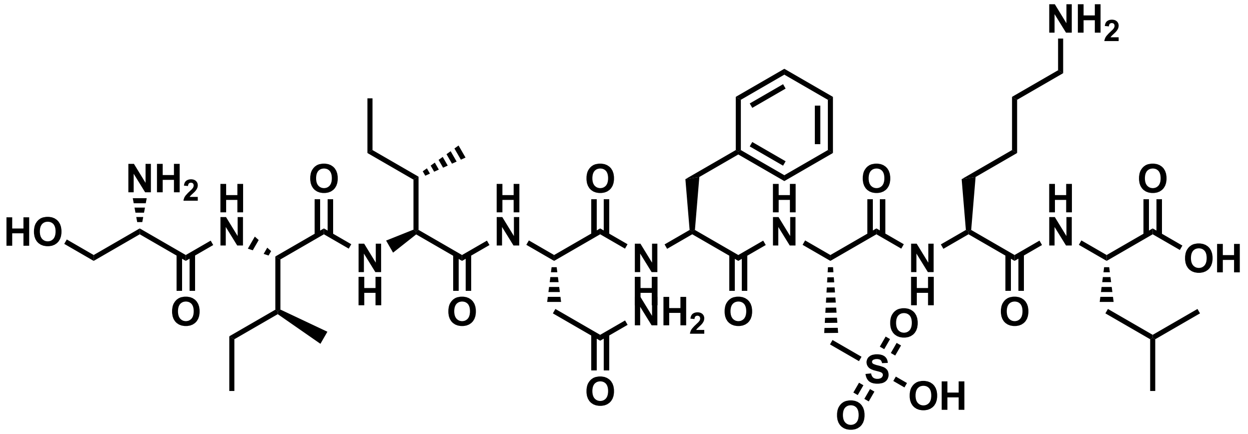


Chemical structure of **ovaE6C$_{SO3}$**.

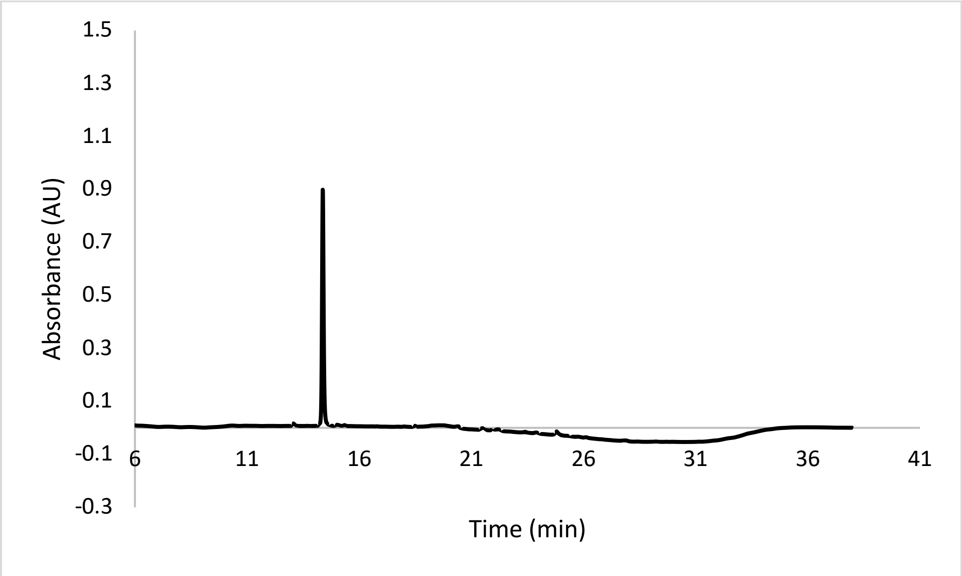


Analytical HPLC Chromatogram of **ovaE6C$_{SO3}$**.

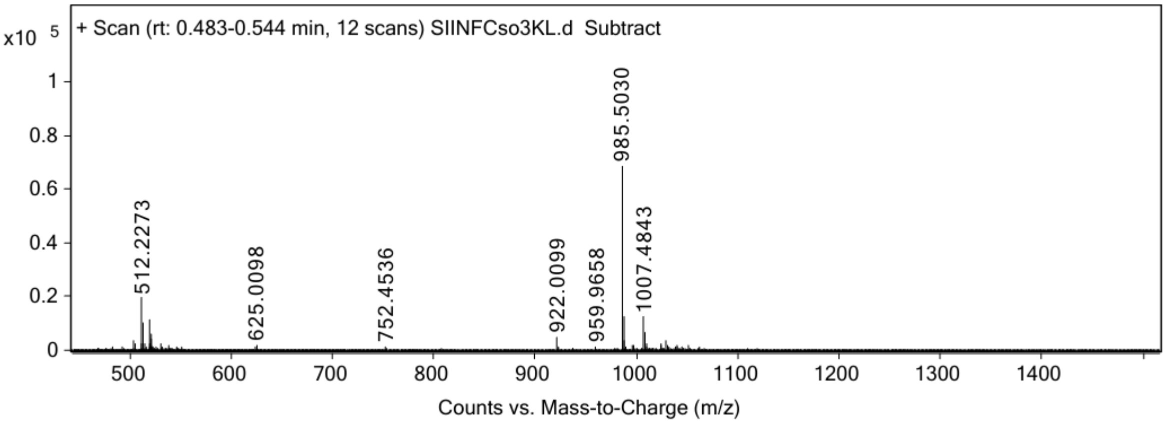


ESI-MS for **ovaE6C$_{SO3}$** (calculated [M+H$^+$]: 985.5023, found 985.5030).

**ovaE6C$_{2sc}$**

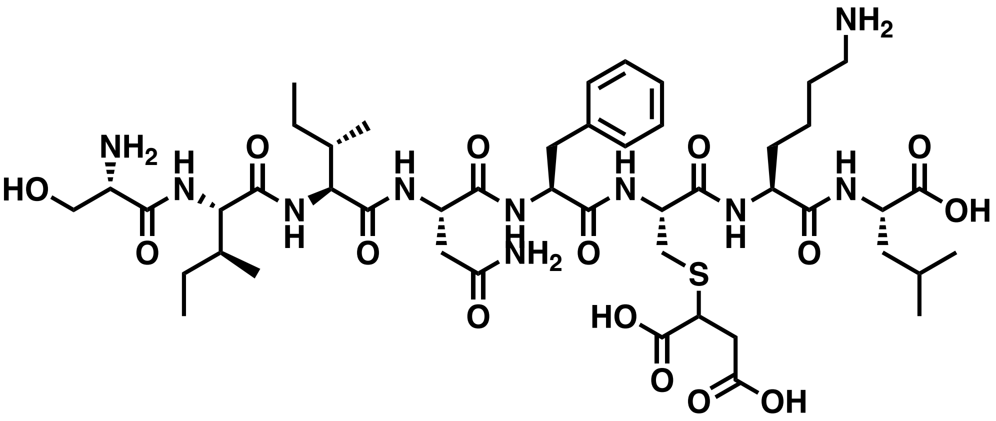


Chemical structure of **ovaE6C$_{2sc}$**.

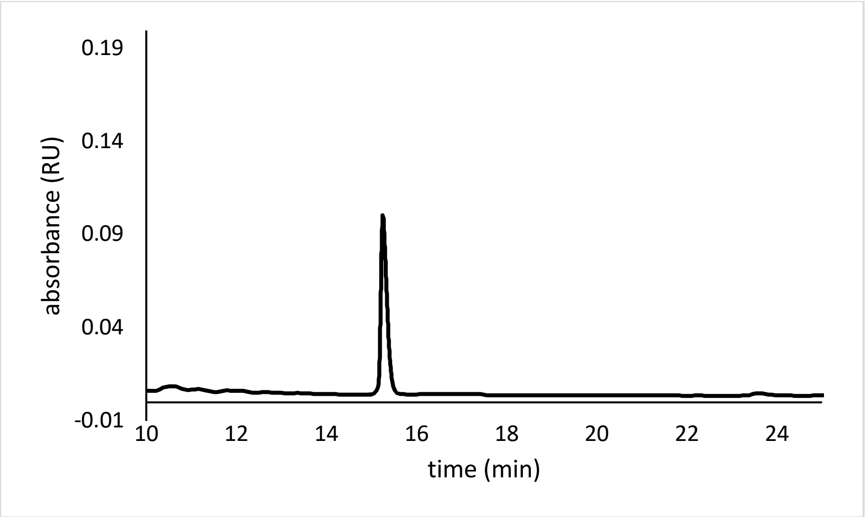


Analytical HPLC Chromatogram of **ovaE6C$_{2sc}$**.

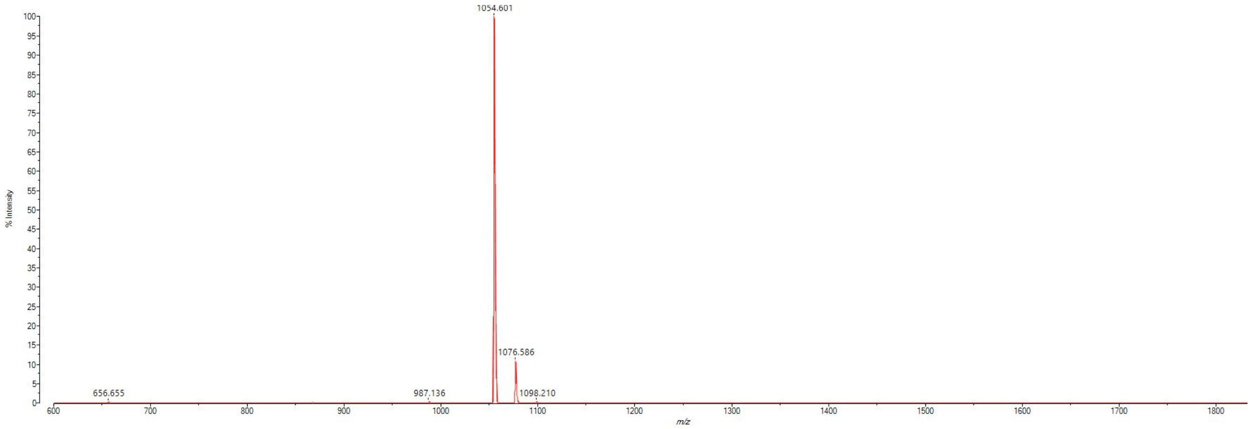


MALDI-TOF mass spectrum for **ovaE6C$_{2sc}$** (m/z 1053.529 found 1054.601).

**ovaN4D** (SIIDFEKL)

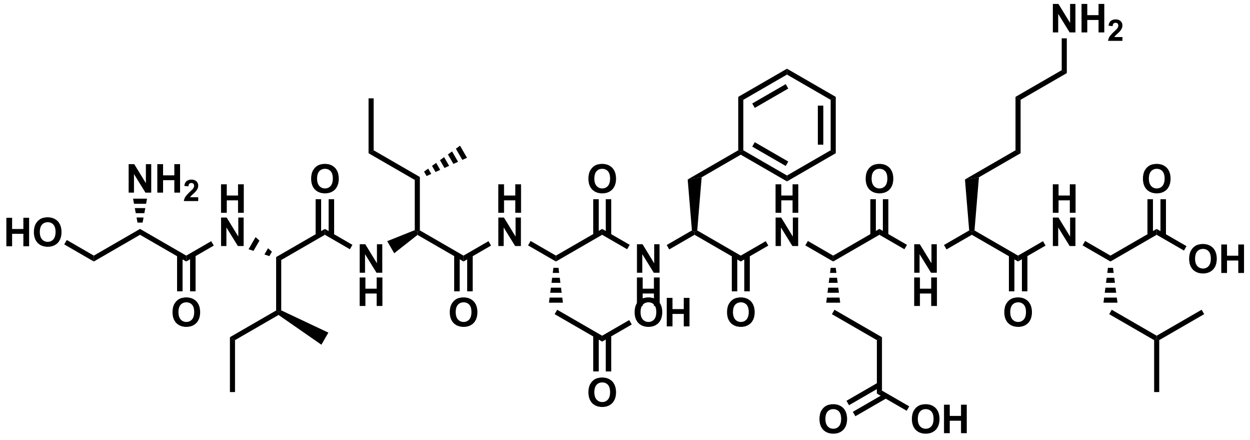


Chemical structure of **ovaN4D**.

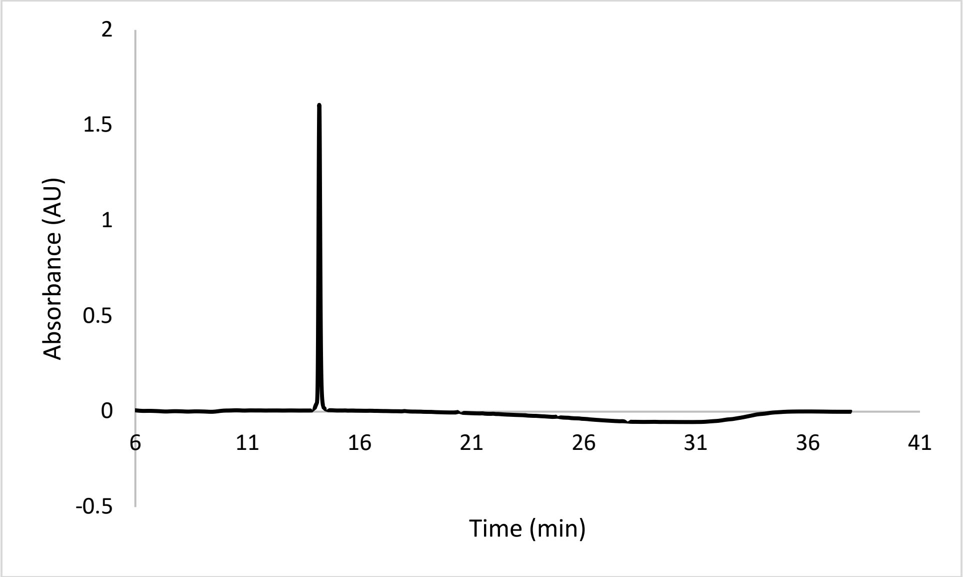


Analytical HPLC Chromatogram of **ovaN4D**.

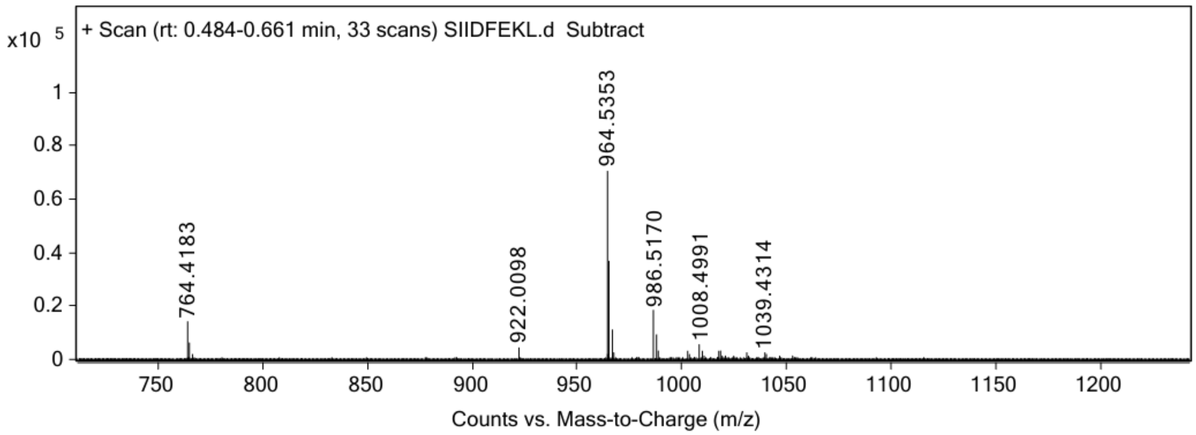


ESI-MS for **ovaN4D** (calculated [M+H$^+$]: 964.5350, found 964.5353).

**ovaN4D$_{iso}$**

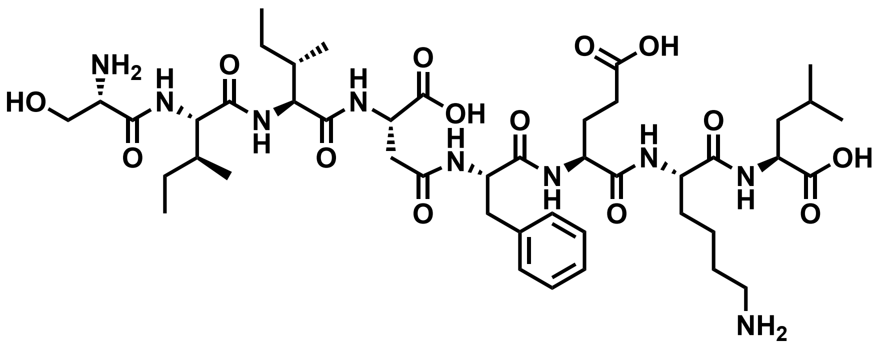

Chemical structure of **ovaN4D$_{iso}$**.

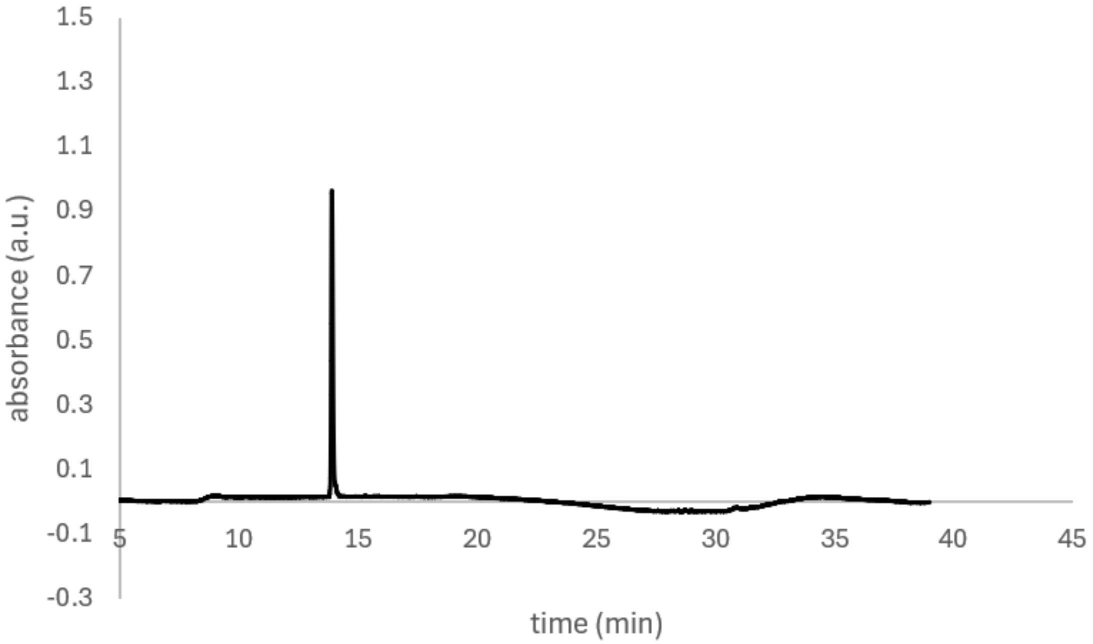


Analytical HPLC Chromatogram of **ovaN4D$_{iso}$**.

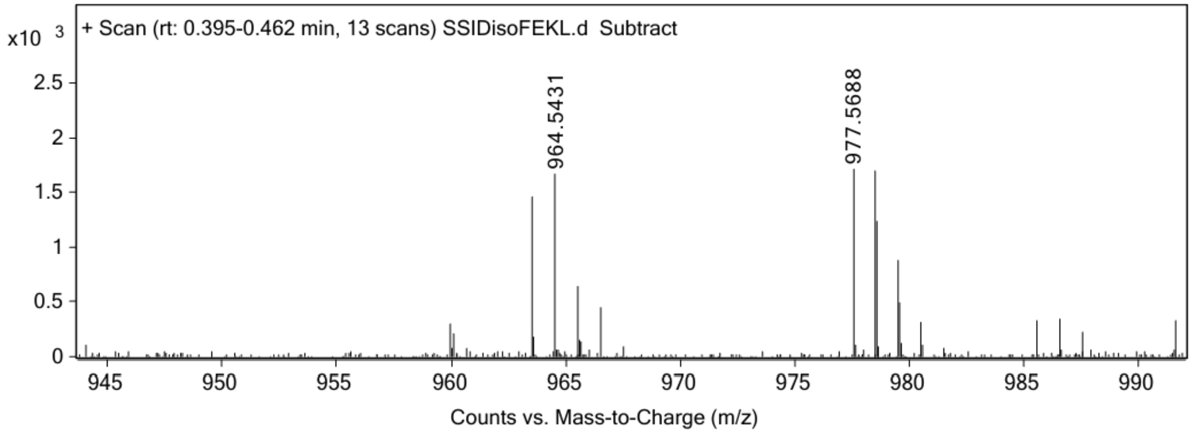


ESI-MS for **ovaN4D$_{iso}$** (calculated [M+H$^+$]: 964.5350, found 964.5431).

**ovaF5Y** (SIINYEKL)

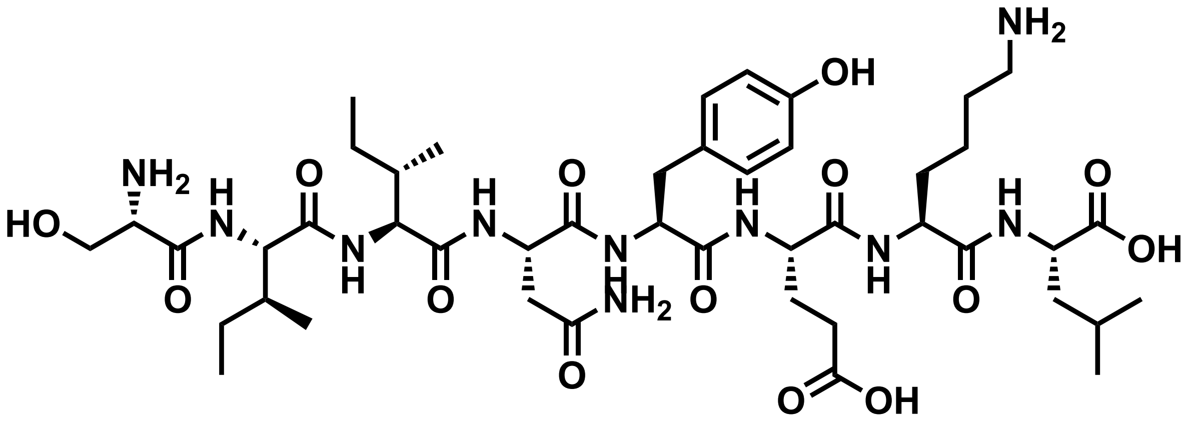


Chemical structure of **ovaF5Y**.

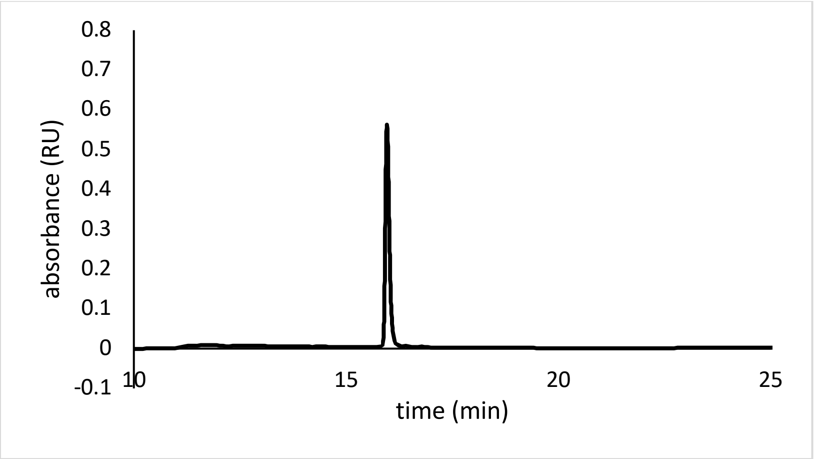


Analytical HPLC Chromatogram of **ovaF5Y.**

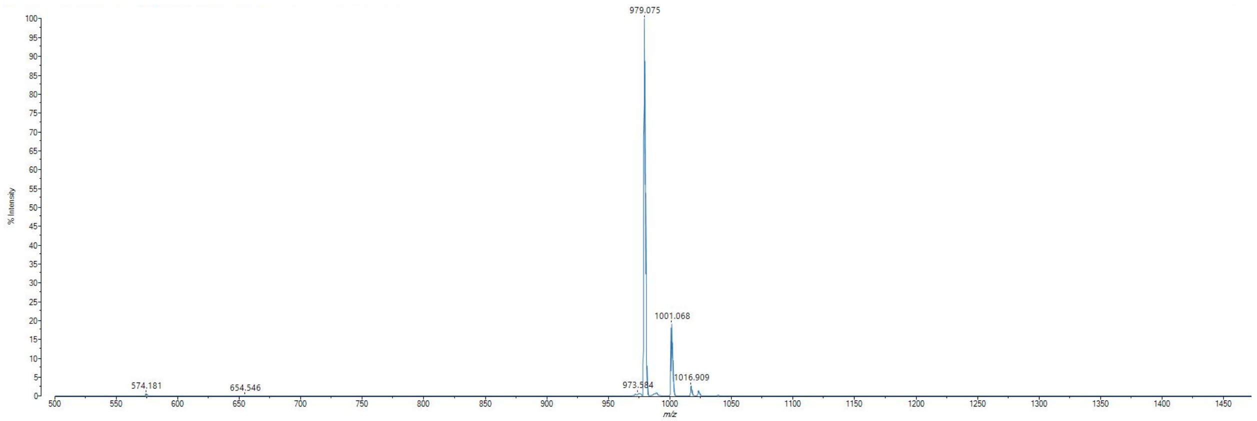


MALDI-TOF mass spectrum for **ovaF5Y** (m/z 979.546 found 979.075).

**ovaF5Y$_{NO2}$**

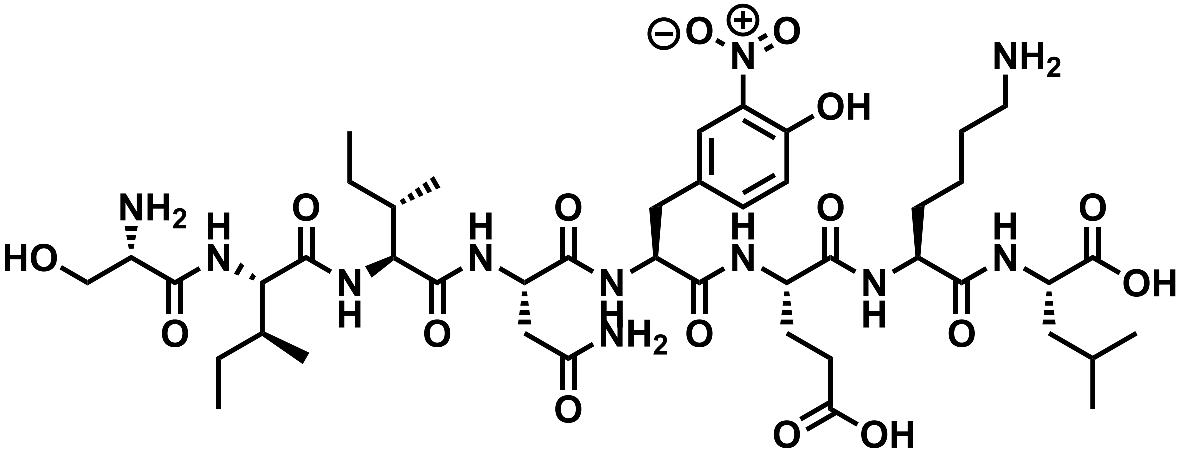

Chemical structure of **ovaF5Y$_{NO2}$**.

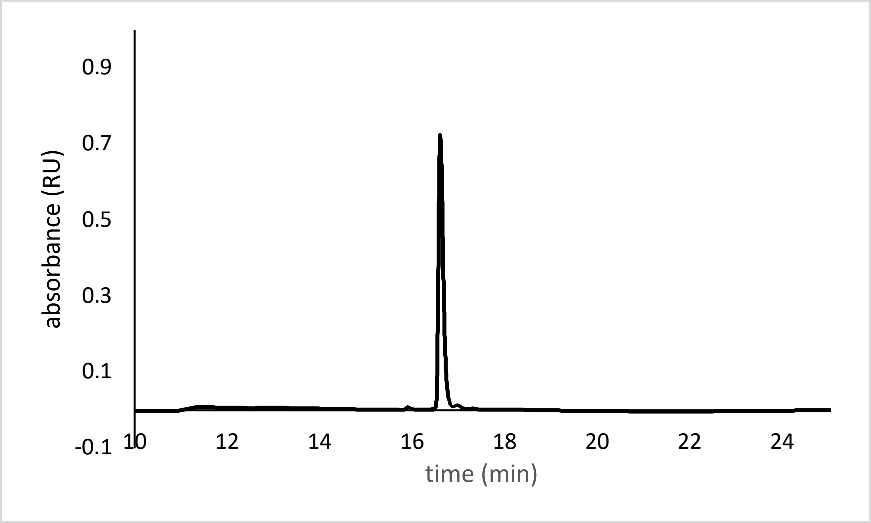


Analytical HPLC Chromatogram of **ovaF5Y$_{NO2}$.**

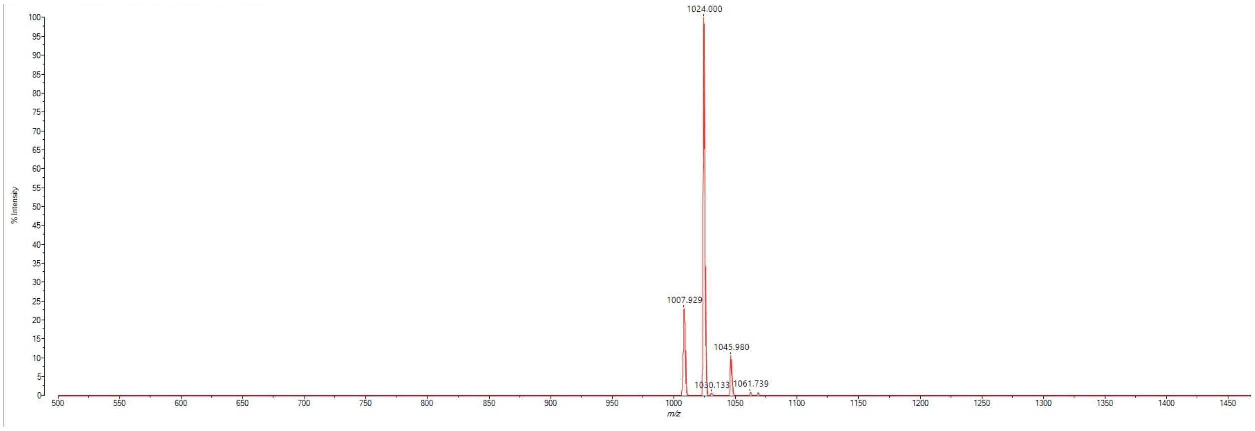


MALDI-TOF mass spectrum for **ovaF5Y$_{NO2}$** (m/z 1024.531 found 1024.000).

**ovaF5Y$_{Cl}$**

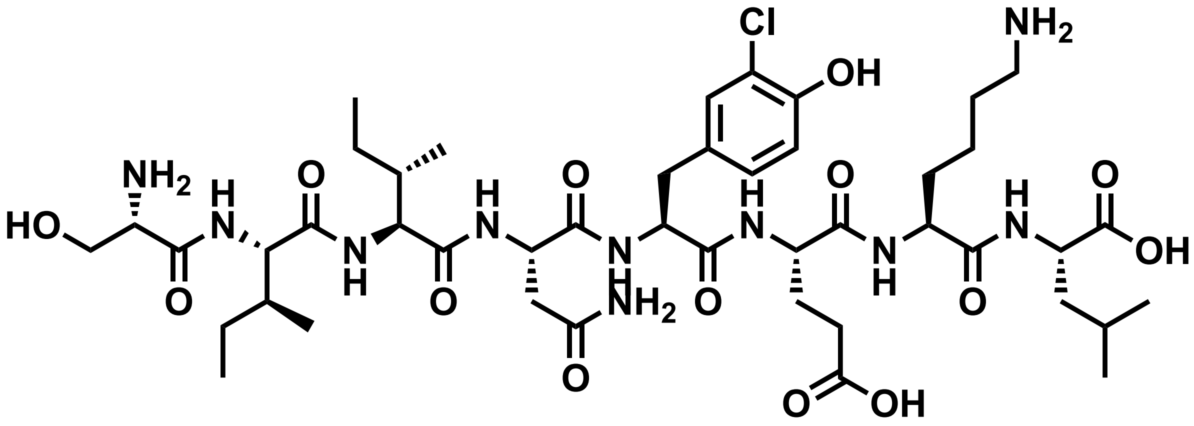


Chemical structure of **ovaF5Y$_{Cl}$**.

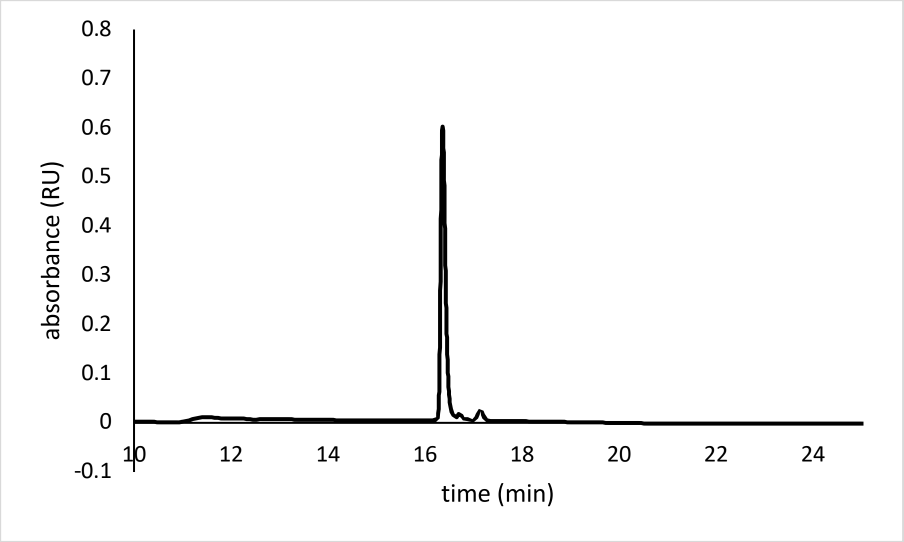


Analytical HPLC Chromatogram of **ovaF5Y$_{Cl}$.**

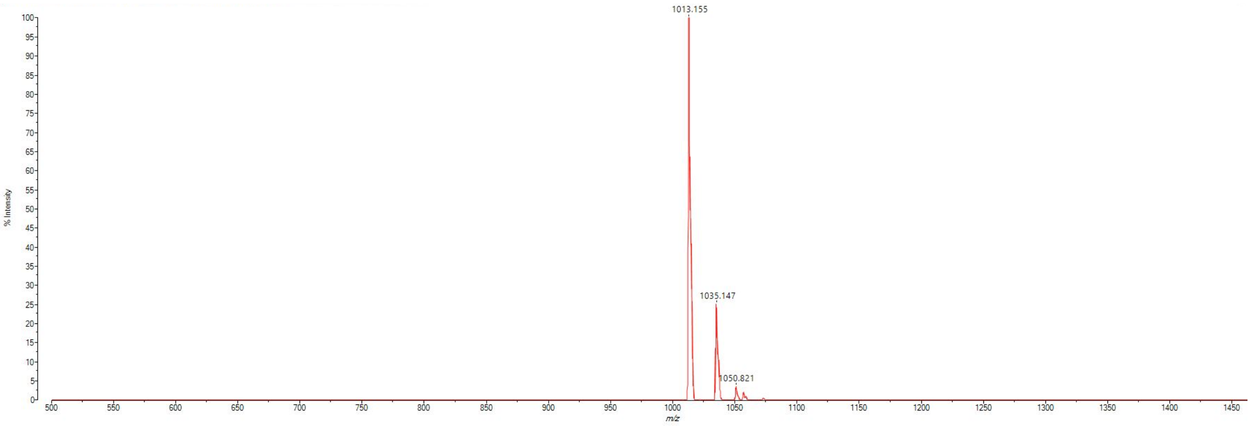


MALDI-TOF mass spectrum for **ovaF5Y$_{Cl}$** (m/z 1013.507 found 1013.155).

**ovaK7$_{CTN}$**

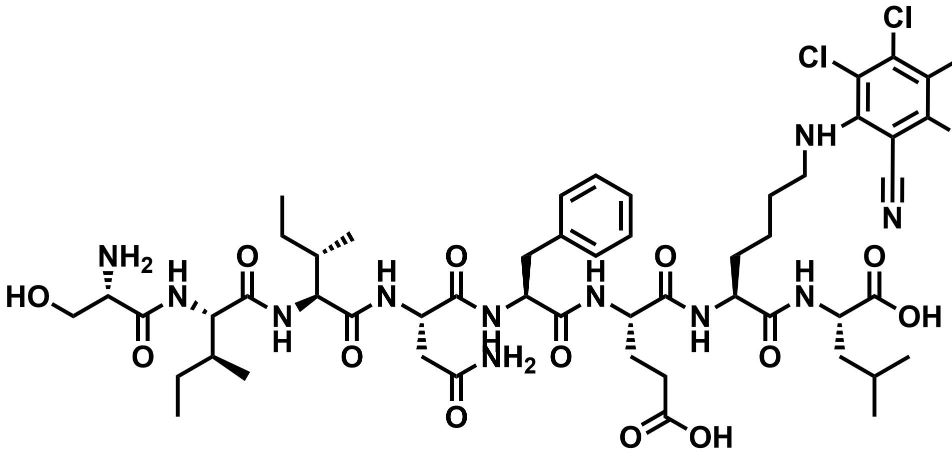

Chemical structure of **ovaK7$_{CTN}$**.

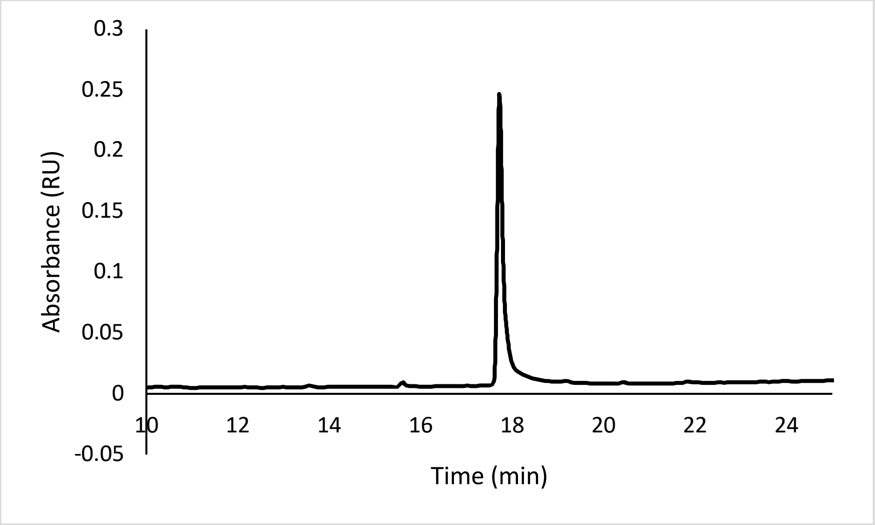


Analytical HPLC Chromatogram of **ovaK7$_{CTN}$.**

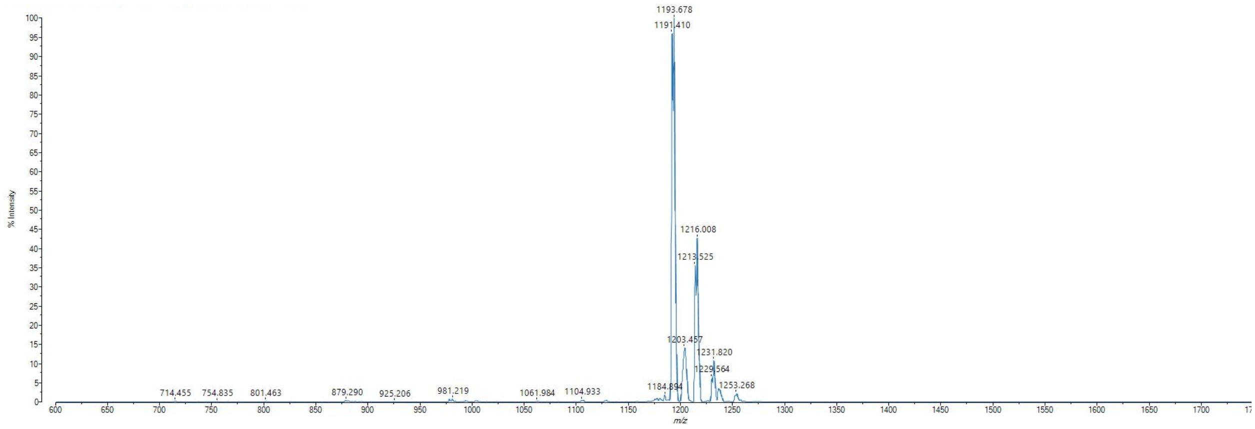


MALDI-TOF mass spectrum for **ovaK7CTN** (m/z 1191.456 found 1191.410).

**ovaK7$_{ASA}$**

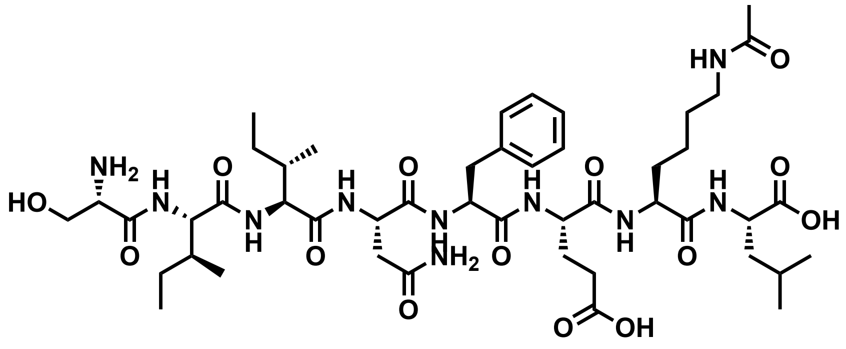


Chemical structure of **ovaK7$_{ASA}$**.

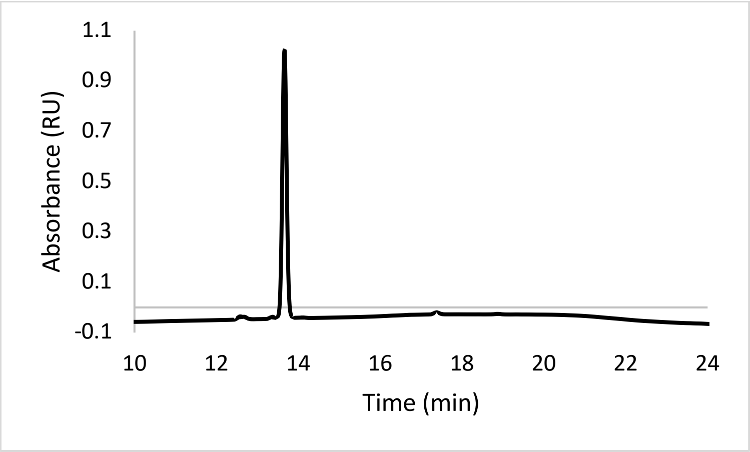


Analytical HPLC Chromatogram of **ovaK7$_{ASA}$.**

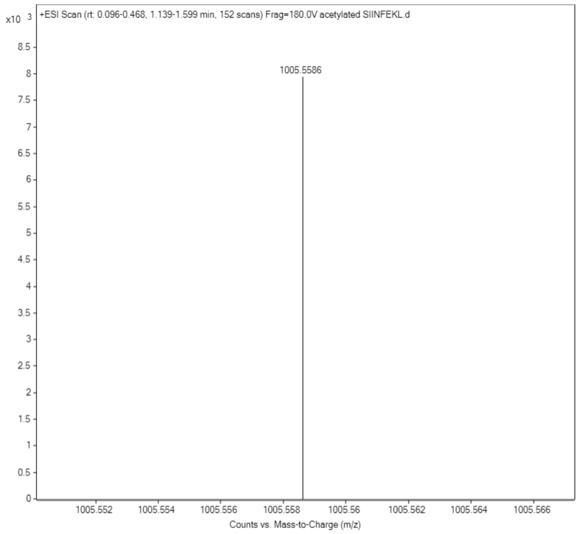


ESI-MS for **ovaK7$_{ASA}$** (calculated [M+H$^+$]: 1005.5620, found 1005.5586)

**ovaK7$_{CFX}$**

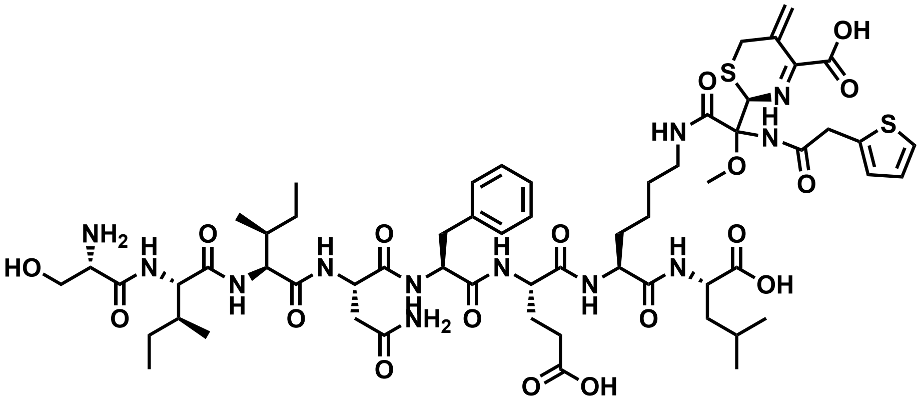

Chemical structure of **ovaK7$_{CFX}$**.

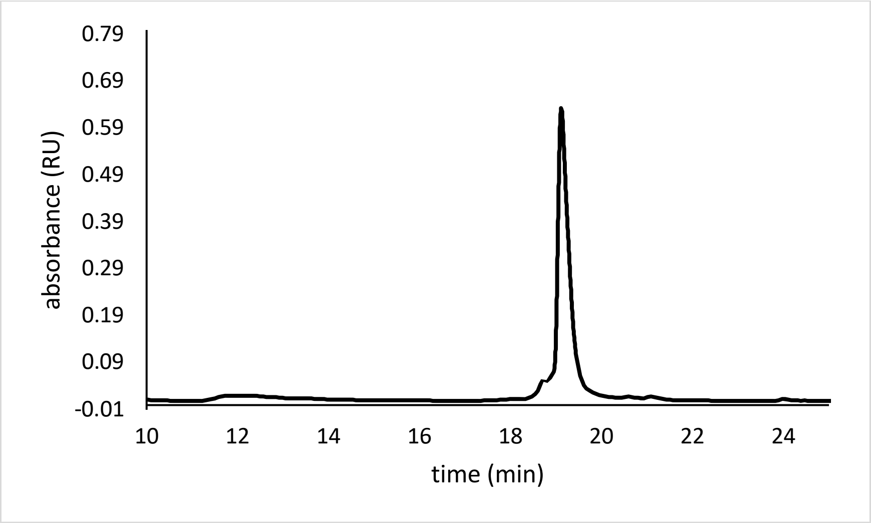


Analytical HPLC Chromatogram of **ovaK7$_{CFX}$.**

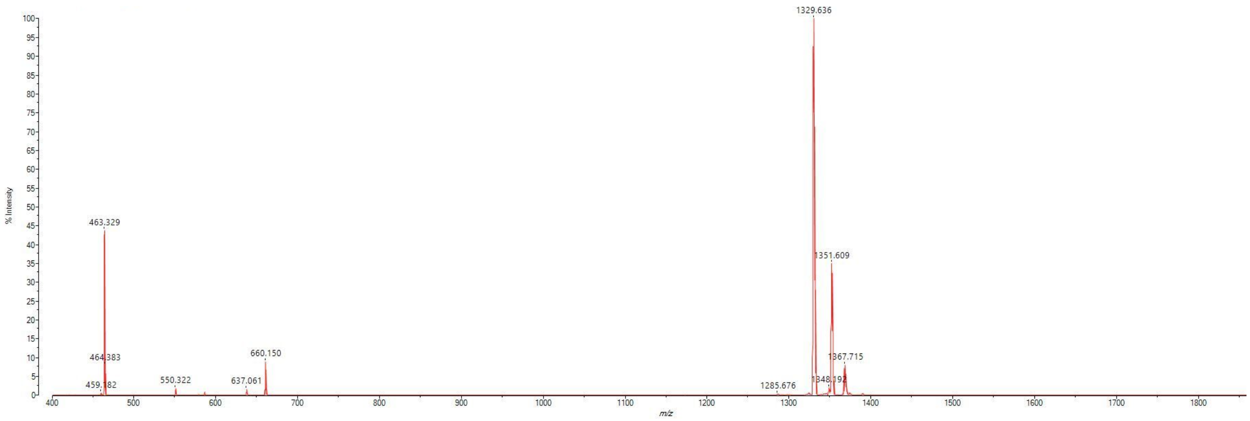


MALDI-TOF mass spectrum for **ovaK7$_{CFX}$** (m/z 1329.585 found 1329.636).

**ovaK7$_{PEITC}$**

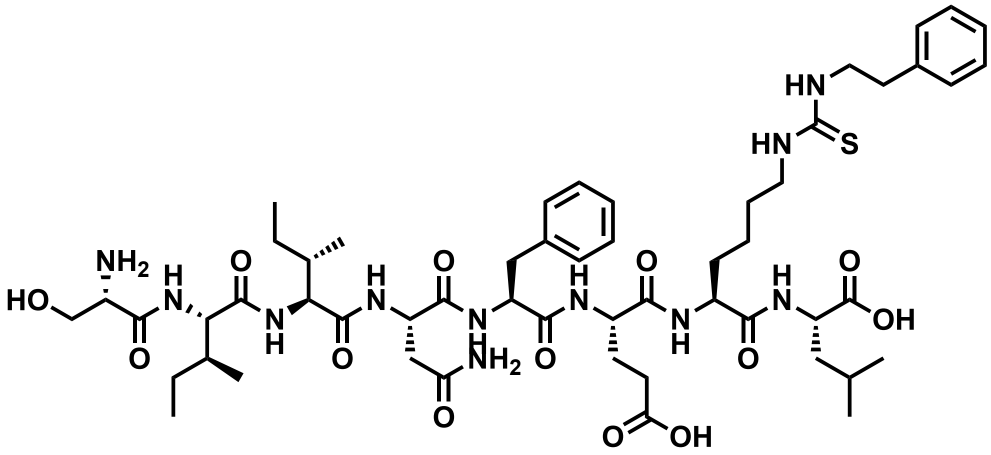


Chemical structure of **ovaK7$_{PEITC}$**

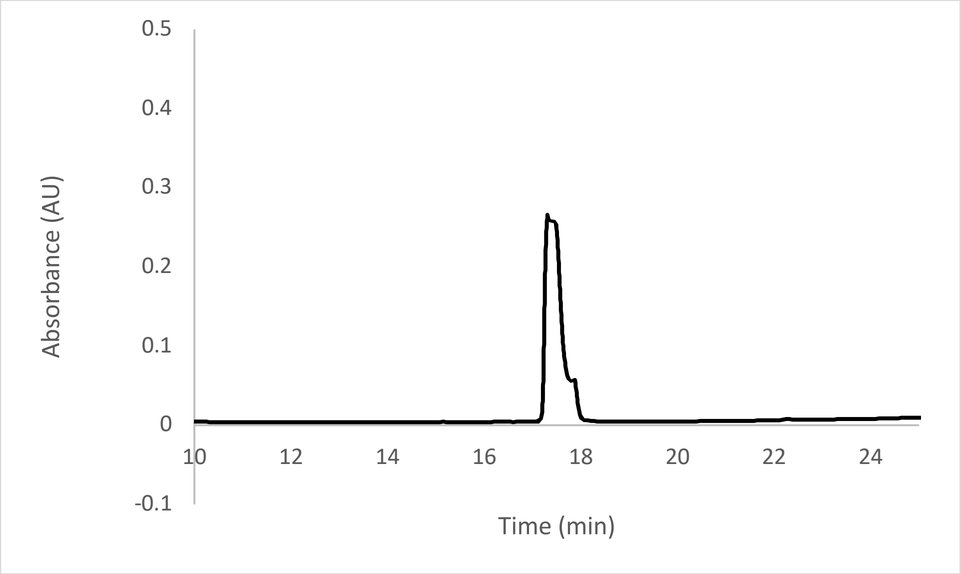


Analytical HPLC Chromatogram of **ovaK7$_{PEITC}$**

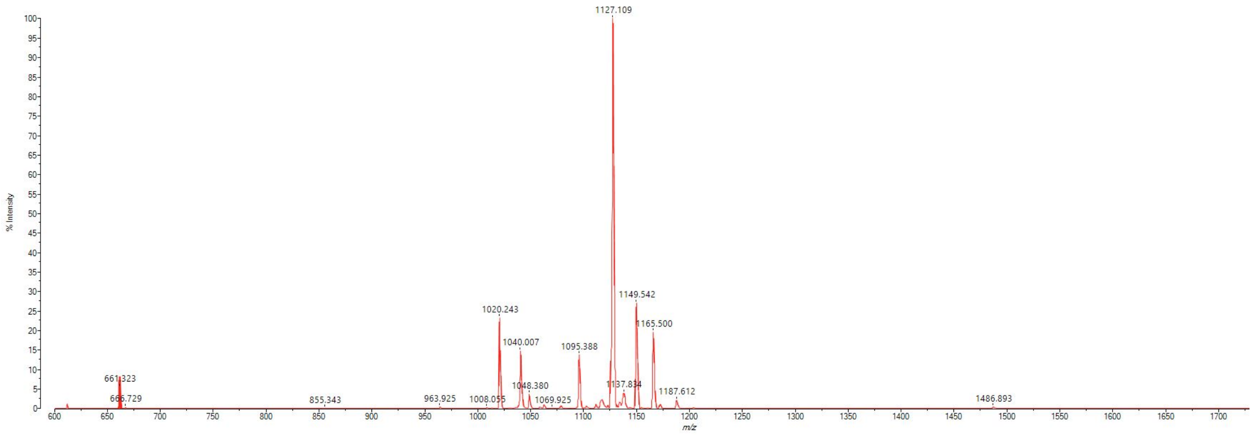


MALDI-TOF mass spectrum for **ovaK7$_{PEITC}$** (m/z 1126.597 found 1127.109)

**ovaK7$_{AITC}$**

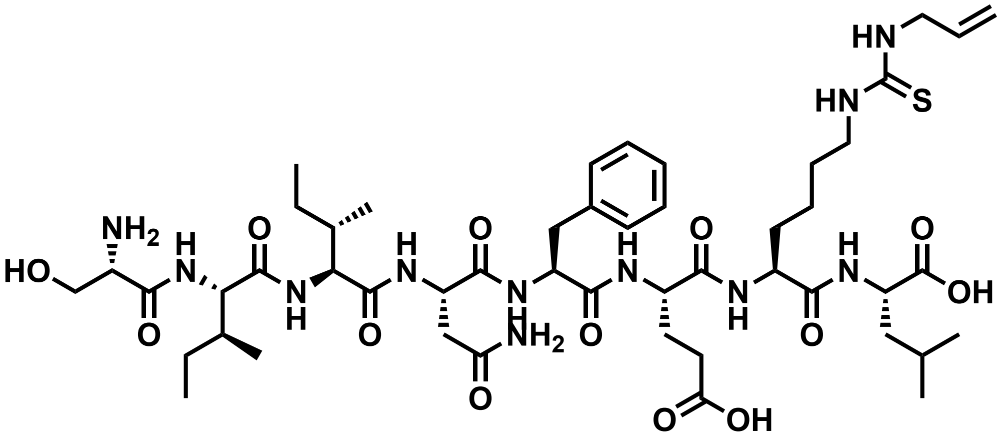

Chemical structure of **ovaK7$_{AITC}$**

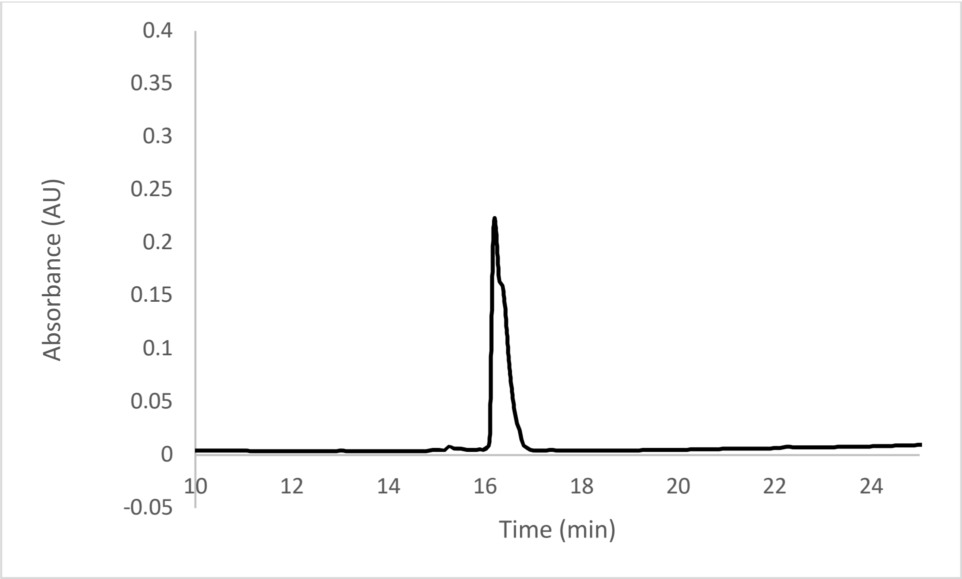


Analytical HPLC Chromatogram of **ovaK7$_{AITC}$**

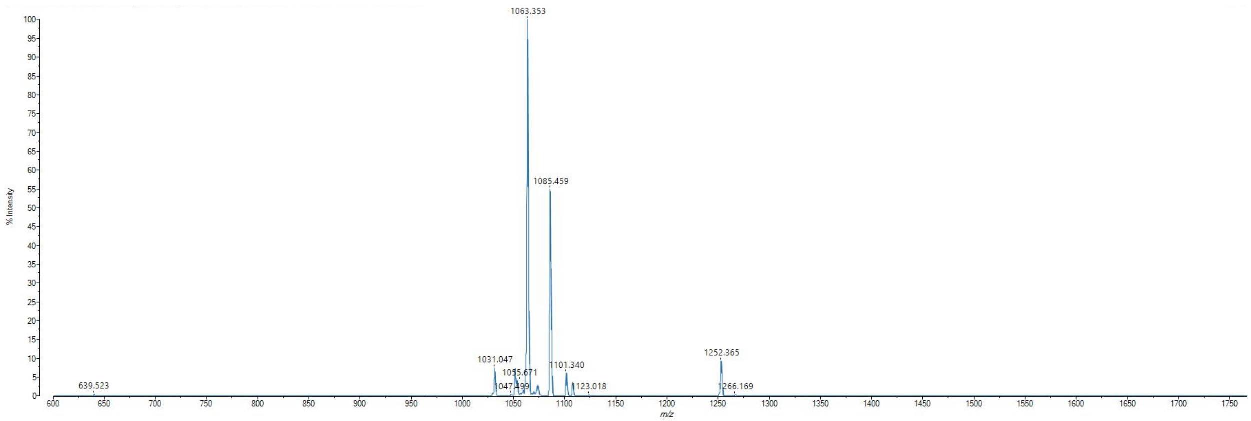


MALDI-TOF mass spectrum for **ovaK7$_{AITC}$** (m/z 1062.565 found 1063.353)

**ovaK7C** (SIINFECL)

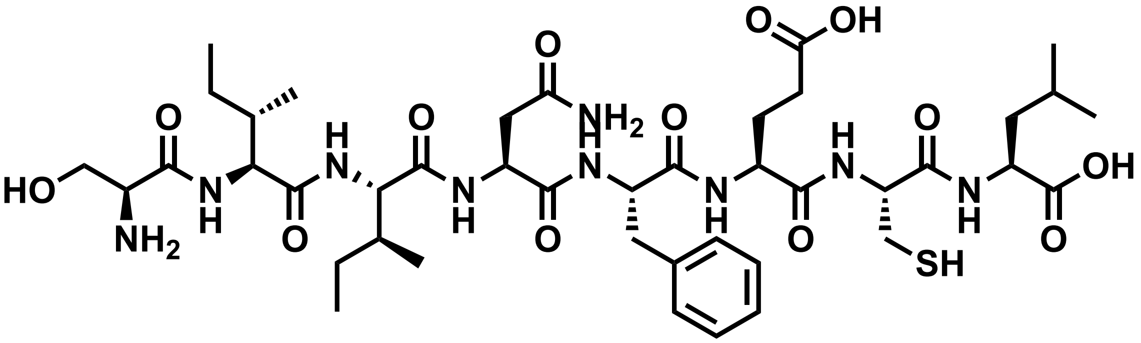

Chemical structure of **ovaK7C**

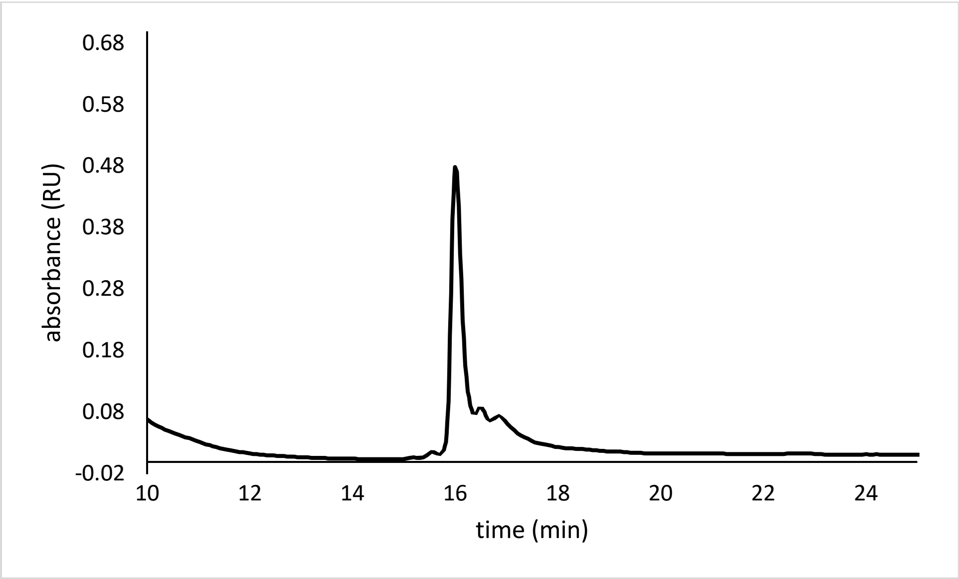


Analytical HPLC Chromatogram of **ovaK7C**

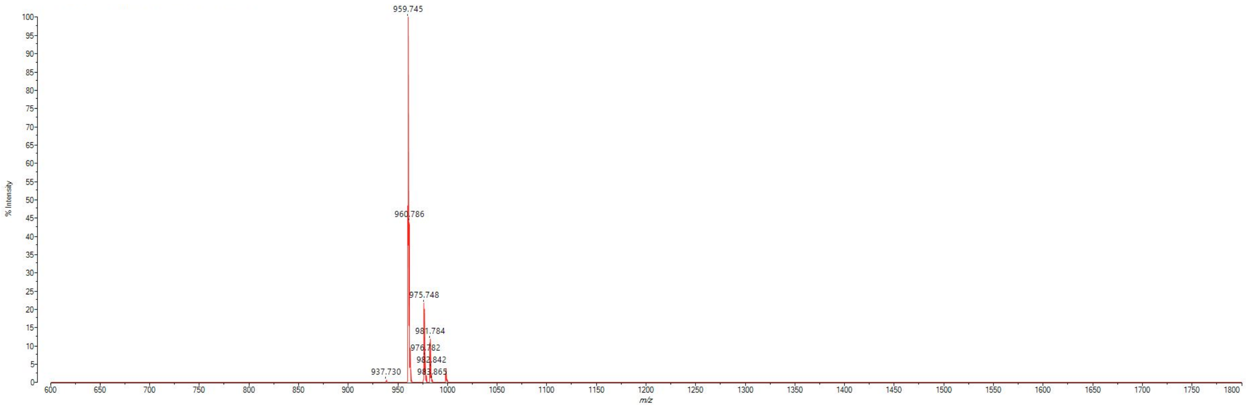


MALDI-TOF mass spectrum for **ovaK7C** (m/z 938.465 found 937.730)

**ovaK7C$_{PEITC}$**

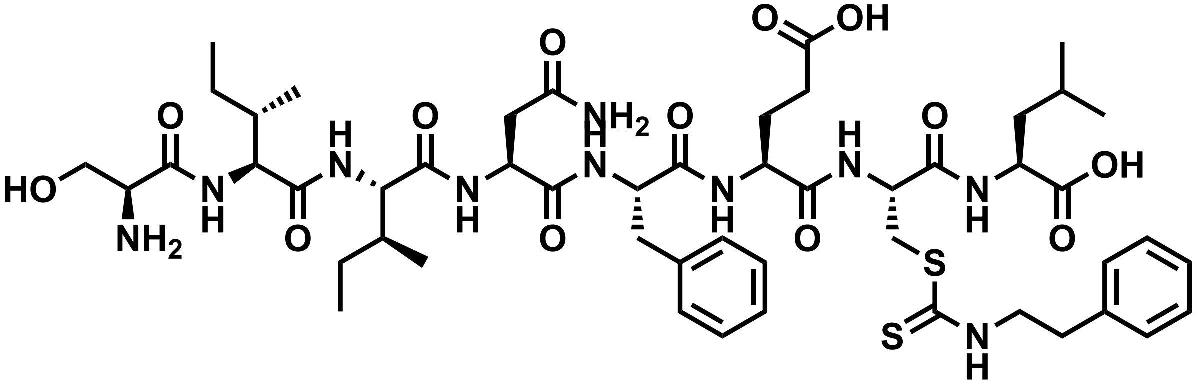

Chemical structure of **ovaK7C$_{PEITC}$**

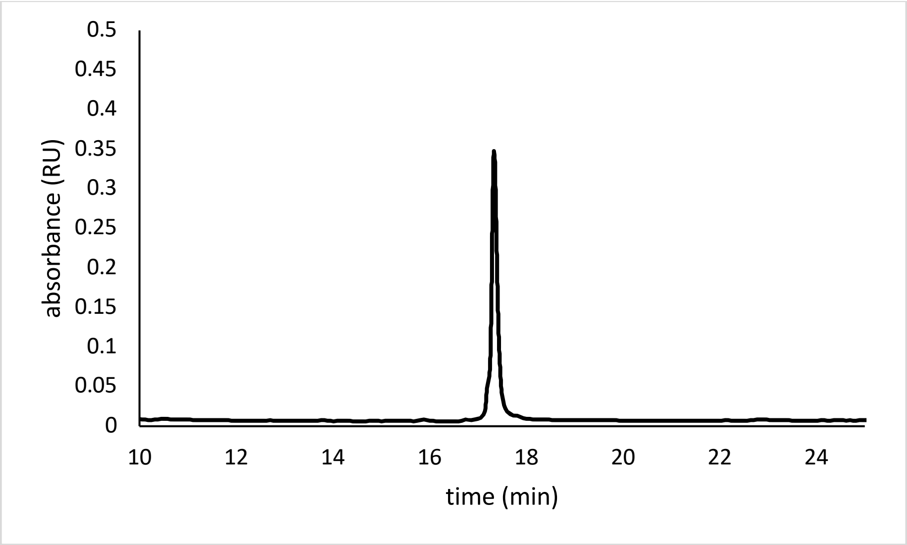


Analytical HPLC Chromatogram of **ovaK7C$_{PEITC}$**

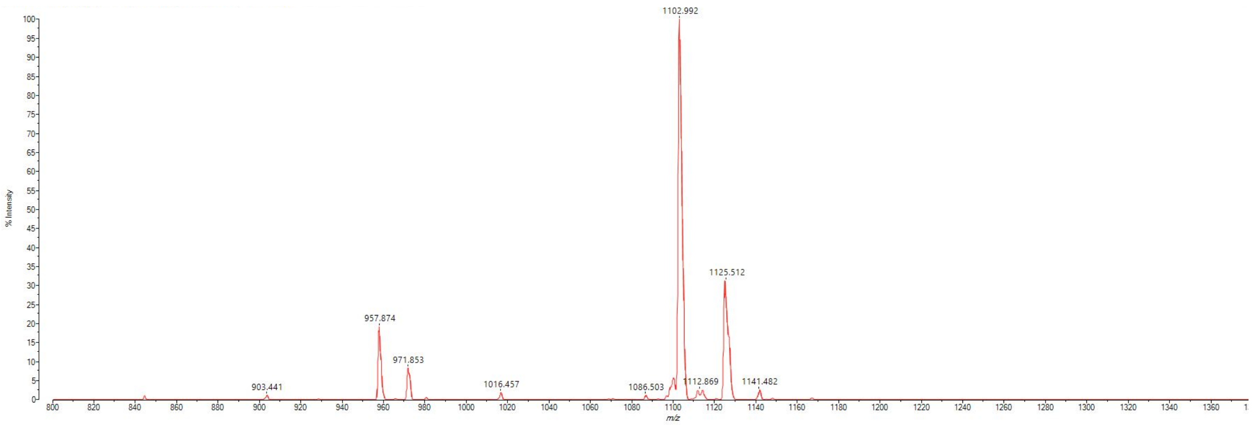


MALDI-TOF mass spectrum for **ovaK7C$_{PEITC}$** (m/z 1101.511 found 1102.992)

**ovaK7C$_{\mathbf{BrEA}}$**

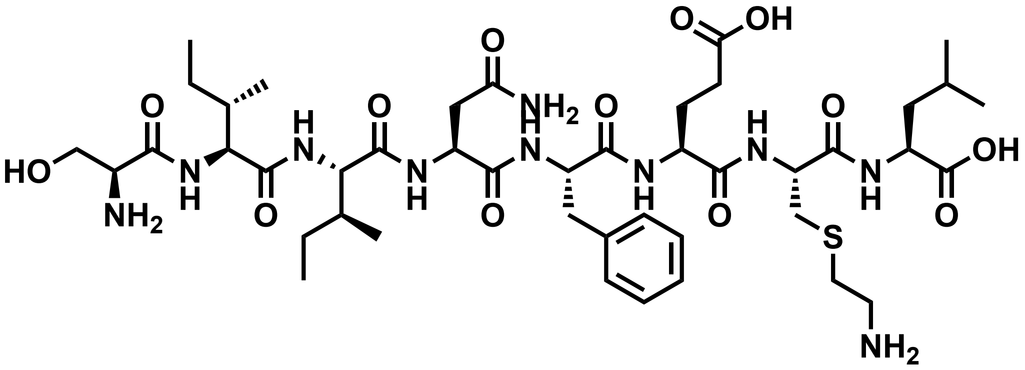

Chemical structure of **ovaK7C$_{\mathbf{BrEA}}$**

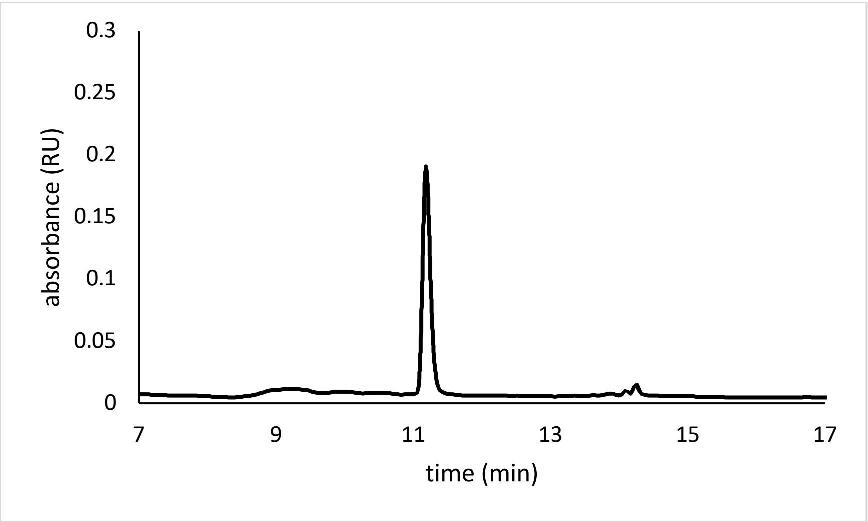


Analytical HPLC Chromatogram of **ovaK7C$_{\mathbf{BrEA}}$**

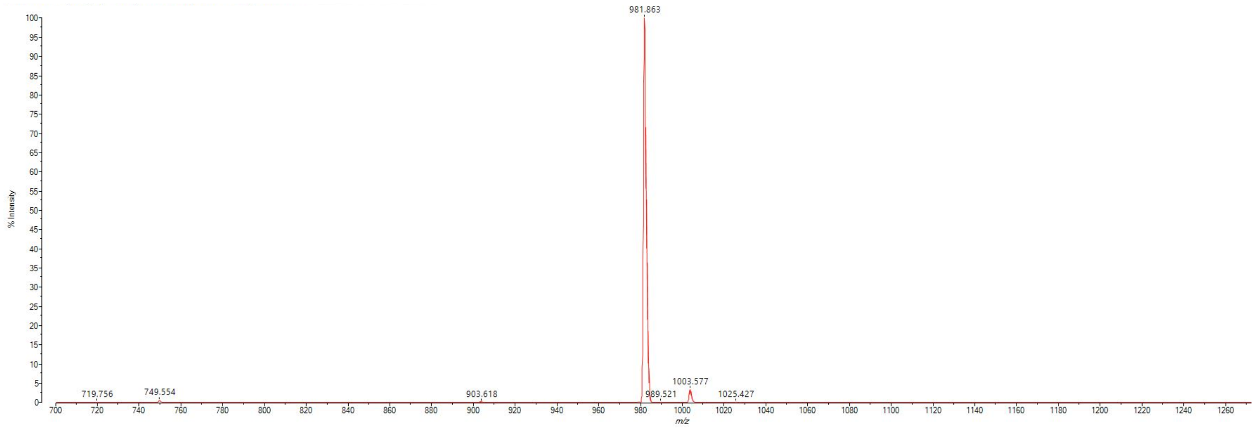


MALDI-TOF mass spectrum for **ovaK7C$_{\mathbf{BrEA}}$** (m/z 981.507 found 981.863)

**ovaK7C$_{\mathbf{CMQ}}$**

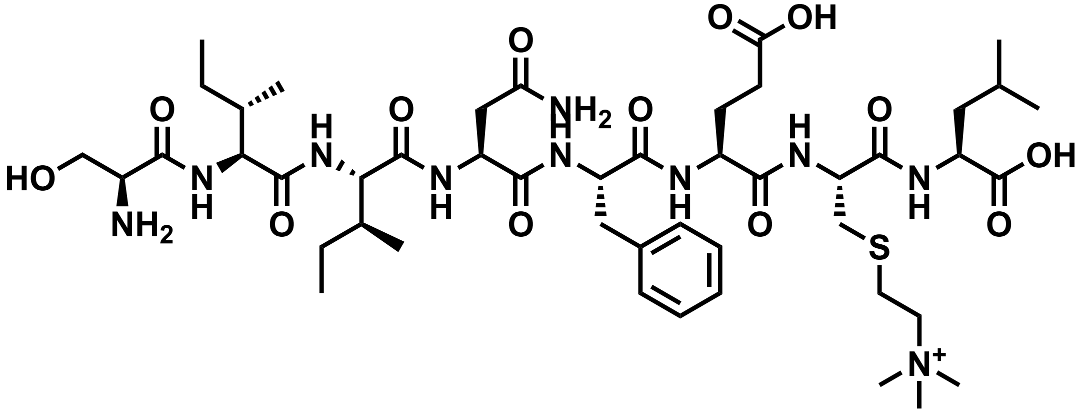

Chemical structure of **ovaK7C$_{\mathbf{CMQ}}$**

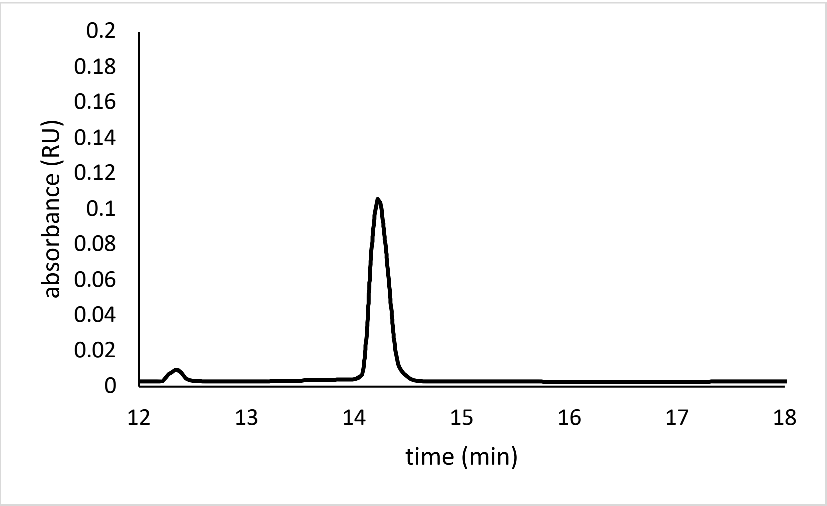


Analytical HPLC Chromatogram of **ovaK7C$_{\mathbf{CMQ}}$**

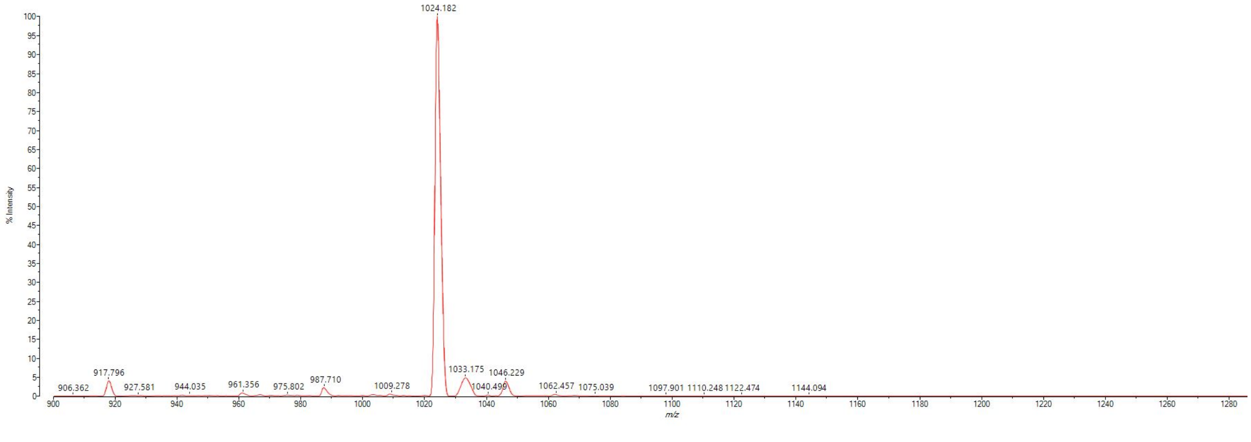


MALDI-TOF mass spectrum for **ovaK7C$_{\mathbf{CMQ}}$** (m/z 1024.562 found 1024.182)

**ovaK7R** (SIINFERL)

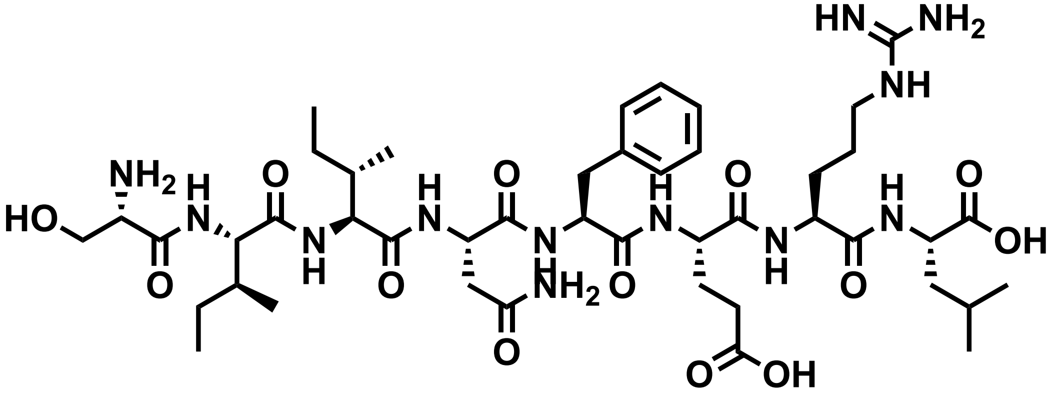


Chemical structure of **ovaK7R**

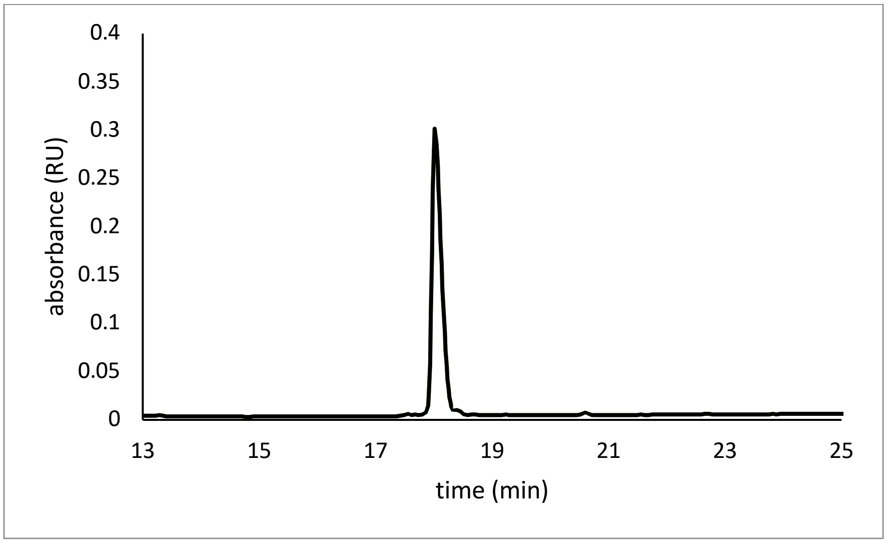


Analytical HPLC Chromatogram of **ovaK7R**

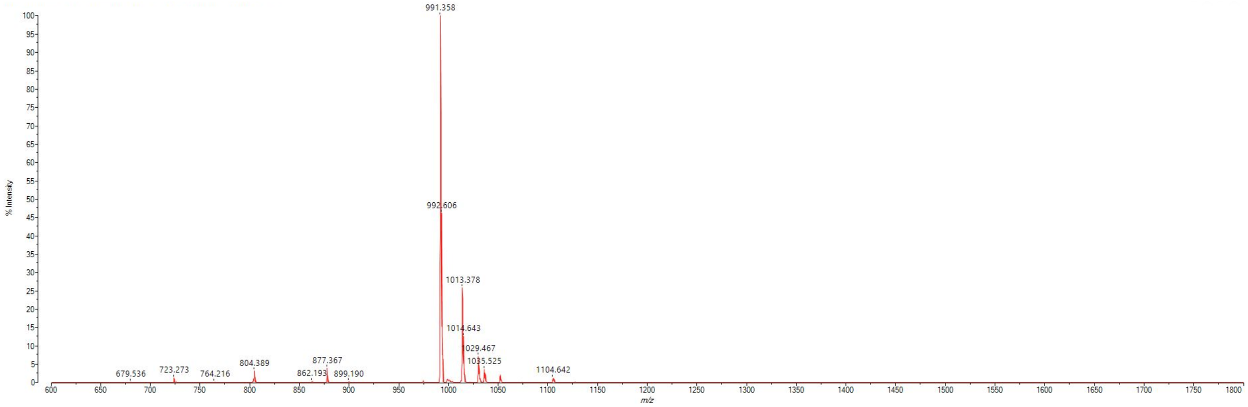

MALDI-TOF mass spectrum for **ovaK7R** (m/z 991.557 found 991.358)
**ca1 (KGMNYTVRL)**

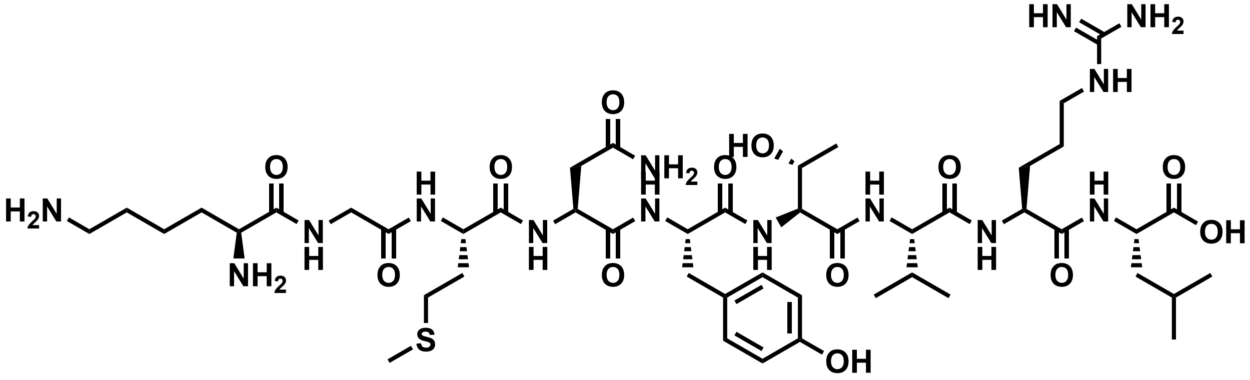


Chemical structure of **ca1**.

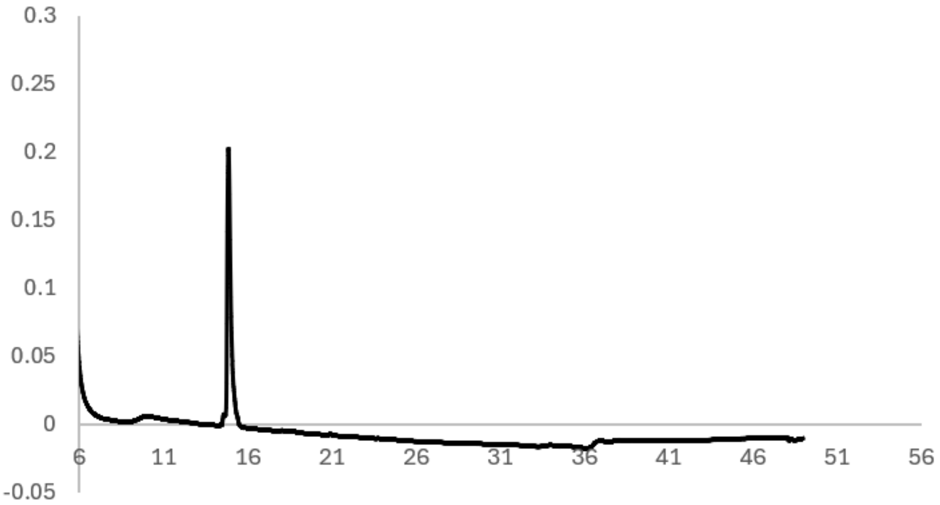


Analytical HPLC Chromatogram of **ca1**.

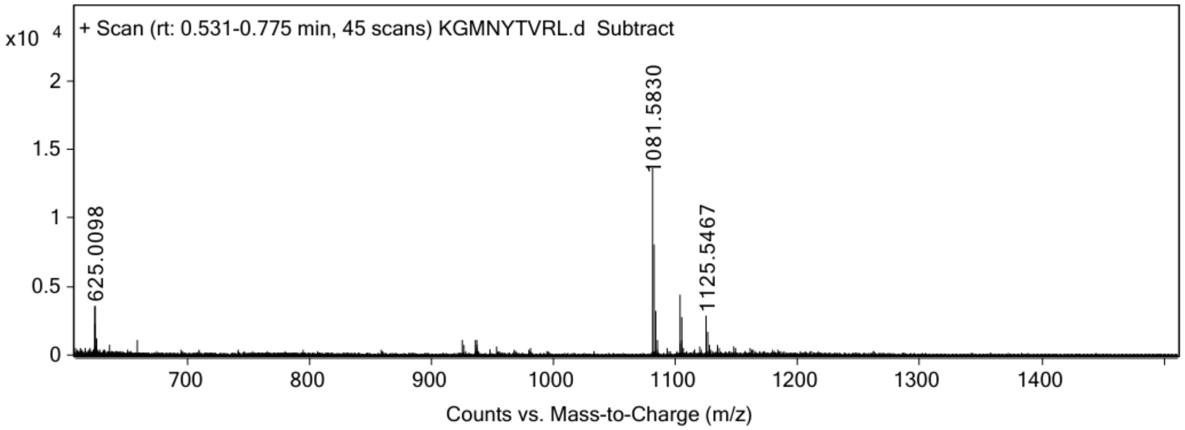


ESI-MS for **ca1** (calculated [M+H$^+$]: 1081.5823, found 1081.5830).

**ca1$_{oxide}$**

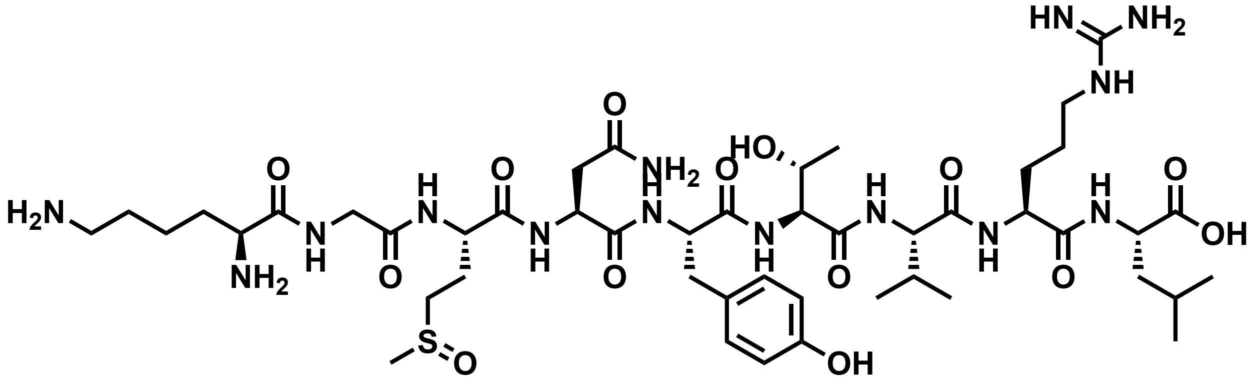


Chemical structure of **ca1$_{oxide}$.**

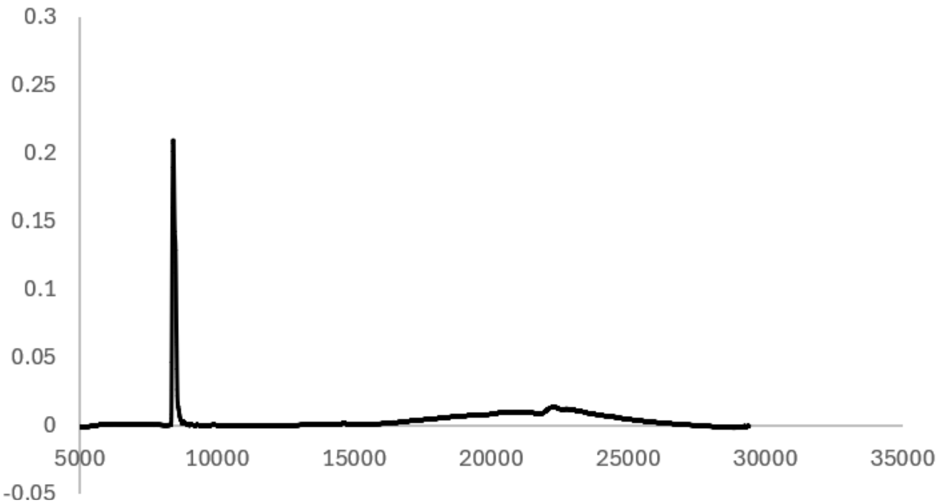


Analytical HPLC Chromatogram of **ca1$_{oxide}$.**

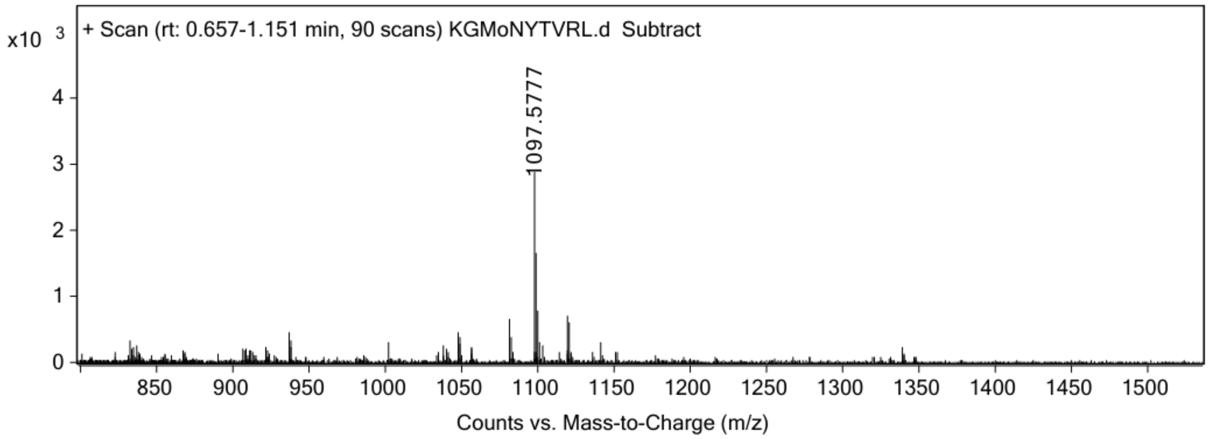


ESI-MS for **ca1$_{oxide}$** (calculated [M+H$^+$]: 1097.5772, found 1097.5777).

**ca1$_{oxone}$**

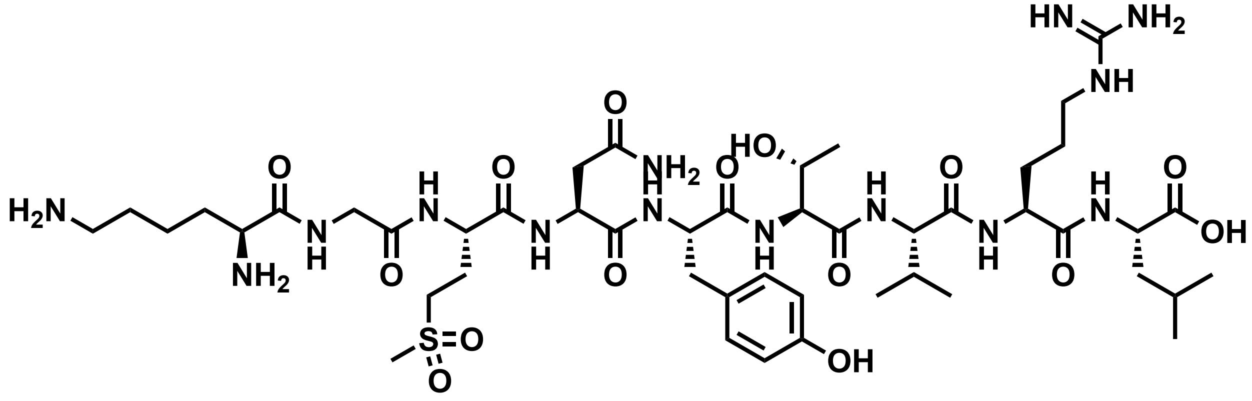

Chemical structure of **ca1$_{oxone}$.**

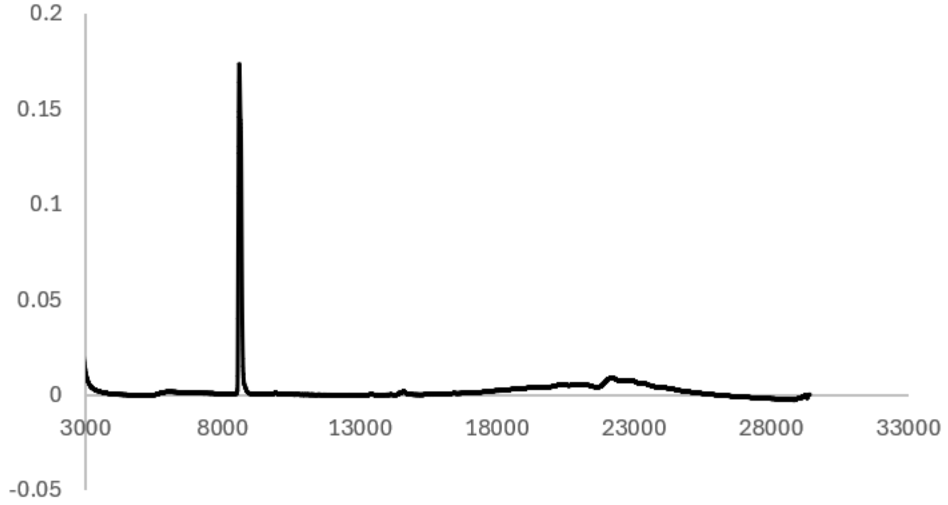


Analytical HPLC Chromatogram of **ca1$_{oxone}$.**

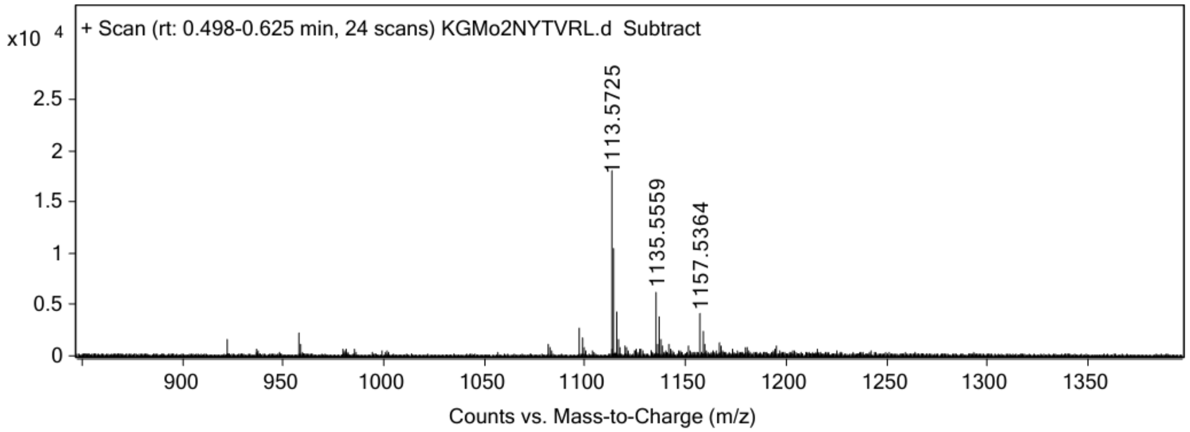


ESI-MS for **ca1$_{oxone}$** (calculated [M+H$^+$]: 1113.5721, found 1113.5725).

**ca2 (SAPENAVRM)**

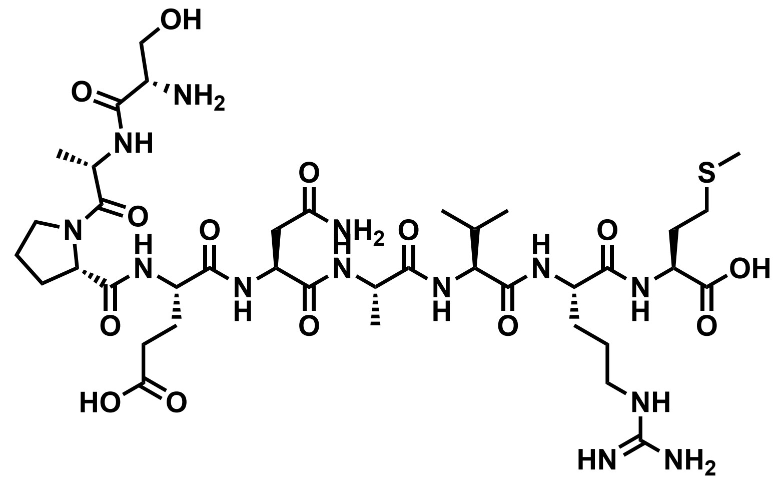


Chemical structure of **ca2**.

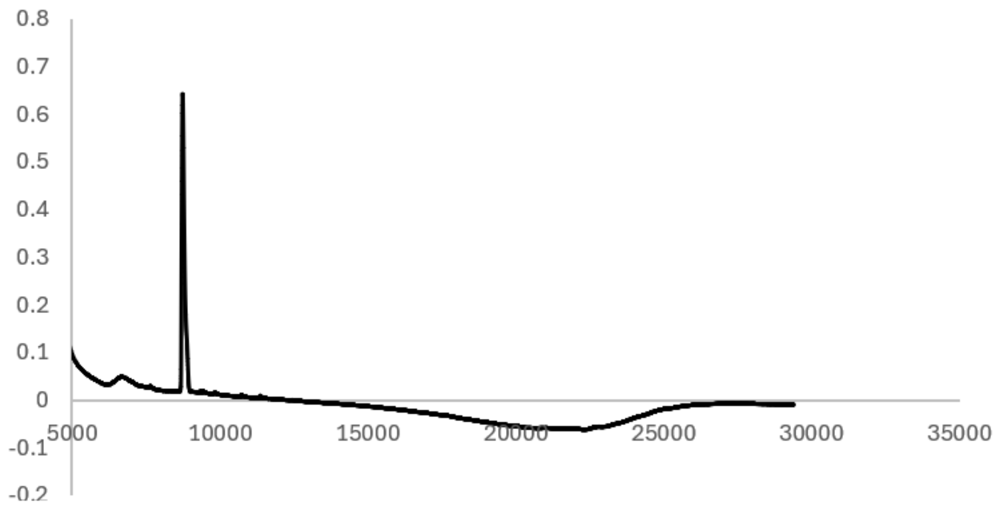


Analytical HPLC Chromatogram of **ca2.**

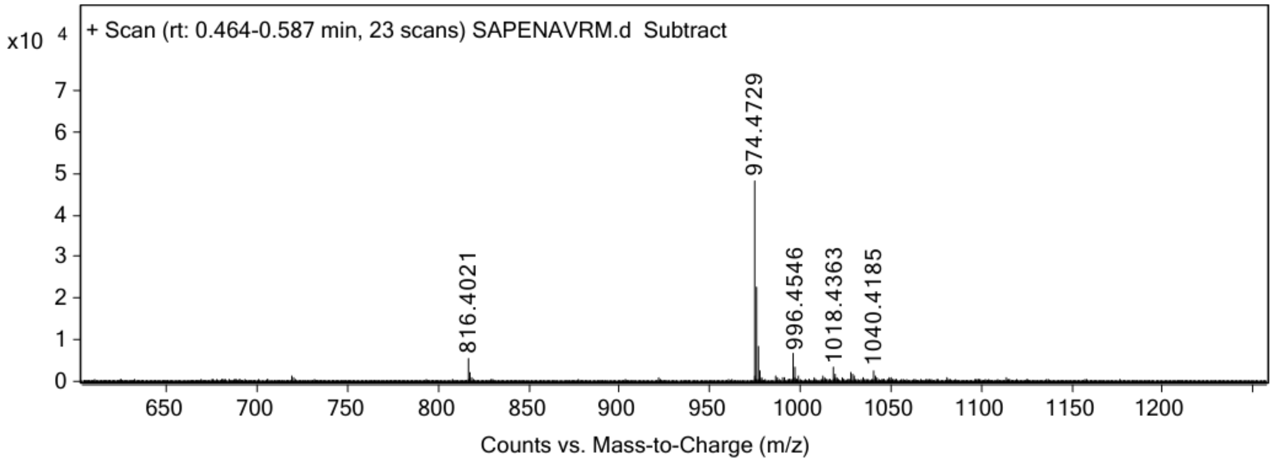


ESI-MS for **ca2** (calculated [M+H$^+$]: 974.4724, found 974.4729).

**ca2$_{oxide}$**

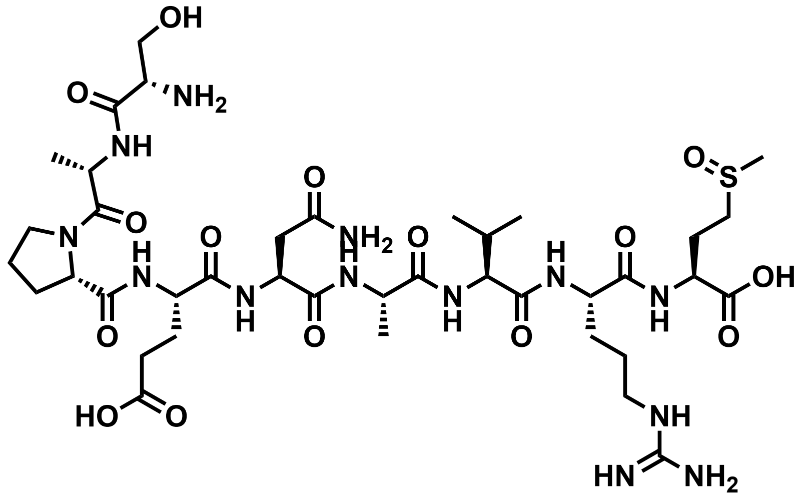


Chemical structure of **ca2$_{oxide}$**.

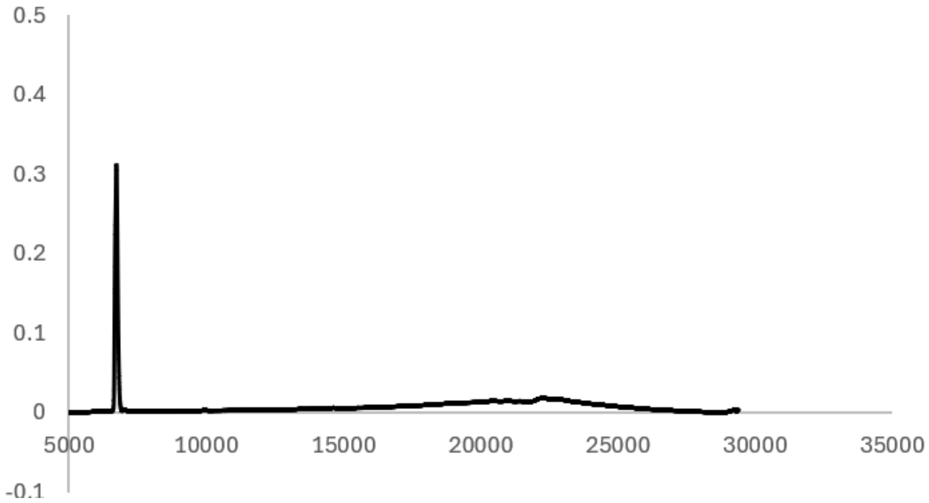


Analytical HPLC Chromatogram of **ca2$_{oxide}$**.

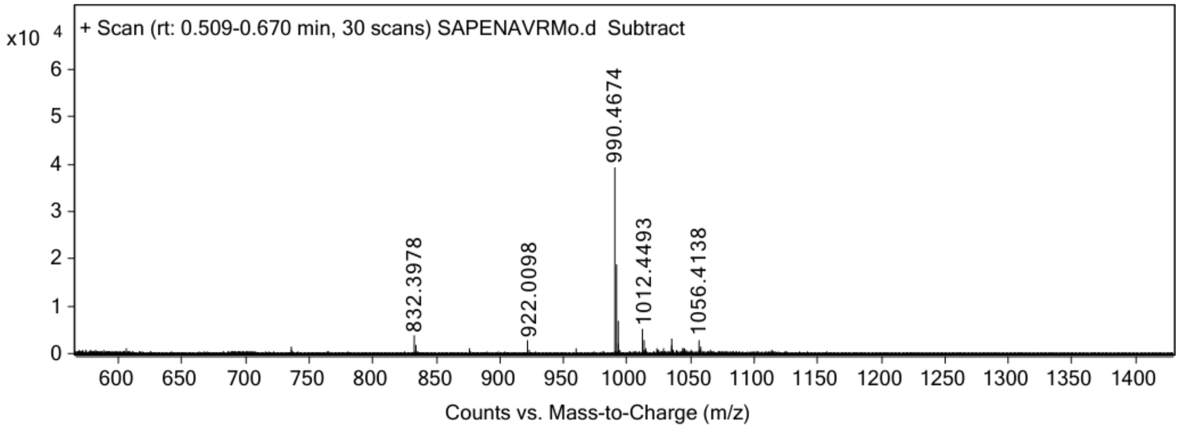


ESI-MS for **ca2$_{oxide}$** (calculated [M+H$^+$]: 990.4673, found 990.4674).

**ca2$_{oxone}$**

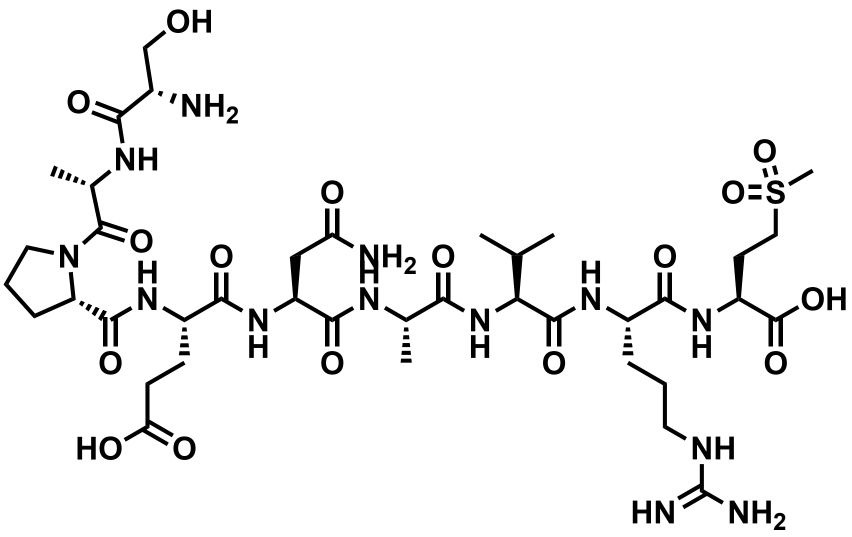


Chemical structure of **ca2$_{oxone}$.**

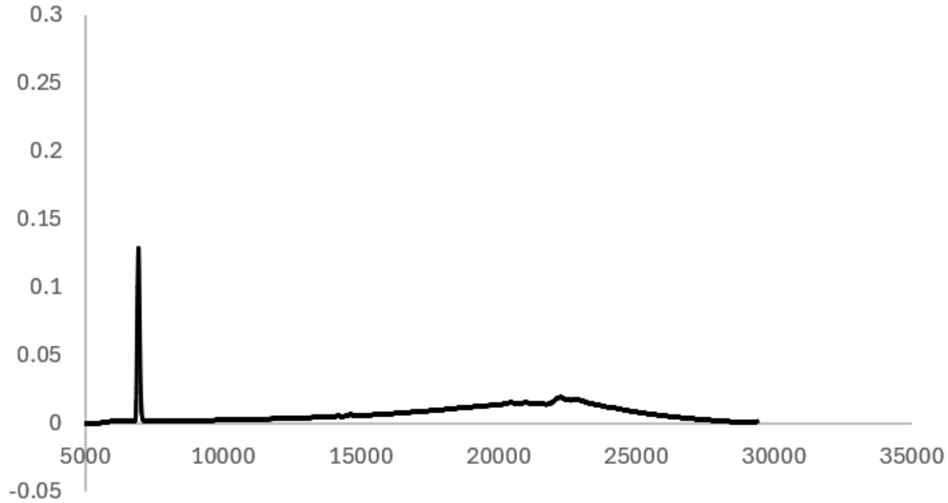


Analytical HPLC Chromatogram of **ca2$_{oxone}$.**

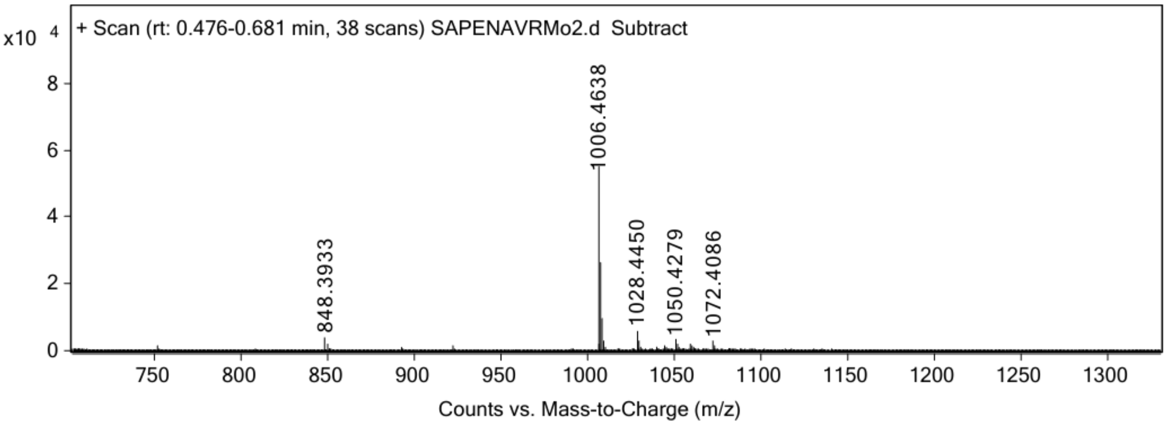


ESI-MS for **ca2$_{oxone}$** (calculated [M+H$^{+}$]: 1006.4622, found 1006.4638).

**ca3 (INFDFPKL)**

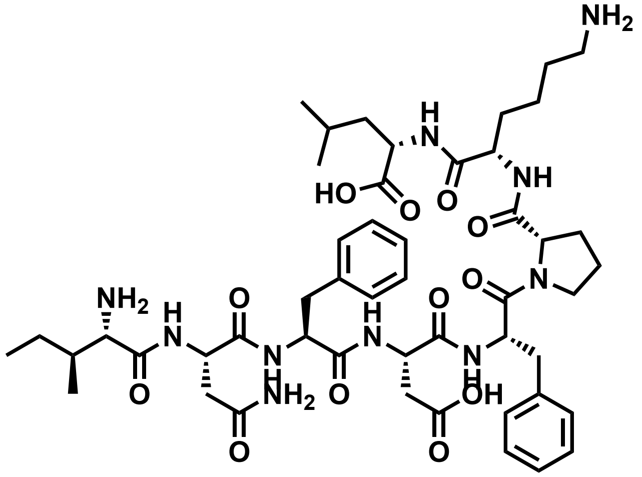

Chemical structure of **ca3**.

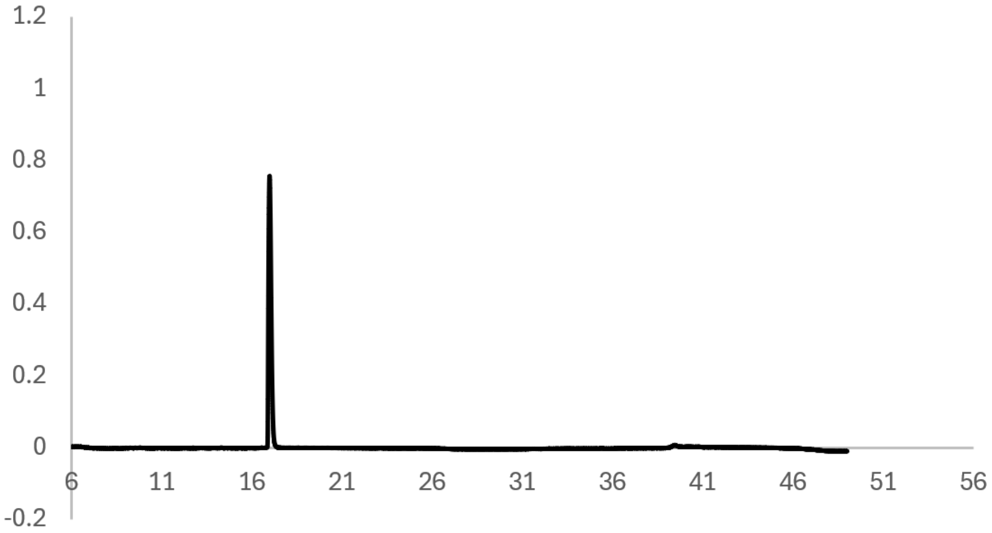


Analytical HPLC Chromatogram of **ca3.**

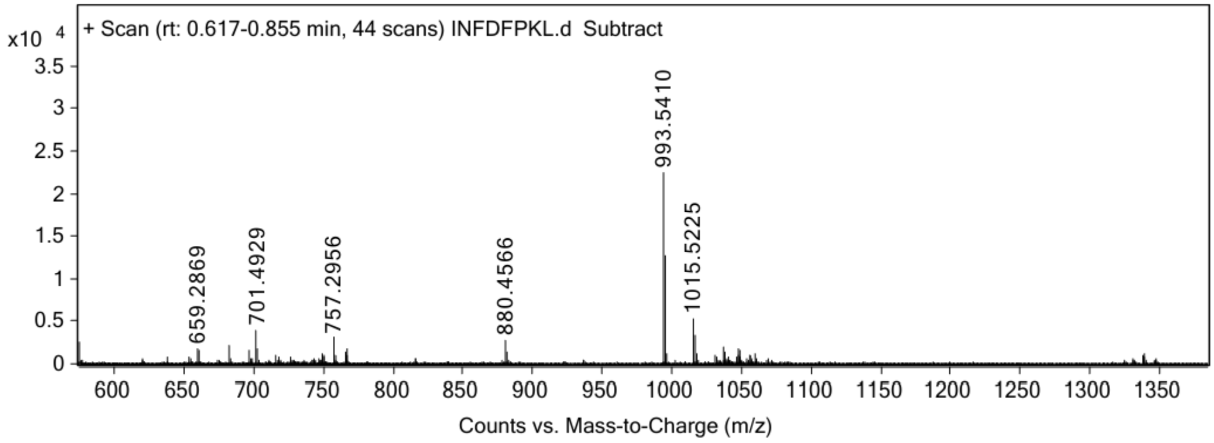


ESI-MS for **ca3** (calculated [M+H$^+$]: 993.5404, found 993.5410).

**ca3$_{asp}$**

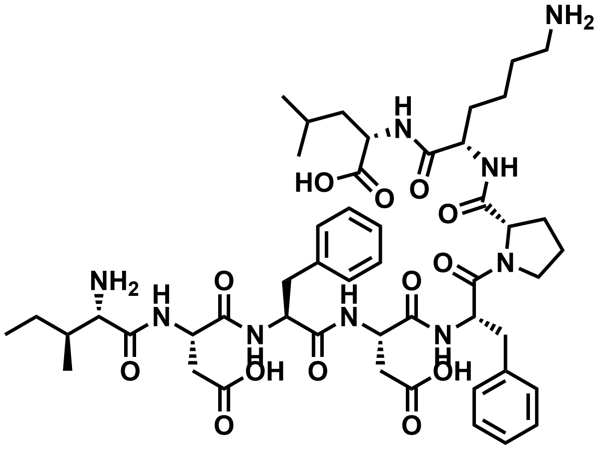

Chemical structure of **ca3$_{asp}$.**

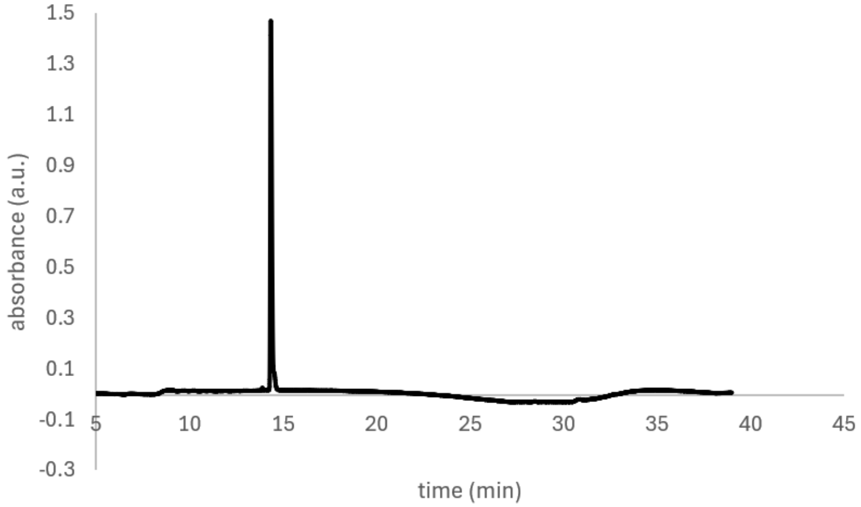


Analytical HPLC Chromatogram of **ca3$_{asp}$.**

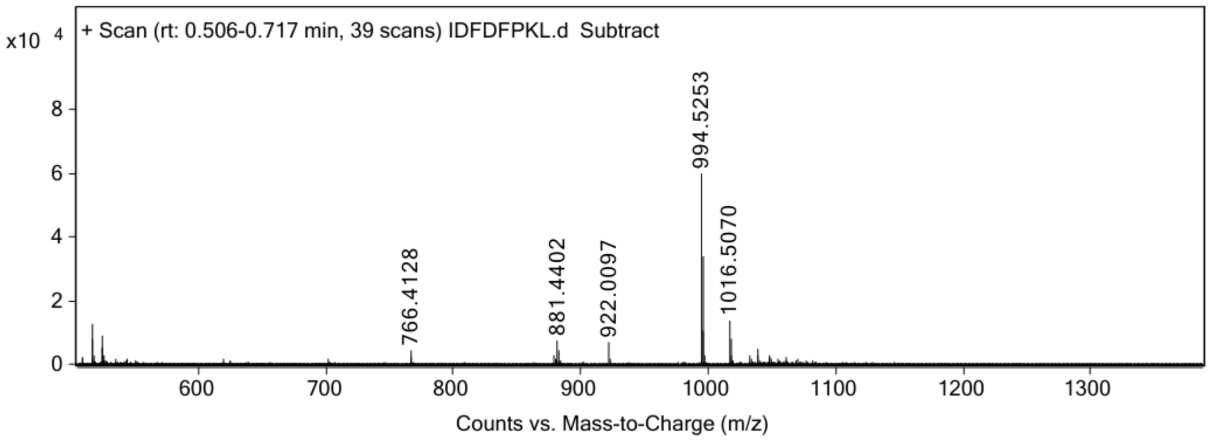


ESI-MS for **ca3$_{asp}$** (calculated [M+H$^+$]: 994.5244, found 994.5253)

**ca3$_{isoasp}$**

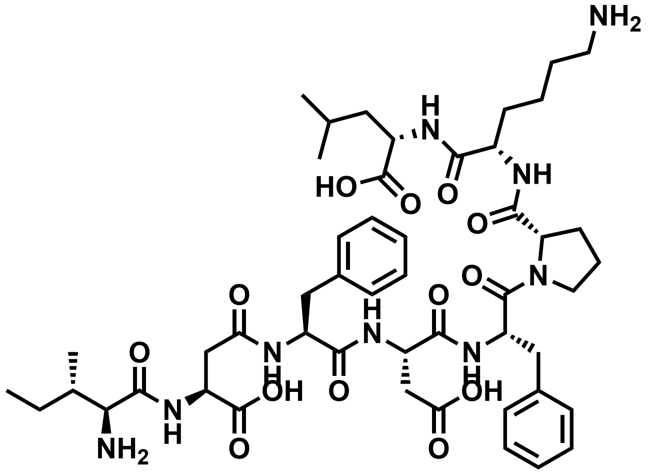

Chemical structure of **ca3$_{isoasp}$.**

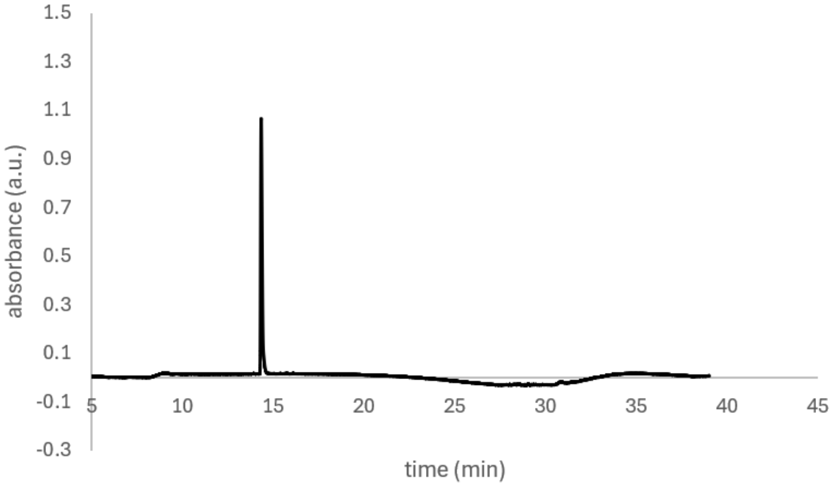


Analytical HPLC Chromatogram of **ca3$_{isoasp}$.**

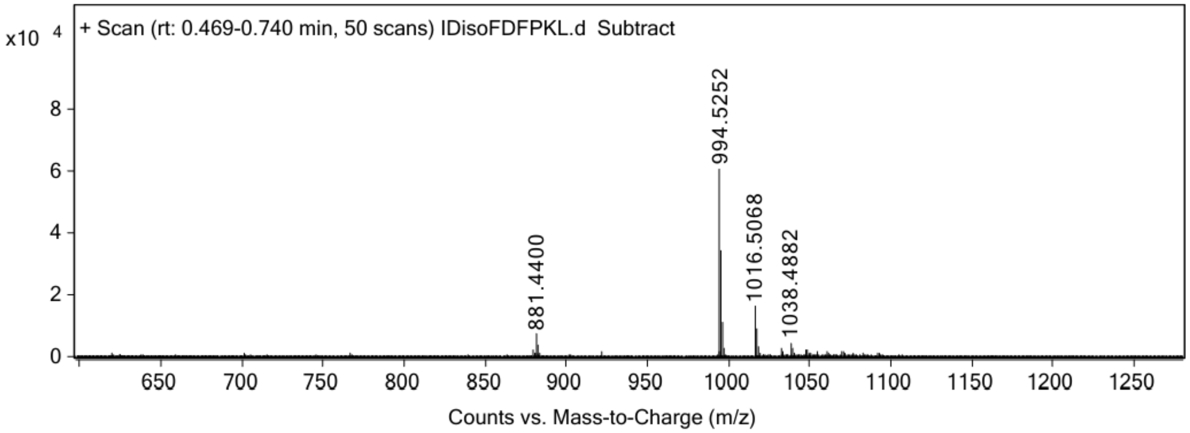


ESI-MS for **ca3$_{isoasp}$** (calculated [M+H$^+$]: 994.5244, found 994.5252).

**ca4 (CGYEFTSKL)**

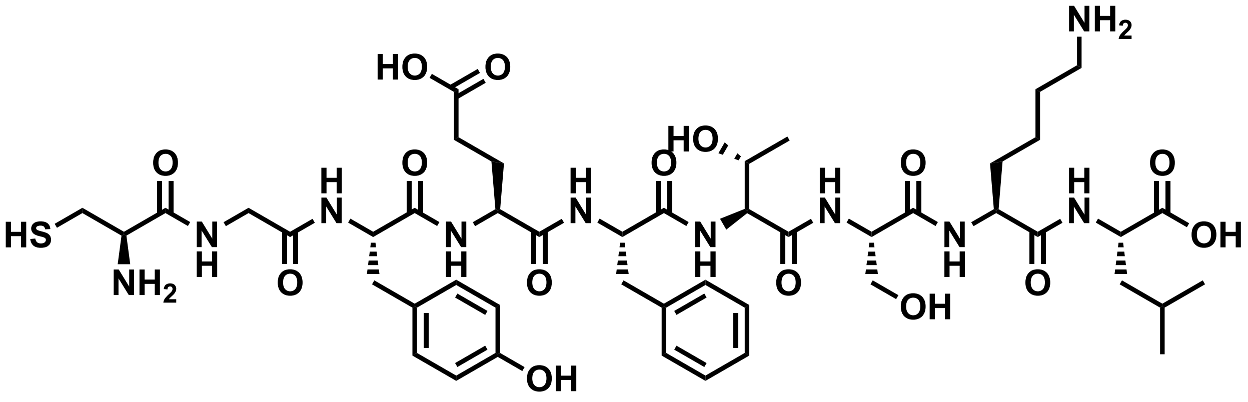

Chemical structure of **ca4**.

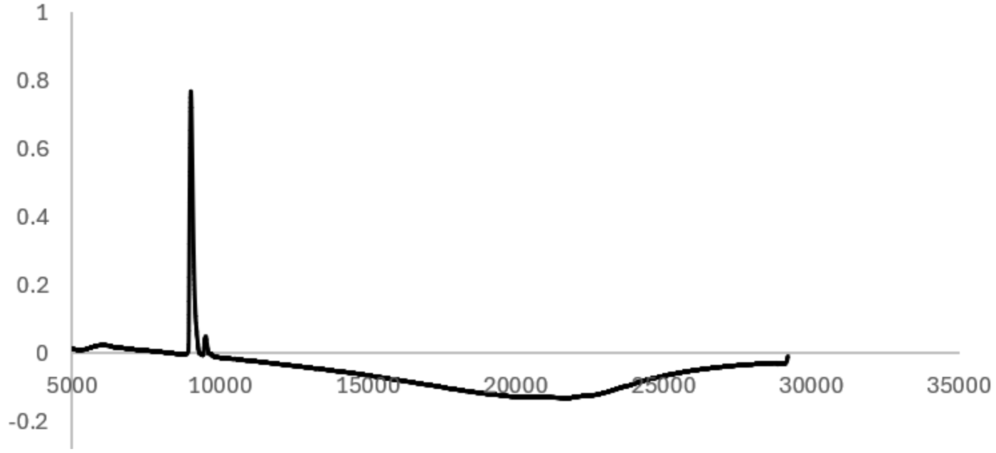


Analytical HPLC Chromatogram of **ca4**.

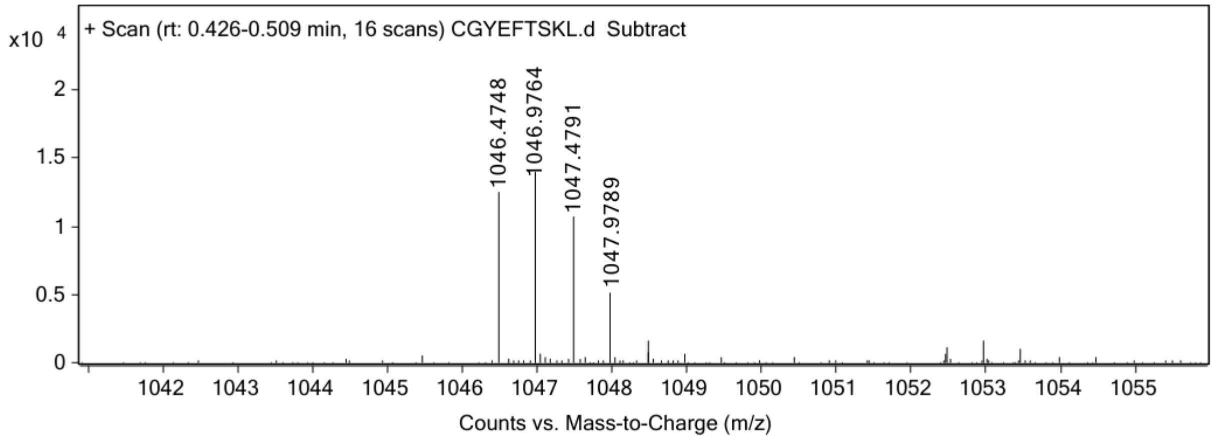


ESI-MS for **ca4** (calculated [M+H$^+$]: 1047.4816, found 1047.4791).

**ca4$_{cystine}$**

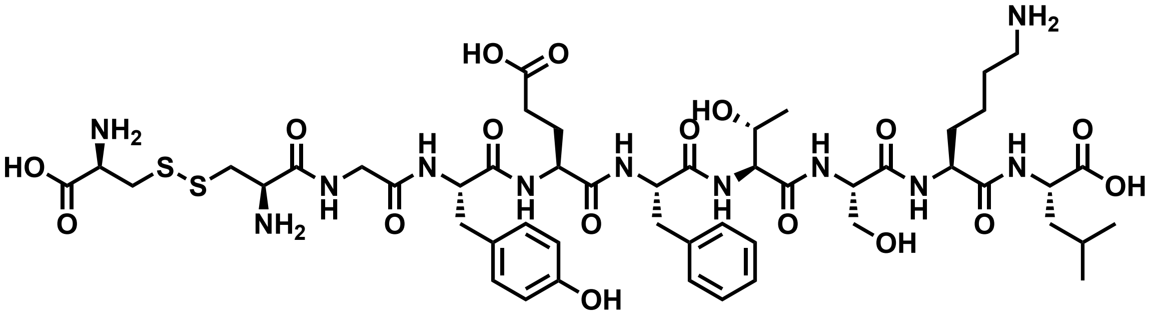


Chemical structure of **ca4$_{cystine}$**.

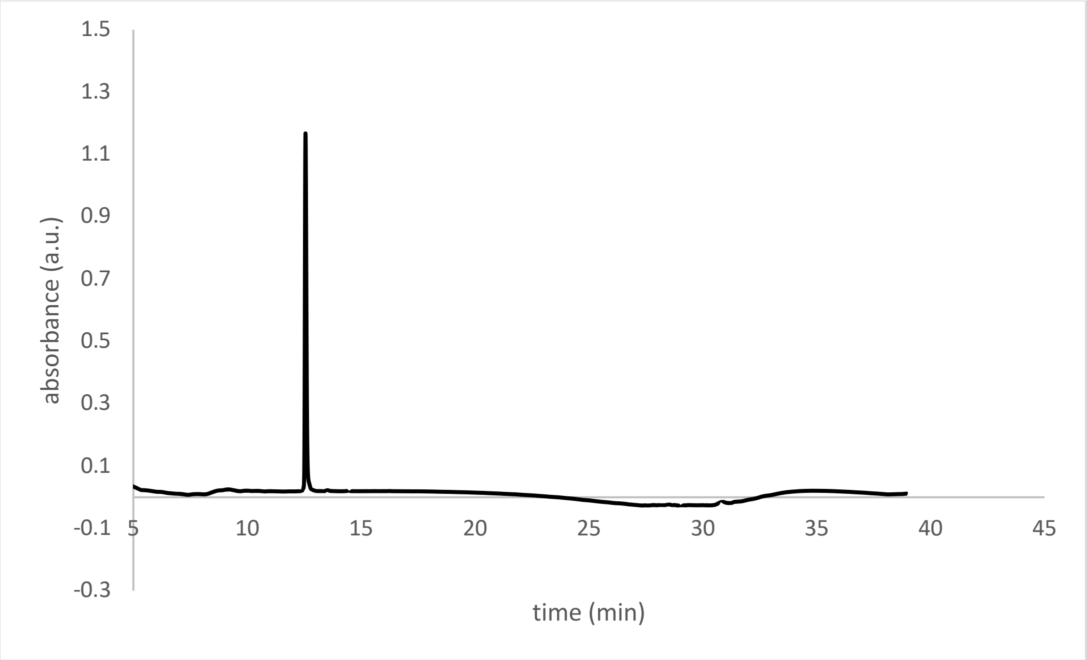


Analytical HPLC Chromatogram of **ca4$_{cystine}$**.

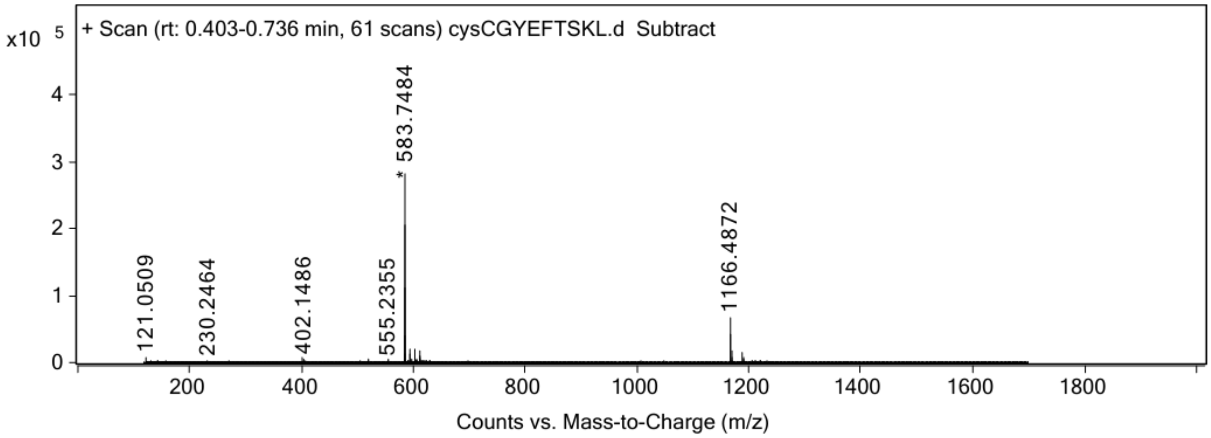


ESI-MS for **ca4$_{cystine}$** (calculated [M+H$^+$]: 1166.4857, found 1166.4872).

**ca5 (VAYEYLCHL)**

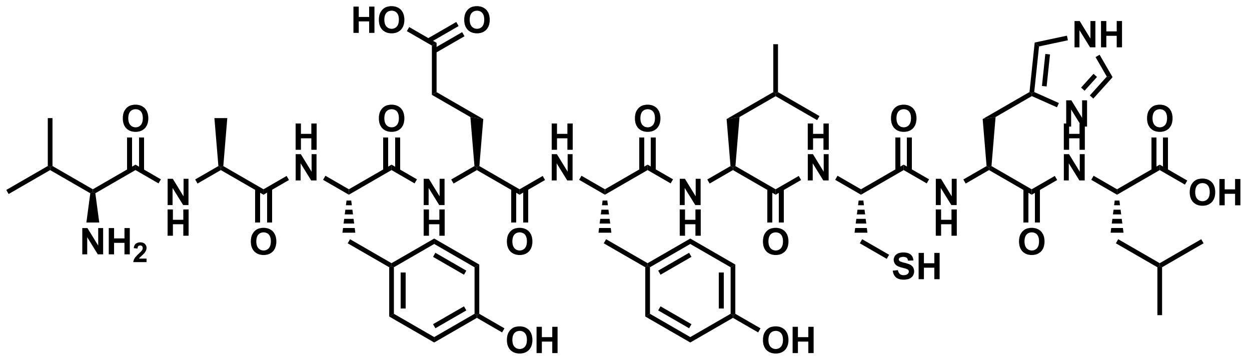


Chemical structure of **ca5.**

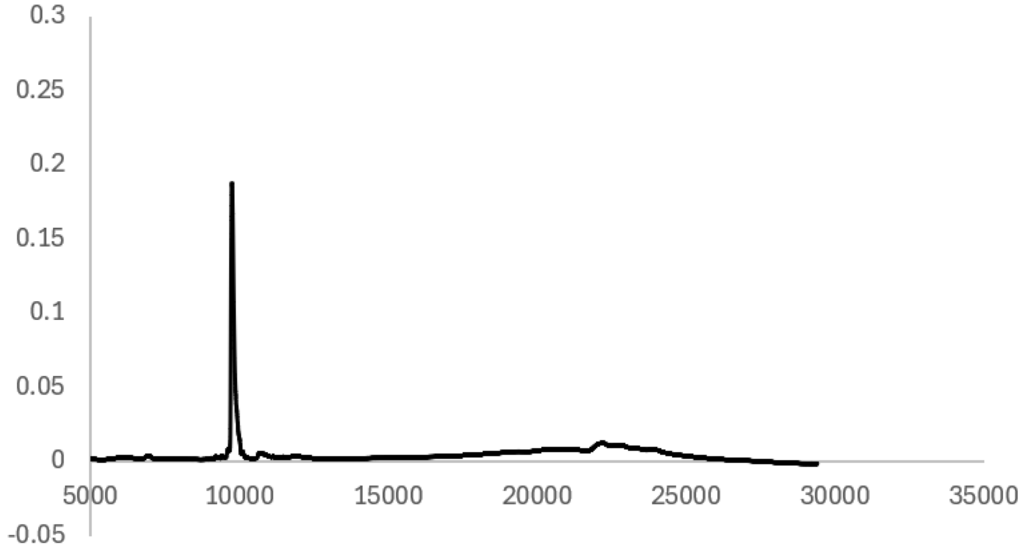


Analytical HPLC Chromatogram of **ca5**.

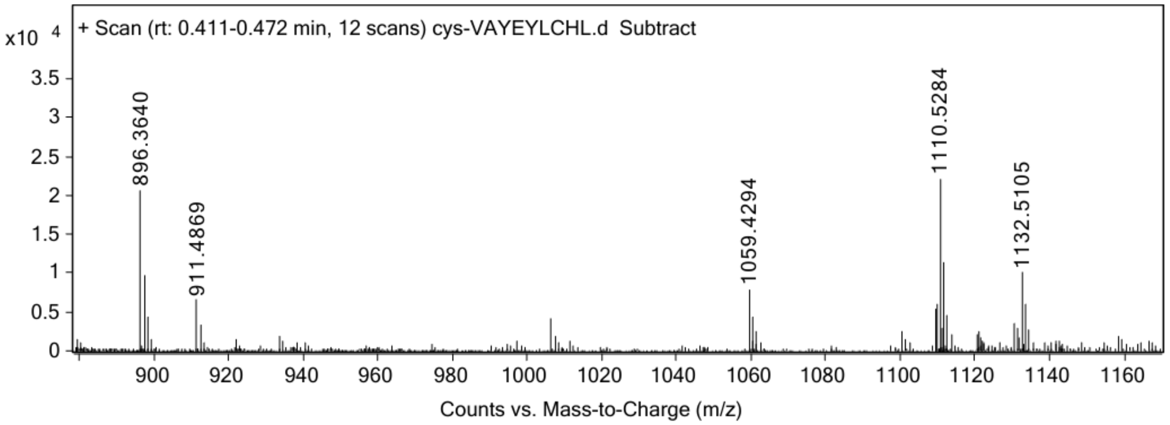


ESI-MS for **ca5** (calculated [M+H$^+$]: 1110.5288, found 1110.5284).

**ca5$_{cystine}$**

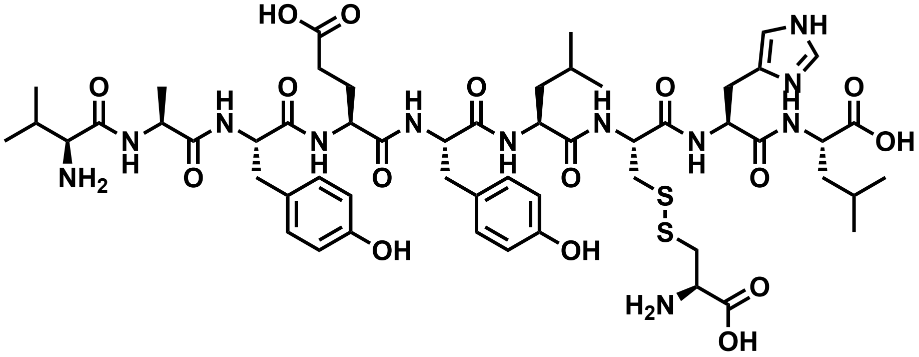


Chemical structure of **ca5$_{cystine}$**.

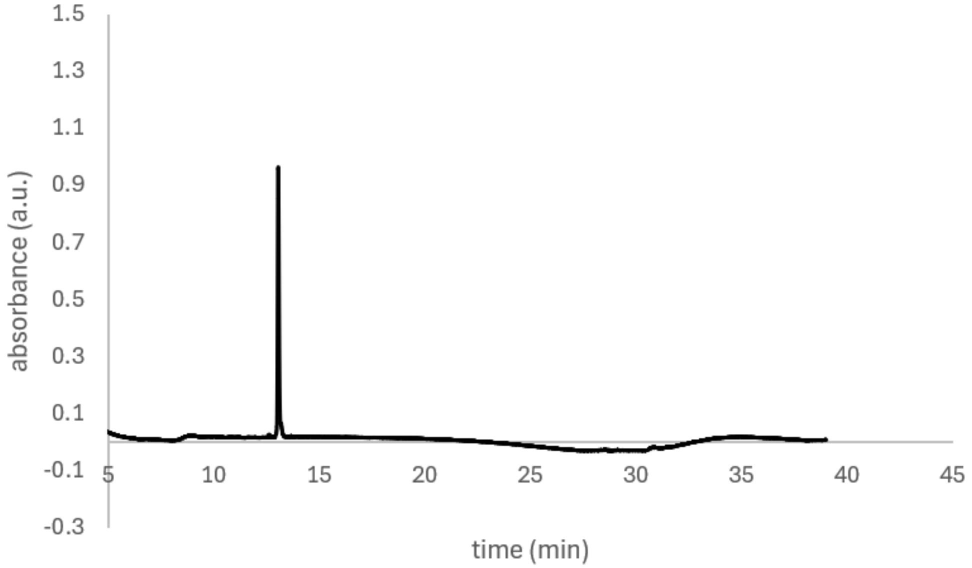


Analytical HPLC Chromatogram of **ca5$_{cystine}$.**

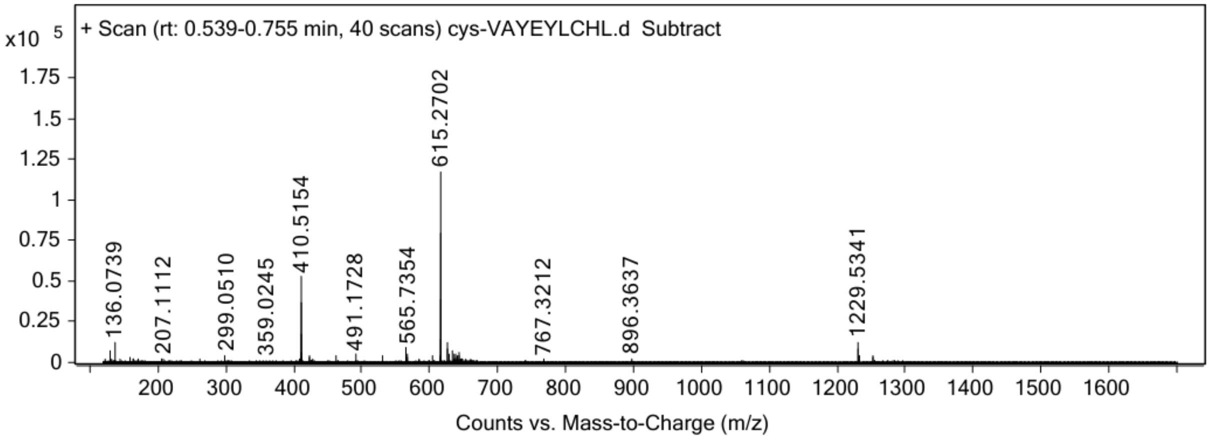


ESI-MS for **ca5$_{cystine}$** (calculated [M+H$^+$]: 1229.5329, found 1229.5341).
